# Deciphering the evolutionary origin of the enantioselectivity of short-chain dehydrogenases from plants toward 1-borneol

**DOI:** 10.1101/2025.07.17.664155

**Authors:** Jasmin Zuson, Carl P.O. Helmer, Bruno Di Geronimo, Andrea M. Chánique, Katarína Kavčiaková, Rosa Teijeiro-Juiz, Sebastian Brickel, Nuria Ramirez Molina, Ivana Drienovská, Daniel Kracher, Eric A. Gaucher, Shina Caroline Lynn Kamerlin, Bernhard Loll, Robert Kourist

## Abstract

Enzyme engineering has produced numerous methods to optimize enzymes for biotechnological processes; however, less is known about how natural evolution creates new functionalities. We investigate the evolutionary emergence of enantioselectivity in plant borneol dehydrogenases (BDHs), which feature hydrophobic active-sites and are enantioselective towards dibornane-type monoterpenols. Ancestral sequence reconstruction provided a trajectory from the oldest unselective BDH ancestor N30 (E=12) toward a more recent selective ancestor N32, involving 19 mutations: 18 mutations are peripheral, one (I111L) occurs in the active-site. The mutation L111I in the hydrophobic pocket increased the selectivity of N30, while the back-mutation I111L decreased the selectivity of N32. Additional peripheral mutations (V136L/G169A/V183I) were required for high selectivity. Crystal structures suggested that protein dynamics, rather than structural changes shape these catalytic properties; this was confirmed by ML/MM simulations of ligand binding. Funnel-metadynamics simulations revealed a correlation between the active-site’s solvent-accessible surface area (SASA) and selectivity. This potential evolutionary pathway shapes enantioselectivity, and guides future enzyme engineering campaigns.

## Introduction

Stereoselectivity enables the striking enantiomerical purity of Nature’s molecules.^1^ The often high enantioselectivity of enzymes towards natural and unnatural substrates is a reason for their broad application in chemical and pharmaceutical industries.^2^ Despite this importance, the evolutionary pathways leading to the formation of stereoselectivity in enzymes remain largely unknown. Enzyme engineering often aims to improve enzymes for the conversion of unnatural substrates, but these engineering campaigns rarely follow the principles observed in natural evolutionary trajectories leading to stereoselectivity. For practical reasons, rational design and saturation mutagenesis focus mostly on the active-site, while the periphery is largely ignored. In contrast, gene randomization methods, such as error-prone PCR, overrepresent peripheral amino acids.^3^ Deep mutational scanning (DMS) can offer insights on structure-function relationships, and, to our knowledge, is starting to be applied in investigation of the stereoselectivity of enzymes.^4,5^

Borneol-type dehydrogenases (BDHs) encompass both highly selective and unselective enzymes towards their natural, chiral substrate 1-borneol. This makes them an interesting model for the study of the natural emergence of stereoselectivity. BDHs are produced during the maturation of the plant, and catalyze the enzymatic production of camphor in oil glands on the leaf surface.^6^ Most known bornyl-diphosphate synthases produce (+)-bornyl-pyrophosphate, which is then hydrolysed to (+)-1-borneol.^7^ Bornyl-diphosphate synthases leading to the (−)-stereoisomer are rare, with only one enzyme described from the plant Sambong.^8^ Stereoselective 1-borneol oxidation does not offer any obvious evolutionary advantages in most plants. This is reflected by the very low stereoselectivity of most characterized enzymes.^9–14^ In the 1980s, Croteau *et al.* reported a stereoselective dehydrogenase from the *Salvia* genus where the essential oil contains (−)-1-borneol and (+)-camphor.^15,16^ The presence of (−)-1-borneol in the essential oil is an indicator for the high (+)- selectivity of its BDH. By mining the genomes of *S. officinalis* and *S. rosmarinus*, we identified several highly stereoselective short-chain dehydrogenases (SDRs) and determined their structures.^17–19^ BDHs have been applied for the kinetic resolution of 1-borneol and 1-isoborneol, which can be derived as racemate from α-pinene, allowing the synthesis of optically pure ingredients of therapeutics, cosmetics, and fragrances from an inexpensive bio-based starting material (**Fig. 1**).^20^

**Fig. 1.**
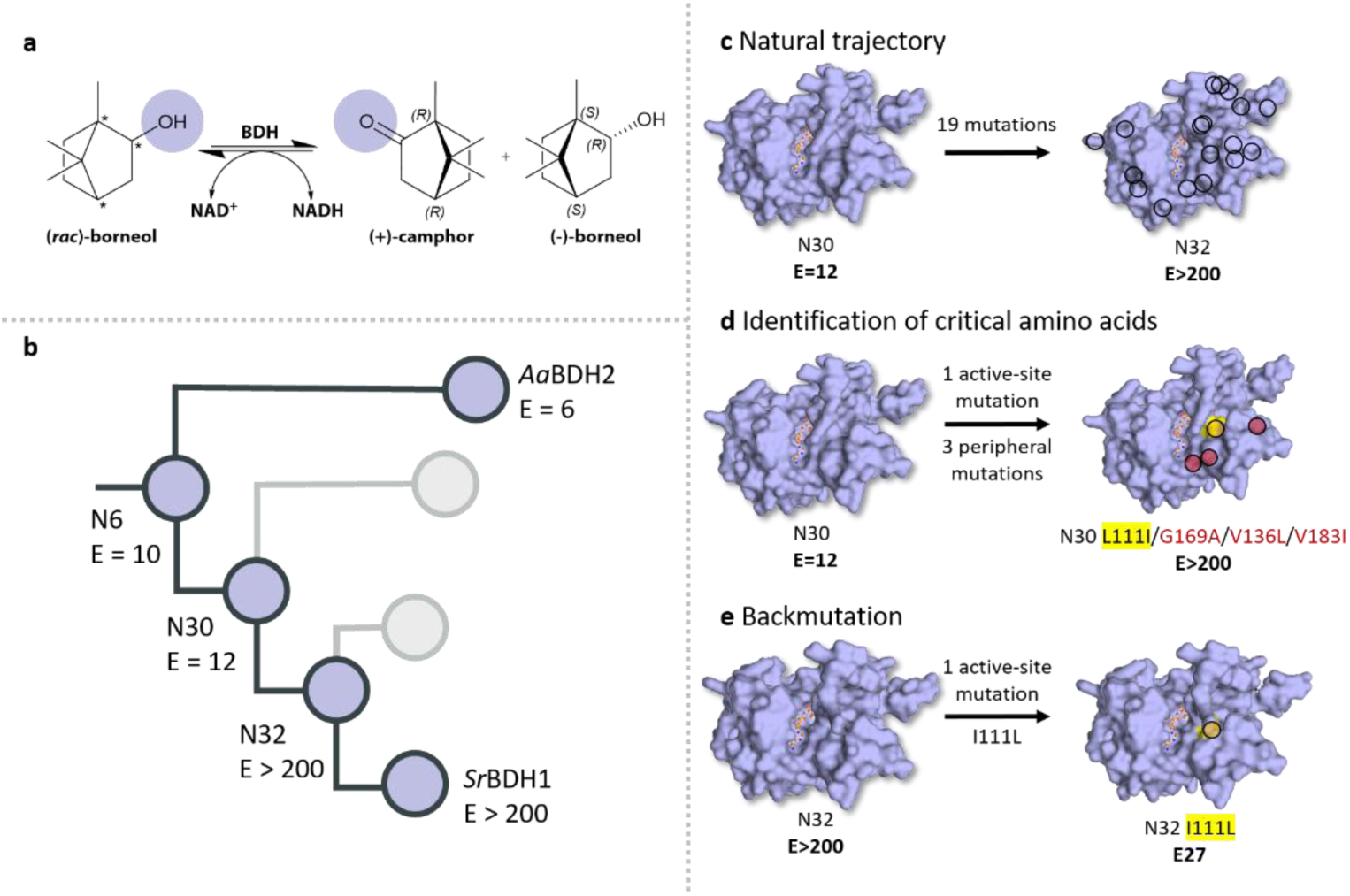
Overview of the evolutionary trajectories. (A) Enantioselective oxidation of 1-borneol by BDHs. (B). Simplification of the evolutionary trajectories leading to the enantioselective *Sr*BDH1. (C) The natural trajectory between N30 and N32. (D) 4 out of 19 mutations between N30 and N32 suffice to make N30 highly selective (E>200). The amino acids in the active-site (yellow) and second shell (red) lead to enantioselectivity. (E) Mutation of the active-site residue (yellow) in the selective ancestor N32 leading to a loss in selectivity.

The structure of *Sr*BDH1 (PDB: 6ZYZ)^19^ shows that the substrate-binding pocket is notably hydrophobic. The 1-borneol molecule is highly compact and has few rotational degrees of freedom. Only the alcohol function of the hydrophobic substrate can engage in hydrogen bonds with the active-site amino acids. To initiate oxidation, the OH-group forms a hydrogen bond with the hydroxyl-tyrosinate ion, and the α-CH must be positioned close to the nicotinamide cofactor.^21^ These two interactions determine productive binding poses. Therefore, discrimination between the two enantiomers must be accomplished by their interactions with the residues lining the hydrophobic active-site pocket. The composition of the hydrophobic pockets of the unselective and selective BDH is very similar. Several point mutations in the first shell of the active-site of the highly selective *Sr*BDH1 did not reduce the high selectivity towards (+)-1-borneol.^19^ This poses a significant challenge to understanding the molecular basis underlying the remarkable selectivity of these enzymes.

Hydrophobic pockets are often crucial elements of enzyme catalysis. For example, unfavourable interactions of a carboxylic group of substituted malonic acids in a hydrophobic pocket of bacterial arylmalonate decarboxylase induce carboxylate cleavage.^22^ Increasing the hydrophobicity of the surface residues by iterative saturation mutagenesis significantly enhanced activity.^23^ Placing a Cys-His-Glu catalytic triad in a hydrophobic lumen of a *de novo* generated catalytic scaffold allowed the creation of an artificial hydrolase.^24^ Baker and Green created a tuneable hydrophobic pocket within a designed heme-binding protein that catalyzed cyclopropanation reactions.^25^ Hydrophobic pockets are also crucial for the stereoselectivity of enzymes. For instance, the different accommodation of isomers in hydrophobic pockets is decisive for the stereoselectivity of α/β-fold hydrolases, and saturation mutagenesis of amino acids in the hydrophobic surface increased stereoselectivity.^1,26–30^

In these examples, improvements of the enzymes were achieved by combinatorial methods. These have in common that precise predictions on the outcome of amino acid substitutions within hydrophobic pockets are difficult with the currently available knowledge and methodology. A comparison of unselective and selective extant enzymes may reveal clues to the difference in selectivity. In horizontal approaches, extant enzymes are directly compared with each other to identify potential target residues.^31,32^ While this method compares individual residues, it neglects any context-dependent changes that the enzymes have accumulated during their evolutionary history. Without context, transplantation of potential selectivity-determining residues is likely to have detrimental effects on activity. In contrast, vertical approaches use ancestral sequence reconstruction (ASR) to create evolutionary information, which was gradually lost over time.^31–33^ Narayan *et al*. resurrected flavin-dependent monooxygenases and demonstrated that exchanging a small number of active-site residues could switch the stereochemical outcome of oxidative dearomatization.^31^ Similarly, a study on imine reductases revealed the residues involved in stereoselectivity, allowing for inversion of the reaction.^34^ A campaign on vanillyl alcohol oxidases successfully identified key residues for broadening the substrate scope of these enzymes.^35^ A study on spiroviolene synthases created ancestral variants that exhibited higher stability and solubility, enabling the crystallization of an ancestor.^36^ The ancestral proteins further pointed to specific amino acid substitutions, allowing for fine-tuning of the substrate specificity in the extant enzyme.^36^ Furthermore, evolutionary information can guide through fitness landscapes.^37^

Comparison of the last unselective ancestor with more recent, selective ancestors may reveal evolutionary ‘switches’, i.e. residues and mutations involved in the emergence of stereoselectivity in the BDH enzyme family. Herein, we use ancestral sequence reconstruction, structural analysis, site-directed mutagenesis and computational simulations to demonstrate how epistatic effects determine the evolutionary emergence of stereoselectivity in enzyme catalysis.

## Results and discussion

### Ancestral sequence reconstruction of BDHs

The sequences of ten known plant BDHs were used as queries for a search against the 1KP database^38,39^. We parsed out the data to make sure that all BDH sequences selected possessed the conserved catalytic residues (SxxxxxxxxxxxxYxxxK) **(Supplementary Fig. 1**) and the cofactor binding motif (TGxxx[AG]xG) (**Supplementary Fig. 1)**. With that, ninety-seven sequences that belong to 59 plant genera were selected to construct a phylogenetic tree **(Fig. 2A)**. Among these ninety-seven sequences, three unselective enzymes (*Aa*ADH2, *Aa*BDH2 and *Ps*BDH) and four selective borneol dehydrogenases (*Sr*BDH1, *Sr*BDH2, *So*BDH1 and *So*BDH2) have been biochemically characterized^12,19,40–42^. These three unselective enzymes are present in species that are within the plant kingdom and are, as such, evolutionarily distant to the selective borneol dehydrogenases. Ten ancestral proteins from different nodes of the tree (N4, N5, N6, N7, N30, N32, N39, N48, N49, and N74) were selected for biochemical analysis.

**Fig. 2.**
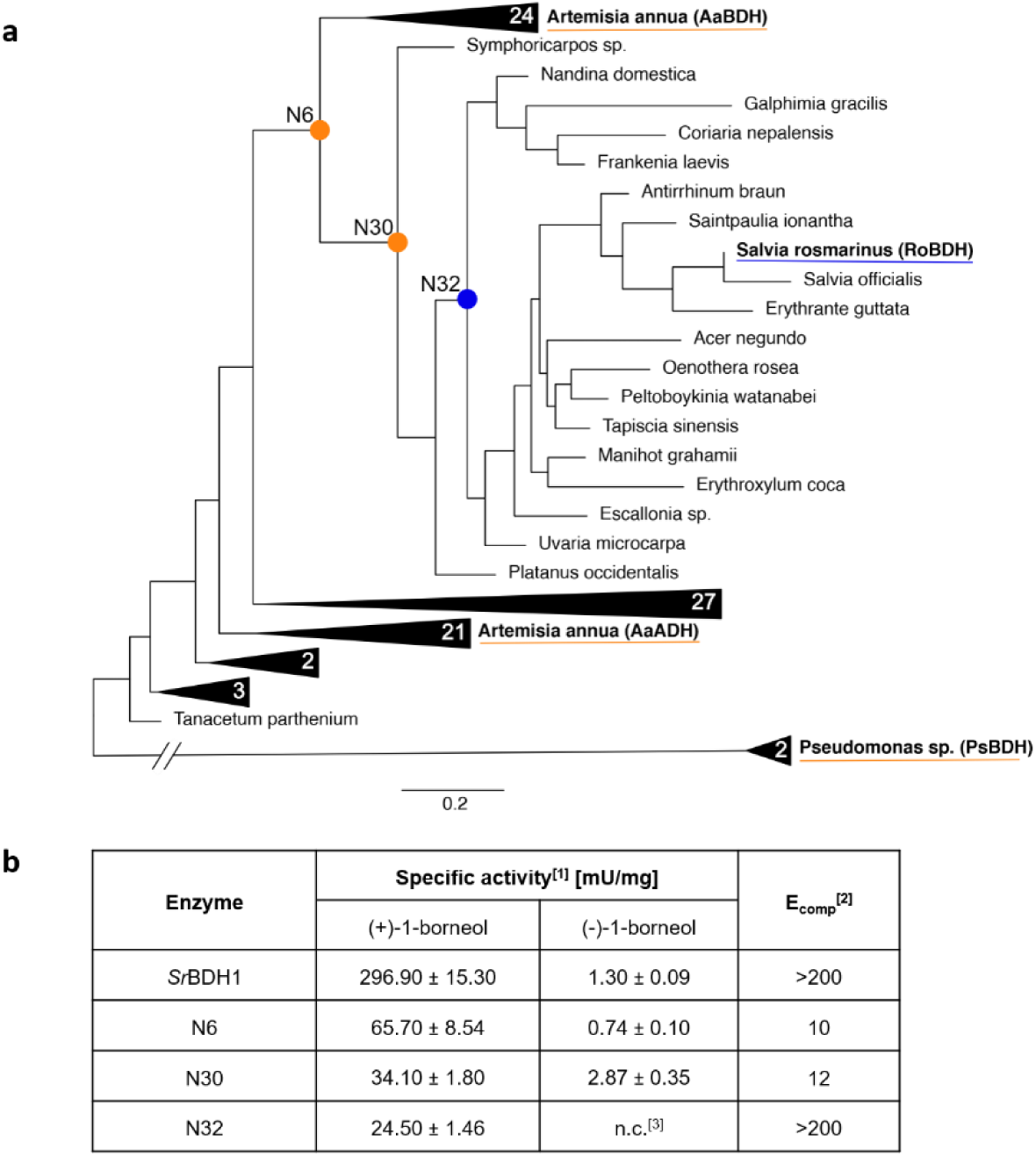
(A) Phylogenetic tree of plant BDHs. Circles at nodes represent ancestral proteins that highlight the transition from unselective to selective enzymes. Unselective BDHs are represented in orange, while selective BDHs are dark blue. The number inside the triangles represent the total number of sequences in that specific node. The aLRT of nodes N6, N30, and N32 as calculated by PhyML was 0.95, 0.94, and 0.81, respectively. The posterior probability of nodes N6, N30, and N32 are calculated by GRASP as 0.96, 0.94, and 0.88, respectively. Scale bar represents 0.2 amino acid replacements per site per unit evolutionary time (branch leading to the outgroup is truncated). (B) Specific activity and enantioselectivity E_comp_-values determined for *Sr*BDH1 and ancestors N6, N30 and N32 by oxidation of (+/−)-1-borneol enantiomers in comparison to the competitive oxidation between (+/−)-1-borneol and the reference compound. ^[1]^ determined by GC; ^[2]^ determined as the ratio of specific activities of an enantiomer over the rate of a competing co-substrate divided by the rate of the other enantiomer over the competing co-substrate (**Supplementary Equation S1**); ^[3]^ n.c.: no conversion.

Most ancestral variants could be produced as soluble proteins in *Escherichia coli* (**Supplementary Table 1**), except N48 that precipitated during purification, similar to several extant BDH,^18^ which prevented their further characterization.

### Determination of the enantioselectivity of ancestral proteins

The ancestors closest to the last common ancestor (N4, N5, N6) displayed the highest activity in the oxidation of (−)-1-borneol, the slower-reacting enantiomer for all characterized BDH (**Supplementary Fig. 3)**. N32, is the first ancestor showing high selectivity (E>200) in the evolutionary trajectory from N6 to *Sr*BDH1. We assume that the evolutionary ‘switch’, *i.e.* the mutations responsible for the high selectivity of N32 and *Sr*BDH1, lie within the sequence space between N6 and N32. Therefore, ancestors N6, N30, and N32 were chosen for further characterization.

We determined the Michaelis-Menten kinetics of the three ancestors towards the faster-converted (+)-1-borneol, which was complicated by the low solubility of the substrate in aqueous solution^43^ and required extrapolation of data obtained below the *K*_M_-value. Addition of 20% DMSO enhanced the solubility of borneol from 4.8 to 9.0 mM, while retaining around 40% of the overall enzymatic activity (**Supplementary Fig. 4**). Nevertheless, substrate saturation could not be reached, thus values are reported as apparent *K*_M_-values (*K*_app_). Ancestor N6 exhibited substrate inhibition with (+)-1-borneol (*K*_i(+)-bor_ [mM] = 2.5). Surprisingly, the *K*_app_-values for the ancestors N30 (*K*_app_ = 20.27±7.21 mM) and N32 (*K*_app_ = 17.36±5.40 mM) were ten-fold higher than those of the wild-type *Sr*BDH1 (*K*_app_ = 2.02±0.18 mM)^19^, while the *K*_app_ of N6 (*K*_app_ = 0.01±0.01 mM) was below that of the wild-type (**Supplementary Fig. 5**, **Supplementary Table 2**).

The strong substrate inhibition of ancestor N6 rendered measurement of the E-value from enantiomeric excess and conversion unsuitable.^44,45^ Furthermore, the low solubility of 1-borneol prevented measuring catalytic efficiencies for the determination of the E-value. The Quick-E method developed by Kazlauskas *et al*. provided a suitable method to include the competition of both enantiomers.^45,46^ In this approach, the activity of an enantiopure substrate is measured against a non-chiral surrogate competitor and is correlated to the reaction of the opposing enantiomer. This setup mimics the natural competition between two enantiomers. Oxidation of the slower-converted enantiomer (−)-1-borneol with and without surrogate substrate (**Supplementary Fig. 6**) confirmed that the competition with the surrogate substrate decreased the activity towards (−)-1-borneol, thus mimicking a racemic reaction. Determination of E_comp_-values (**Fig. 2**) showed that under competition, ancestors N6 and N30 have low selectivity towards the 1-borneol enantiomers. N30 lost the substrate inhibition observed in N6, and N32 is the first ancestor to be specific for (+)-1-borneol. While having similar total activity toward (+)-1-borneol compared to the N30, no activity toward (−)-1-borneol was detectable. N32 is highly selective under competitive conditions (E>200), highlighting the transition from N30 to N32 as the critical evolutionary step towards enantioselectivity.

### Structure determination of ancestral proteins

BDHs can exhibit two distinct conformations.^47^ Upon substrate binding, the enzyme transforms from an ‘open’ to a ‘closed’ conformation by the movement of the αFG-helix and loop toward the active-site (**Fig. 3**).^47–50^

**Fig. 3.**
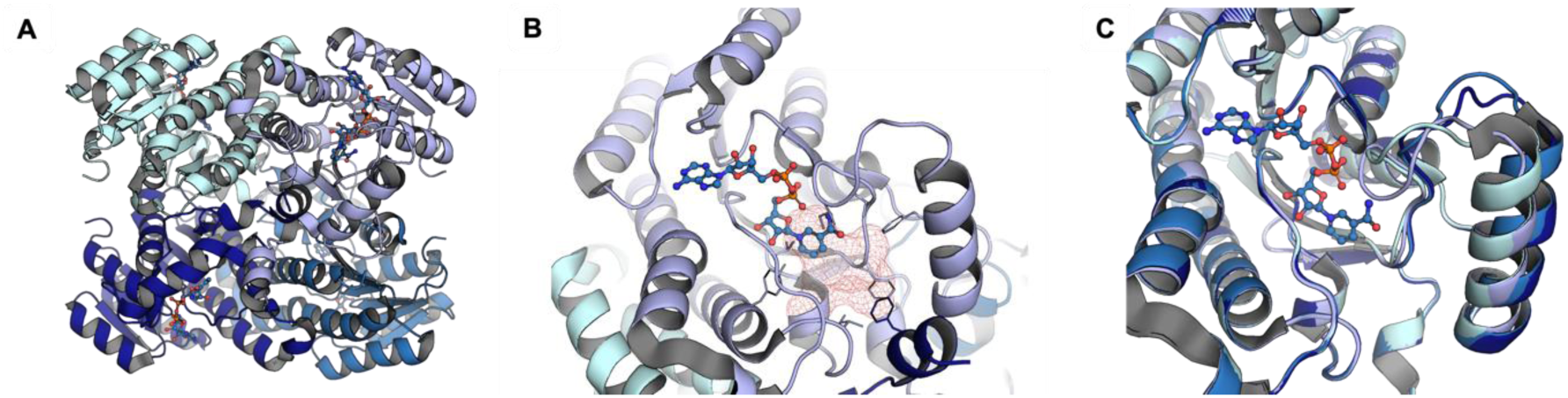
(A) Structural overview of *Sr*BDH1 based on PDB: 6ZYZ^19^ showing its tetrameric composition. (B) The active-site of *Sr*BDH1 with NAD^+^ depicted in stick representation. The active-site pocket is highlighted by a red mesh. (C) Superimposition of the four chains from PDB: 6ZYZ^19^, highlighting the αFG-helix and the preceding loop conformations among them.

To account for these conformational dynamics, we attempted to obtain crystal structures of the ancestors N6, N30, N32 and N39, but were only able to collect high-resolution diffraction data of the tetrameric N32 and N39. N32 is arranged as a tetramer in the unit cell (**Supplementary Tables 3-4**, **Supplementary Fig. 7**). As there is a single N39 protomer in the asymmetric unit, the tetrameric arrangement unit was calculated based on crystal symmetry. Both structures superimpose with an RMSD of 1.19 Å (**Supplementary Table 5**). The overall structure of N32 resembles the structures of *Sr*BDH1 as well as *So*BDH2 (**Fig. 4**). Co- crystallization and soaking experiments of the protein crystals with either (+)-1-borneol or (−)-camphor did not lead to structures bound to these compounds.

**Fig. 4.**
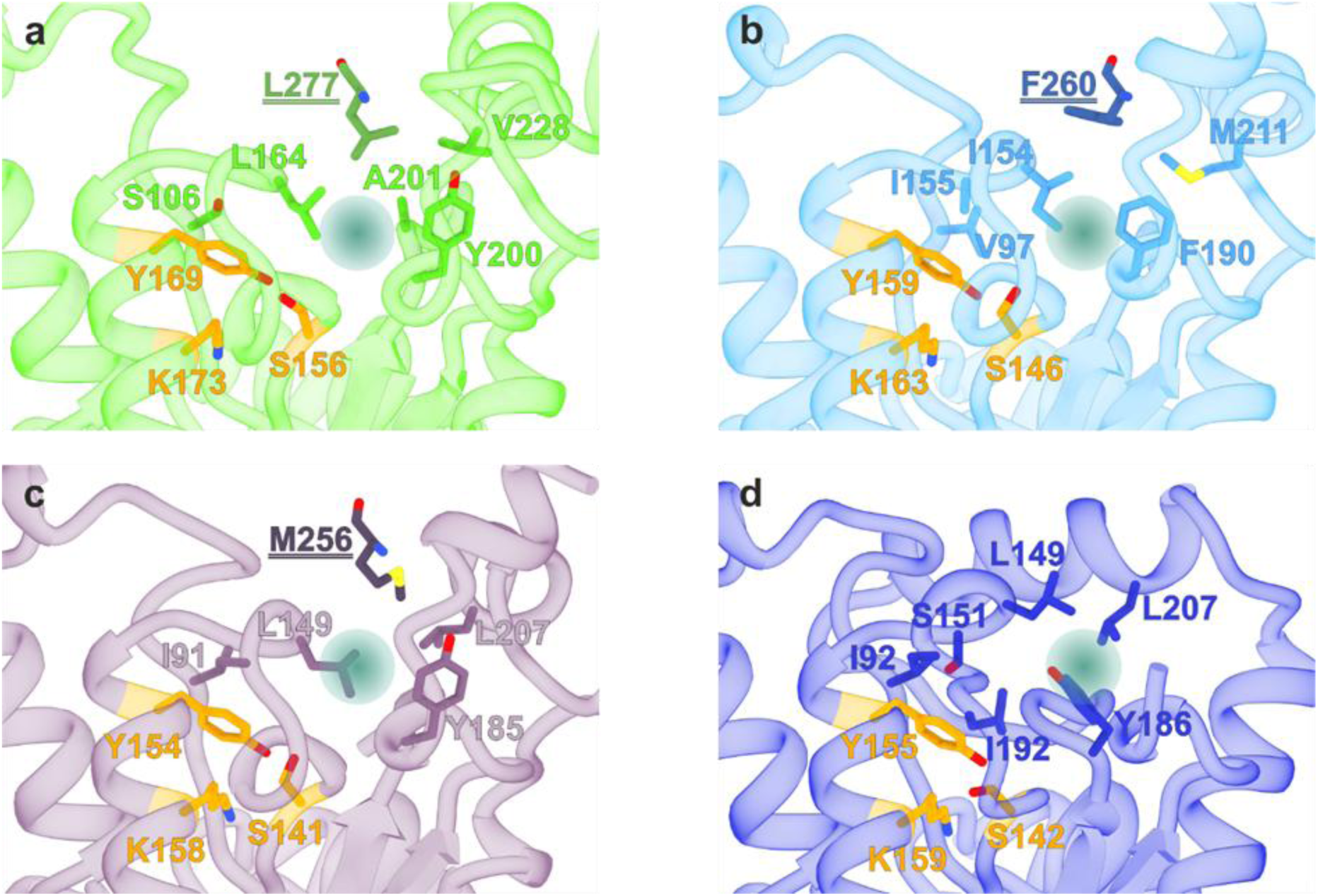
Comparison of the active sites of (a) SrBDH1 (PDB: 6ZYZ)^51^, (b) SoBDH2 (PDB: 7O6P^18^), (c) N32, and (d) N39. Catalytic residues are represented in orange. Residues originating from the adjacent protomer within the tetramer are underlined and represented in a darker shade, except for N39, where only chain A was resolved. Most residues lining the 4 active sites are hydrophobic. The teal-colored sphere represents the potential substrate binding-site.

As no high-resolution crystal structure of N6 was obtained, AlphaFold3^52^ was used to generate a model, which showed a mean pLDDT of 89.39 with the lowest scores located at the termini (**Supplementary Fig. 8**). Despite an overall high similarity with *Sr*BDH1, differences can be observed in the loop-αFG motif of chain A, whereas chains C and D have a closed conformation, which aligns with AlphaFold3 models (**Supplementary Fig. 9**). Aligning the model of N6 to the crystal structure of *Sr*BDH1 yields an RMSD of 0.308 (**Supplementary Table 5**). Since the open confirmation appears to be an artefact unique to chain A of PDB: 6ZYZ^51^, the generated models of non-crystallized variants were used for structural analysis.

### Analysis of evolutionary pathways between ancestors

The first shell of the enzyme was defined as the residues within a 6 Å radius of the nicotinamide moiety of the cofactor, while residues outside were considered as secondary shell. In N6, the first shell hosts 16 residues, in N30 15 residues, in N32 17 residues and in *Sr*BDH1 14 residues. N6 and N30 differ by 38 amino acids (three in the first shell), while N30 and N32 differ by 19 amino acids (only one in the first shell). A full combinatorial approach, would result in a sequence space that is too large for a comprehensive kinetic analysis.^53^ Motivated by prior studies indicating that interactions with second-shell residues can affect enzyme selectivity^19^ we incorporated second-shell variants into our investigation.

To probe the three residues differing in the first shell of N6 and N30, we incorporated the first shell of N30 into N6, creating four variants (N6_V167A; N6_A211G; N6_A213V; N6_V167A/A211G/A213V). Point mutations of N6 did not relieve the substrate inhibition. Oxidation of (+)-1-borneol by N6_V167A and N6_A211G was lower at concentrations below and above the *K*_i_ of N6 (**Fig. 5**). In contrast, the substitution A213V lowered the substrate inhibition of N6. Combining the residues in N6_V167A/A211G/A213V lead to an activity similar to that of N6, neutralizing any negative or positive effects. The inhibition was still present when the complete set of first shell mutations of the non-inhibited N30 was introduced to N6, indicating that it is heavily influenced by peripheral residues, independent of or synergistic to the first shell residues.

**Fig. 5.**
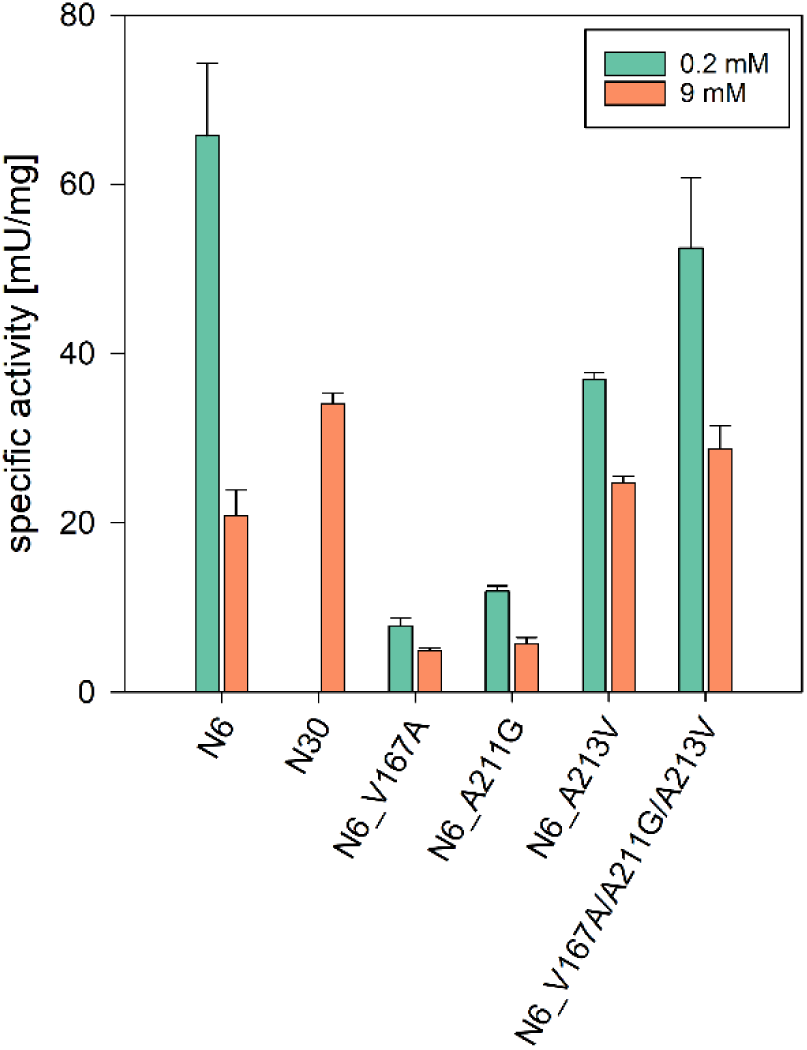
Oxidative rates for (+)-1-borneol given in [mU/mg] catalyzed by the purified mutant variants of N6. Reaction conditions: 2 µM enzyme, 20% DMSO, 25°C, 300 rpm, 1 or 9 mM NAD^+^, 0.2 or 9 mM (+)-1-borneol, 0.1 M Tris-HCl pH 9.0. Measurements were carried out in triplicates.

While selectivity increases slightly between ancestor N6 (E_comp_ = 10) and ancestor N30 (E_comp_ = 12), the major gain of selectivity is achieved in the mutational transition from N30 to N32 (E_comp_ > 200). The loss of the ability to oxidize (−)-1-borneol in the transition between ancestors N30 and N32 highlights key mutations. Interestingly, the transition between N30 and N32 involves a single mutation L111I in the first shell. To test its significance as a potential evolutionary switch, a bidirectional exchange of the first shell residues was performed by site-directed mutagenesis (N32_I111L, N30_L111I). Mutants were purified, rates and enantioselectivity were determined.

N30_L111I (E_comp_ = 21) had a two-fold higher selectivity, while the backmutation of I to L in N32 induced activity in this ancestor towards (−)-1-borneol. N32_I111L is converting both enantiomers (E_comp_ = 27), whereas N32 did not show any activity towards (−)-1-borneol. We note that this mutation in the hydrophobic pocket is crucial for the specificity of N32, underlining that residue 111 is a switch in the evolutionary emergence of selectivity in the BDH.

The underlying ASR data suggest different ancestral protein sequences based on both the marginal and the joint ancestral sequence reconstructions.^54^ In the consensus sequence, the position in question (**Supplementary Fig. 10,** position 158; position 111 in N30/N32) is highly variable, with 11 amino acids present in the multiple sequence alignment (MSA). The most abundant amino acid is S, followed by I, V, P, and A. In the marginal reconstruction, for N30 V (posterior probability (PP)=0.54), I (PP=0.38), or L (PP=0.06) are suggested, while N32 hosts an I at this position. In the joint reconstruction, which was chosen for this study, the inferred residues are N30 L111 and N32 I111.

As the extant *Sr*BDH1 contains a Val in the homologous position 97,^19^ all three proposed mutations are covered by the set of ancestors obtained from the joint reconstruction. Position 111 is located directly downstream of the NNAG motif, another highly conserved motif in SDRs stabilizing the central β-sheet of the Rossman-fold.^21,48,49^ Previous studies on SDRs, including hydroxysteroid dehydrogenases^48^, actinorhodin polyketide ketoreductase^55^, and farnesol dehydrogenases^56^ suggested that in the hydrophobic pocket of SDRs a water relay mechanism involves the N of the NNAG motif. The NNAG motif bridges the bulk water to the catalytic K in the hydrophobic pocket. Local water molecules relay a proton over the NNAG motif to stabilize the positive charge of the catalytic K. It was stated that small residues in the proximity of Asp facilitate easier proton transfer than bulky ones due to their higher rotational degrees of freedom.^48,49^ Extending this hypothesis, the substitutions of the local residue 111 to I/L/V are likely to influence the proton transfer, probably impacting the conversion of the two stereoisomers.

While this residue proved key to the enantioselectivity, the complete suppression of conversion of the slower-reacting enantiomer appeared to depend on additional interactions within the remaining 18 secondary shell residues. Additional peripheral substitutions were required to achieve reversal of selectivity. In addition to the ancestral mutant, *Sr*BDH1_V97L was designed to investigate the effects of exchanging the position homologous to position 111 in N30 on the enantioselective extant enzyme. *Sr*BDH1_V97L showed similar activity toward (+)-1-borneol as N30_L111I but did not convert (−)-1-borneol, confirming the importance of additional peripheral mutations (**Table 1**).

**Table 1.** Specific activity and enantioselectivity E_comp_-values determined for BDH variants by oxidation of (+/−)-1-borneol enantiomers in comparison to the competitive oxidation between (+/−)-1-borneol and the reference compound.

| Enzyme | Specific activity<sup>[1]</sup><br>[mU/mg] | | $E_{\text{comp}}$<sup>[2]</sup> |
| --- | --- | --- | --- |
|  | (+)-1-borneol | (–)-1-borneol |  |
| N32_I111L | 35.9 ± 1.8 | 2.9 ± 0.1 | 27 |
| N30_L111I | 14.2 ± 0.2 | 0.05 ± 0.01 | 21 |
| N30_L111I/M213I/A214G | 12.6 ± 0.9 | 0.4 ± 0.01 | 8.3 |
| N30_L111I/A222V/E223A/S227A/S232V | 62.4 ± 2.9 | 0.5 ± 0.06 | 128 |
| N30_L111I/S166A/V167S/G169A | 271.3 ± 9.7 | 0.4 ± 0.2 | 31 |
| N30_L111I/V136L/S139G/V183I | 174.2 ± 9.7 | 0.4 ± 0.02 | 25 |
| N30_L111I/L47W/E74K/V79I/E153Q/Y197H | 9.1 ± 0.2 | n.c. | >200 |
| N30_L111I/V136L/G169A/V183I | 9.9 ± 1.7 | n.c. | >200 |
| *SrBDH1* V97I<sup>[5]</sup> | 10.7 ± 2.1 | n.c. | >200 |
| N32_I111V | 5.8 ± 0.7 | n.c. | >200 |
| SrBDH1 | 296.9 ± 15.3 | 1.3 ± 0.09 | >200 |
| <sup>[1]</sup> determined by gas chromatography; <sup>[2]</sup> determined as the ratio of specific activities of an enantiomer over the rate of a competing co-substrate divided by the rate of the other enantiomer over the competing co-substrate (**Supplemental Equation S1**); <sup>[4]</sup> n.c.: no conversion; <sup>[5]</sup> V97 in SrBDH1 is equal to Ile111 and Leu111 in N30 and N32. |  |  |  |

Rational mutational approaches were chosen to uncover epistatic effects of position 111 with further peripheral amino acids in N32. The variant N30_L111I was chosen as base for the introduction of further co-mutations, to construct an enantiospecific variant. For this, the remaining 18 amino acids between N30 and N32 were targeted. Targeted mutations were based on structure-function relations in SDRs (N30_L111I/M213I/A214G)^21,49^, the Grantham’s distance matrix N32/N30 (N30_L111I/S232V)^57^, mutational hotspots (N30_L111I/A222V/E223A/S227A/S232V), evolutionary coupling analysis (N30_L111I/S166A/V167S/G169A)^58^, and conserved residues (N30_L111I/V136L/G169A/V183I). Mutations in N30_L111I/M213I/A214G are located in the vicinity of the loop region between the βF-sheet and αG helix of N30_L111I. Residue S232 shows the highest Grantham’s distance, which is located in the αG helix. The αG helix is a hotspot of mutations, comprising four substitutions of N30_L111I/A222V/E223A/S227A/S232V within a distance of ten amino acids. Evolutionary coupling analysis highlighted three residues that possibly co-evolved with position 111 (**Supplementary Fig. 11**), mirrored in N30_L111I/S166A/V167S/G169A. N30_L111I/V136L/S139G/V183I targets three mutations which reside in antiparallel arranged α-helices close to the active-site. N30_L111I/L47W/E74K/V79I/E153Q/Y197H includes all remaining mutations between the two BDH ancestors. These mutations display the largest distance to the active-site of the enzyme. Residues which are conserved between N32 and *Sr*BDH1 are introduced in N30_L111I/V136L/G169A/V183I. Since the most unusual amino acid substitution according to Grantham’s distance (N30_L111I/S232V) is also present in N30_L111I/A222V/E223A/S227A/S232V, N30_L111I/S232V was not investigated further.

Variants N30_L111I/M213I/A214G to N30_L111I/V136L/S139G/V183I displayed an increased activity toward (−)-1-borneol and equal or higher activity toward the preferred enantiomer (+)-1-borneol compared to N30. N30_L111I/L47W/E74K/V79I/E153Q/Y197H and N30_L111I/V136L/G169A/V183I, displayed similar conversion to (+)-1-borneol like N30_L111I, however no conversion of (−)-1-borneol was detectable. N30_L111I/S166A/V167S/G169A further stands out, as displaying the closest (+)-1-borneol conversion to the wildtype, while other variants show rather low activities (**Table 1**).

The peripheral mutations in N30_L111I/L47W/E74K/V79I/E153Q/Y197H are mainly located in loop regions or termini of the α-helices pointing away from the active-site (**Supplementary Fig. 12 A**). In N30_L111I/V136L/G169A/V183I, two mutations (L136 and I183) are located on opposing sites in the αE- and αG-helices with a distance of 9.8 Å and 10.7 Å to the α-carbon of the catalytic Y, while A169 is located in the loop region between αGH close to residue 111 in a distance of 11.0 Å (**Supplementary Fig. 12 B**). Evolutionary coupled residue Ala169 does not influence selectivity in N30_L111I/S166A/V167S/G169A. However, the combination with other conserved residues in N30_L111I/V136L/G169A/V183I shows positive epistatic effects towards increased selectivity.

The peripheral mutations L136, I183, and A169 are also present in different combinations in previously described N30 variants (N30_L111I/S166A/V167S/**G169A** and N30_L111I/**V136L**/S139G/**V183I**). While the combined effect of these three conserved residues results in high enantioselectivity, single or double mutations at these peripheral positions are ineffective, suggesting their synergy in N30. To investigate the role of these residues in the selective N32, the mutant N32_L136V/A169G/I183V was created. N32_L136V/A169G/I183V retained the high enantioselectivity observed in N32, further confirming that enantio-discrimination in ancestor N32 is primarily mediated by I111, which remains unchanged in this variant. The results also indicate that N32 is more strongly adapted to high enantioselectivity than N30. This property is likely stabilized by the combined effect of the remaining 15 residues that differ between N30 and N32. Consistent with this interpretation, high enantioselectivity in N30 can also be acquired by introducing a subset of these substitutions (N30_L111I/L47W/E74K/V79I/E153Q/Y197H, **Table 1**). **Fig. 6** summarizes the mutational strategies described above.

**Figure 6.**
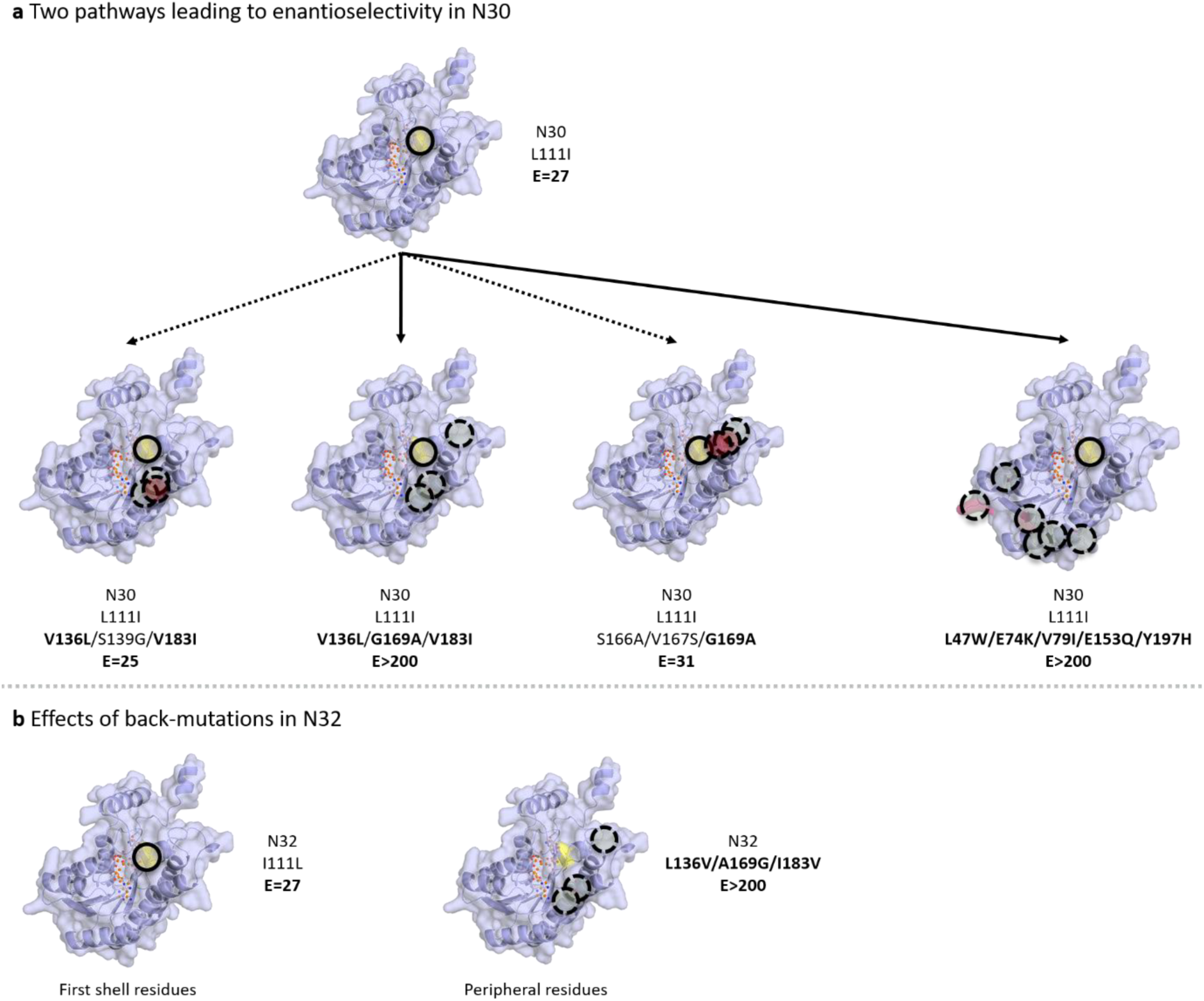
Overview of the changes in enantioselectivity between the N30 and N32 mutants. A) Two separate pathways leading to enantioselectivity in N30. B) Effects of back-mutations on N32 by incorporating residues from N30.

The apparent kinetics of the key variants were assessed to determine whether differences in in enzyme kinetics contribute to the observed enantioselectivity (**Supplementary Fig. 13**, **Supplementary Table 1**). The *K*_app_ was determined as 5.8 ± 2.5 mM for N32_I111L, 18.11 ± 5.0 mM for N30_L111I, and 11.86 ± 1.4 mM for N30_L111I/G169A/V136L/V183I. The *K*_app_ of N30_L111I is comparable to those of N30 and N32, indicating that the change in selectivity introduced by L111I is not driven by altered substrate affinity. In the case of N32_I111L and N30_L111I/G169A/V136L/V183I the *K*_app_ values are lower compared to N30 and N32. However, N30_L111I/G169A/V136L/V183I regains high stereoselectivity despite the reduced *K*_app_, further supporting the conclusion that substrate affinity alone does not determine enantioselectivity. In the case of N32_I111L this comparison is not concise. N32_I111L is a variant of N32 that has lost stereoselectivity; it displays similar specific activity to N30, but has a lower *K*_app_ than that of both variants. Only in this variant could the reduced *K*_app_ be associated with the detectable conversion of (−)-1-borneol. In the case of N30_L111I/V136L/G169A/V183I *K*_app_ could not be extrapolated from the datapoints indicating a high *K*_app_ while retaining enantioselectivity.

### Simulation of BDH selectivity by molecular mechanics

Molecular modeling was employed to elucidate the binding process between borneol and *Sr*BDH1 after repeated attempts to crystallize the substrate proved unsuccessful. Our study aims to uncover the molecular basis of borneol enantioselectivity in both ancestral and current *Sr*BDH1 variants. Publicly available 3D structure indicate that SDRs exhibit two distinct conformations in response to the presence of a substrate or inhibitor. In the unliganded form (PDB IDs: 6ZYZ^19^), the *Sr*BDH1 active site **(Fig. 3**) remains in an ‘open’ conformation, accessible to potential binders. Upon substrate or inhibitor binding, a ‘closed’ conformation is adopted, characterized by the movement of the αFG-helix and loop toward the active site. This conformational shift effectively encapsulates the substrate, optimizing hydrophobic interactions and enhancing catalytic efficiency. These insights into the dynamic structural changes are crucial for understanding the enzyme’s function and selectivity.

We used Chai-1 ^59^ to predict the enzyme-substrate complexes which consistently positioned borneol inside *Sr*BDH1 active-site cavity and reproduce an induced-fit conformational change of surrounding pocket residues observed in the crystal structure (PDB ID: 6ZYZ^19^) increasing the number of contacts within the substrate (**Fig. 7A**). Interestingly, the Chai-1 predicted models contained both (+) and (−)-borneol enantiomers inside the *Sr*BDH1 active site, without discriminating between them. To construct enantiomer-specific systems suitable for comparative simulations, we therefore selected the appropriate borneol orientation and replicated it across the homotetrameric assembly to generate fully consistent (+)-borneol and (−)-borneol complexes. Importantly, the predicted binding geometries were fully compatible with a catalytically competent configuration of borneol (<5Å), as assessed by two key reactive distances (**Figs. 7B, C**, **Supplementary Table 7** and **Supplementary Fig. 14**). These distances correspond to the hydride transfer between the C4 atom of the NAD⁺ nicotinamide ring and the oxidizable C5 carbon of borneol (d_1_, red dashed lines in **Fig. 7 B, C** and **Supplementary Fig. 14**), and to the hydrogen bond between the borneol hydroxyl oxygen and the catalytic tyrosine oxygen (d_2_, blue dashed lines). Because the Chai-1 models yielded nearly identical catalytic distances (d_1_ and d_2_) for (+)- and (−)-borneol, static structures did not explain enantioselectivity.

**Fig. 7.**
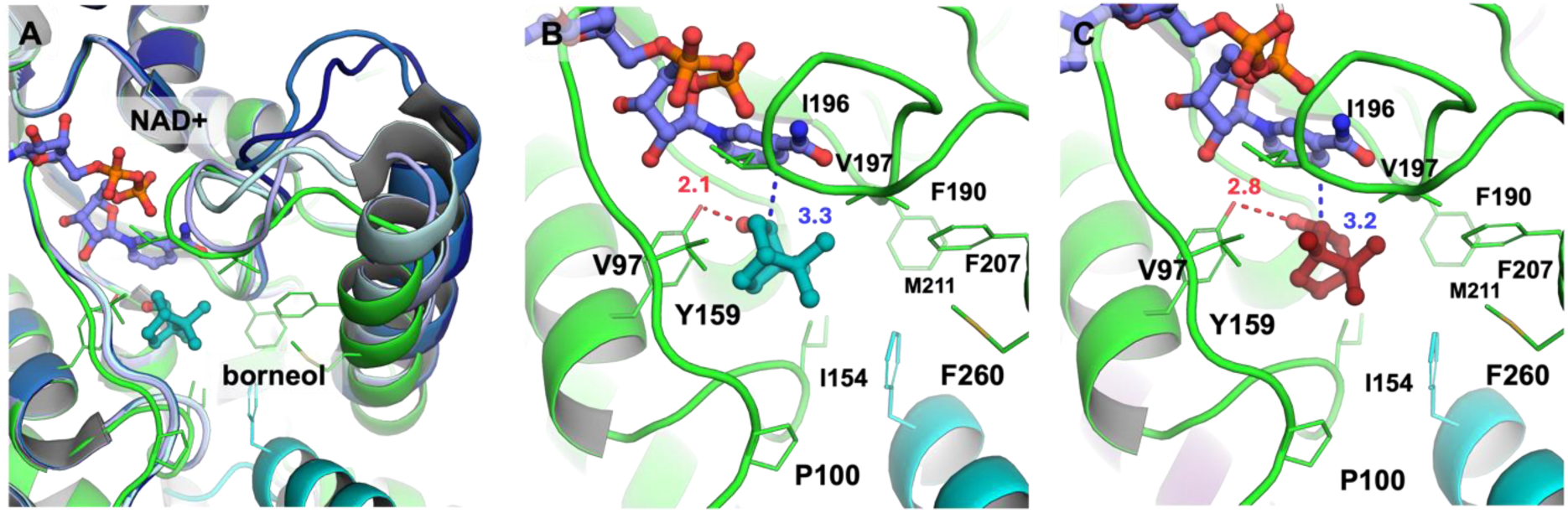
(A) Chai-1^59^predicted model of *Sr*BDH1(green) in complex with borneol, superimposed with 4 different chains from PDB ID: 6ZYZ (blue).^19^ This highlights the induced-fit binding of borneol to *Sr*BDH1 predicted by Chai-1.^59^ (B, C) Models of *Sr*BDH1 in complex with (+)-borneol (teal) and (−)-borneol (brick). Chains A and B are shown in green and blue cartoon representation. Residues interacting with the substrate are represented as sticks. Blue (d_1_) and red (d_2_) dashed lines show the relevant catalytic distances summarized in **Supplementary Table 7**.

Likewise, the predicted structures of variants N32_I111L, N30_L111I, N30_L111I/G169A/V136L/V183I, N6, and N32 failed to discriminate substrate preference based on these catalytic metrics (**Supplementary Fig. 14**, **Supplementary Table 7**). Thus, we turned to molecular dynamic simulations for further insight. Although extensive conventional MD simulations were performed (see **Material and Methods**), these trajectories showed large RMSD fluctuations and rarely maintained catalytically competent geometries, preventing reliable discrimination between (+) and (−)-borneol complexes (**Supplementary Figs. 15** to **26**). We therefore adopted a hybrid machine learning / molecular mechanics (ML/MM) strategy, which consistently stabilized productive enzyme-substrate configurations and resolved enantiomer-specific behaviors. Consequently, the main text focuses on ML/MM results, while detailed comparisons with classical MD are reported in the Supplementary Information (**Fig. 8** and **Supplementary Figures 27** to **38**).

**Fig. 8.**
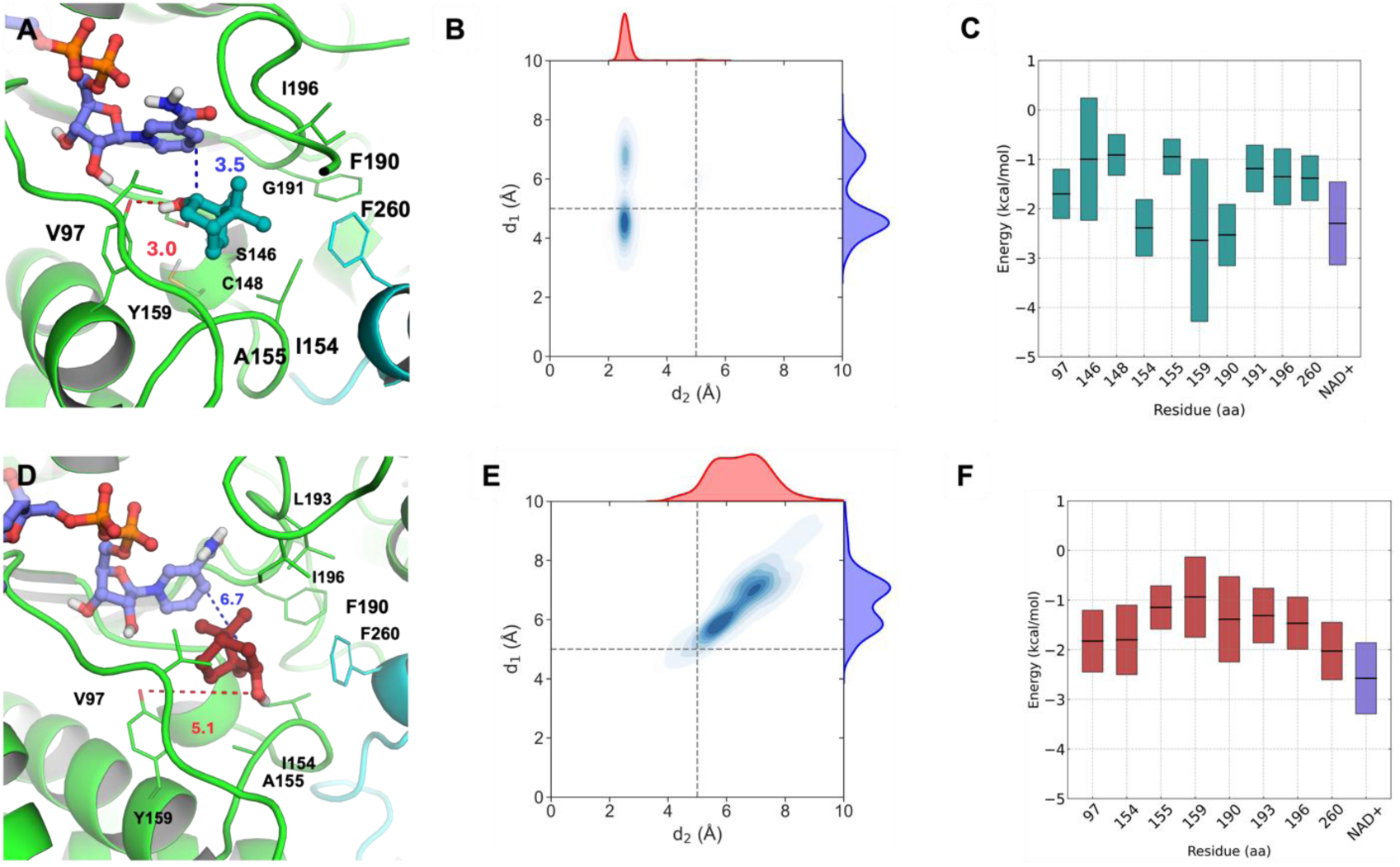
Most populated conformation from each replica, with key active-site residues shown as sticks and distances d_1_ and d_2_ indicated by dashed lines from (A) (+)-borneol and (D) (−)-borneol. Kernel density estimates (KDEs) mean values from 3 x 10 ns ML/MM simulations of the wild-type *Sr*BDH1 complex with (B) (+)-borneol and (E) (−)-borneol. Distance d_1_ (blue) corresponds to the interaction between the catalytic tyrosine and the substrate, while d_2_ (red) represents the hydride-transfer distance (see main text for details). Per-residue energy decomposition mean values obtained from MM-ISMSA^60^analysis showing the residues contributing most strongly to binding in the wild-type *Sr*BDH1 complex with (C) (+)-borneol and (F) (–)-borneol. Box plots show mean values (central black lines) with boxes representing the deviation across replicas.

Analysis of the catalytic distances d_1_ and d_2_ during our ML/MM simulations of the wild-type system in complex with (+)-borneol shows that these distances remain within catalytically competent ranges (**Figs. 8A, B**). In contrast, for the (−)-borneol wild-type complexes (**Figs. 8D, E**), all three replicas exhibit a progressive departure from the catalytic region, with d_1_ and d_2_ shifting away from productive binding ranges. This differential stabilization provides a clear dynamic distinction between the two enantiomers, that is not apparent from conventional MD simulations, at either the same or even 40x-longer time scales (**Supplementary Figs. 25 and 26**).

We subsequently employed MM-ISMSA^60^analysis to compute substrate binding-energy and perform per-residue energy decomposition (**Supplementary Fig. 39**), highlighting key residues involved in borneol binding and elucidating their roles in regioselectivity (**Fig. 8 C and F**). Our results highlight a clear energetic preference of *Sr*BDH1 wildtype for (+)-borneol, with a mean binding energy value 5 kcal/mol more favorable than the one observed for (−)-borneol (**Supplementary Fig. 39 A**). This substantial difference is consistent across replicas and indicates a markedly stronger stabilization of the (+)-enantiomer within the *Sr*BDH1 active site, as indicated also by the monitored distances. Per-residue energy decomposition identifies the catalytic K159 as the residue contributing most strongly to substrate binding further supported by enhanced contributions from I154 and F190, showing significantly more favorable interactions with (+)-borneol whereas (−)-borneol shows increased stabilization from V97 accompanied by reduced interaction with K159.

Solvent-accessible surface area (SASA) values from both the substrate and the enzyme binding pocket were measured along the ML/MM simulations and represented as violin plots (**Supplementary Fig. 39B**). Lower SASA values indicate tighter enclosure of the ligand within the active site, which is expected to favor catalytically competent orientations and thus higher catalytic efficiency^61,62^. Pocket SASA values are 503.9 Å² for (+)-borneol and 546.5 Å² for (−)-borneol, while substrate SASA values are 21.0 Å² and 47.9 Å², respectively in agreement with a more favorable catalytic positioning of the (+)-enantiomer within the *Sr*BDH1 active site. The engineered variants N32_I111L, N30_L111I, N30_L111I/G169A/V136L/V183I, N6, and N32, were analyzed using the previous strategy from wild-type *Sr*BDH1 to obtain corresponding catalytic distances (**Supplementary Figs. 27 to** 32), binding energies (**Supplementary Figs. 40** and **41**) and SASA analysis (**Supplementary Fig. 42**) from 3 replicas of 10ns ML/MM trajectories. For comparison, 10ns conventional MD simulations were again carried out (**Supplementary Figs. 33** to **38**). KDE plots calculated based on these simulations (**Supplementary Figs. 16** and **17**) indicate that variants N32_I111L and N30_L111I do not show a clear discrimination between (+)- and (−)-borneol (**Supplementary Figs. 28** and **29**). In contrast, N30_L111I/G169A/V136L/V183I (**Supplementary Fig. 30**), displays a pronounced preference for (+)-borneol. Among the ancestors, N6 (**Supplementary Fig. 31**) preferentially stabilizes (−)-borneol, whereas N32 (**Supplementary Fig. 32**) favors (+)-borneol, as reflected by their respective catalytic distance distributions.

Consistently, binding-energy analysis (**Supplementary Fig. 40**) shows that for (+)-borneol the most favorable binding is observed for N30_L111I/G169A/V136L/V183I (-23.1 kcal/mol) and N32 (-25.9 kcal/mol), whereas N30_L111I (-20.2 kcal/mol) and N6 (-21.6 kcal/mol) are less favorable. In contrast, for (−)-borneol, N6 (-24.7 kcal/mol) and N32_I111L (-24.7 kcal/mol) show the strongest binding, while N30_L111I/G169A/V136L/V183I exhibits a markedly reduced affinity (-8.0 kcal/mol), highlighting a clear inversion of enantiomer preference across variants.

As observed for the wild-type system, per-residue energy decomposition (**Supplementary Fig. 42**) identifies the catalytic tyrosine as the primary determinant of enantioselectivity. For (+)-borneol, K154 in variants N32_I111L, N30_L111I, N30_L111I/G169A/V136L/V183I, and N32 shows strong stabilizing contributions, with mean interaction energies below -2 kcal/mol, whereas these contributions are markedly reduced for (−)-borneol, particularly in N30_L111I/G169A/V136L/V183I and N32. Tyr185 also contributes significantly, displaying enhanced interactions in N30_L111I, N30_L111I/G169A/V136L/V183I, and N32. In contrast, N6 exhibits a distinct selectivity pattern, where residues in the C-terminal region, L161, V269, and M272, provide increased stabilization of (−)-borneol, thereby driving the observed inversion in enantioselectivity. Overall, these results demonstrate that enantioselectivity in *Sr*BDH1 is governed by an exquisite balance of residue-specific interactions that control substrate orientation and catalytic readiness. This oxidative mechanistic insight provides a clear framework for rationally tuning enzyme selectivity through targeted mutations. SASA analysis (**Supplementary Fig. 42**) shows that for (+)-borneol, the lowest pocket and substrate SASA values are observed for N30_L111I/G169A/V136L/V183I (397.1 and 19.2 Å^2^) and N32 (424.8 and 10.6 Å^2^), consistent with their higher selectivity, whereas N6 and N30_L111I display larger solvent exposure. Conversely, for (−)-borneol this trend is inverted, with N32_I111L (19.2 Å²) and N6 (18.3 Å²) showing the lowest substrate SASA values, and N6 also exhibiting the smallest pocket SASA (468.2 Å²), further supporting their enantiomer-specific selectivity.

### Biased molecular dynamics simulations

Funnel metadynamics simulations^63–67^ were used to further quantify the enantioselective binding process and explicitly sample ligand association and dissociation. Three independent replicas of 300 ns were performed for each enantiomer (**Fig. 9**), enabling extensive sampling of binding and unbinding events (**Supplementary Figs. 43** to **55**). To assess whether observed differences between (+)- and (−)-borneol binding exceed statistical uncertainty, enantioselectivity was quantified as the free-energy difference between enantiomers (ΔΔG), and statistical analyses were performed as described in the **Supplementary Information**.

**Fig. 9.**
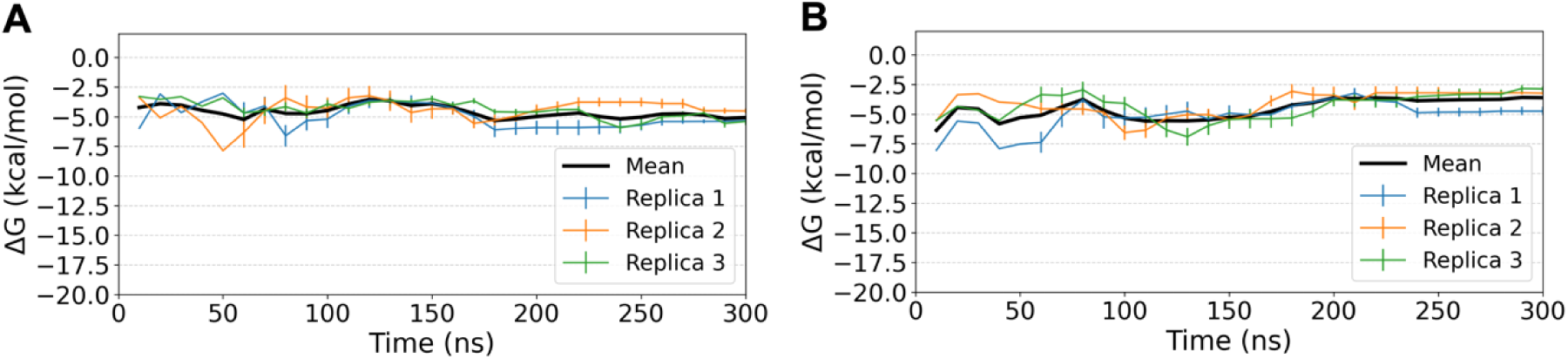
Binding free energy (ΔG) as a function of simulation time for wild-type *Sr*BDH1 in complex with (A) (+)-borneol and (B) (−)-borneol, obtained from funnel metadynamics simulations^63–67^. Data is based on analysis of 3×300 ns simulation trajectories, with individual trajectories shown as colored lines, and the corresponding mean ΔG profile shown in black. Error bars indicate the standard deviation across replicas.

The resulting free-energy profiles (ΔG) (**Supplementary Table 8**) reveal enantiomer-dependent differences in binding stability that correlate with experimentally observed borneol selectivity. In the wild-type enzyme (**Fig. 9**), (+)-borneol displays a deeper binding minimum (ΔG = -5.1 kcal/mol) than (−)-borneol (ΔG = -3.6 kcal/mol), however, the corresponding ΔΔG remains within statistical uncertainty, indicating no significant enantioselective preference. Because binding free energy is directly related to substrate affinity, these values can be qualitatively compared to experimentally measured *K*_app_ values^66,67^. The relatively modest magnitude of the calculated ΔG values (<6 kcal/mol) is consistent with millimolar *K*_app_ values and explains the frequent binding and unbinding events observed in conventional MD simulations (**Supplementary Figs. 15** to **S26**), reflecting the dynamic and reversible nature of substrate recognition in *Sr*BDH1.

For the engineered *Sr*BDH1 variants, funnel-metadynamics simulations reveal mutation-dependent enantioselective binding trends (**Supplementary Table 8**). Variants N32_I111L and N30_L111I display ΔΔG values that do not exceed statistical uncertainty, consistent with reduced or unresolved selectivity. In contrast, variants N30_L111I/G169A/V136L/V183I and N32 exhibit ΔΔG values of approximately -3 kcal/mol, exceeding combined uncertainties and indicating a statistically significant preference for (+)-borneol, while variant N6 shows a comparable preference for (−)-borneol. Overall, the magnitude and direction of the calculated ΔΔG values across variants quantitatively capture the inversion or loss of enantioselectivity observed experimentally. Molecular simulations were employed to elucidate the basis of borneol enantioselectivity in *Sr*BDH1, addressing the lack of structural information due to unsuccessful substrate crystallization.

While Chai-1^59^ static models captured the induced-fit closure of the active site upon ligand binding, they failed to discriminate between (+)- and (−)-borneol, indicating that enantioselectivity arises from dynamic rather than purely structural effects. Conventional MD simulations were unable to resolve these differences due to large fluctuations and loss of catalytically competent geometries. In contrast, TorchANI ML/MM^68–71^ simulations stabilized productive enzyme-substrate complexes and consistently distinguished between enantiomers, highlighting the importance of explicitly accounting for polarization and short-range electronic effects. In the wild-type enzyme, (+)-borneol maintains catalytically competent distances, exhibits lower solvent exposure, and is energetically favored relative to (−)-borneol, with V97, I154 and F190 as the dominant selective residues. Funnel metadynamics further quantified these effects by explicitly sampling ligand binding and unbinding. The resulting free-energy profiles show modest but reproducible differences in binding free energy that correlate with experimental *K*_app_ values, explaining both enantioselectivity and the dynamic, reversible nature of substrate recognition. Extension to engineered variants reveals that enantioselectivity can be attenuated or inverted through subtle redistribution of stabilizing interactions within the active site, rather than through large structural rearrangements. Together, these results demonstrate that *Sr*BDH1 enantioselectivity is governed by differential stabilization of catalytically competent states, and that ML/MM combined with enhanced sampling provides a reliable framework for mechanistic interpretation and rational enzyme engineering.

All of these identified substitutions leading to higher selectivity would be very challenging to predict with our current knowledge on enzyme mechanisms determined by hydrophobic pockets and the available methods for the simulation of interactions in these catalytic motifs. Narayan *et al*.^31^ showed that simply transplanting point mutations between extant enzymes removes amino acids from their natural context, which might interfere with their role in selectivity. While it is not certain that the mutations identified by us represent the true historic trajectory leading to the high stereoselectivity of the *Sr*BDH*s*, all point mutations leading from the common, unselective ancestor N6 of *Sr*BDH1 and to *Aa*BDH2, are amino acid substitutions that occurred in this enzyme family. The amino acid substitutions reported here thus represent a plausible evolutionary pathway to stereoselectivity. In conclusion, our findings highlight how epistatic interactions involving peripheral residues explain the evolutionary development of stereoselectivity, providing valuable insights into the complex interplay between enzyme structure and substrate binding. We believe that this example of the natural emergence of this important catalytic feature will guide future protein engineering endeavors to improve enzyme selectivity.

## Materials and Methods

All chemicals were purchased from Sigma Aldrich (Vienna, Austria) or Roth (Karlsruhe, Germany) and used without further purification unless otherwise stated.

### Plasmids and point mutations

Codon-optimized synthetic gene fragments were either cloned into pET15b or pET28a(+)or purchased in pET28a(+) from GenScript (US) or Twist Bioscience (US) with sequences containing an N-terminal polyhistidine tag. Primers for quick-change mutagenesis were generated by the NEBaseChanger tool (https://nebasechanger.neb.com/). Primer synthesis was facilitated by IDT (US). The reactions were performed according to the Q5^®^ site-directed mutagenesis Kit from NEB (US), whereas chemo-competent *E. coli* TOP10 were used for transformation. Annealing temperatures were adjusted according to the respective primers. The PCR products were treated with 1 μl of DpnI (NEB, US), at 37°C for 1 h followed by inactivation at 80°C for 20 min. After transformation cells were picked for plasmid extraction using the Wizard^®^ *Plus* SV Minipreps DNA Purification System (Promega, US). Plasmid sequences were confirmed by Sanger sequencing (Microsynth, CH) and analyzed by the web tool Benchling (https://benchling.com/).

### Alignment, Phylogenetic tree, ASR

Eleven independent BLAST searches were carried out against the 1KP database^38,39^, with BDH from *Lavandula intermedia* (AFV30207.1, *Li*BDH), alcohol dehydrogenase 2 from *Artemisia annua* (ADK56099.1 *Aa*ADH2), borneol dehydrogenase from *Artemisia annua* (ANJ65952.1, *Aa*BDH), borneol dehydrogenase from *Pseudomonas sp.* TCU-HL1 (WP_032492645.1, *Ps*BDH), borneol-type dehydrogenases 1 and 2 from *Salvia officinalis* (MT525100, *So*BDH1 and MT525099, *So*BDH2), borneol-type dehydrogenases 1 and 2 from *Salvia rosmarinus* (MT857224, *Sr*BDH1 and *Sr*BDH2) and borneol-type dehydrogenases 1 and 2 from *Tanacetum parthenium* (*Tp*BDH1 and *Tp*BDH2) as queries. Hits with an E-value lower than 10^-5^ were selected, and sequence duplicates were deleted and then aligned using ClustalW^36^ and then modified using MUSCLE.^72^ In pairs of sequences displaying a p-distance below 0.25, one sequence was randomly deleted. The remaining sequences were then realigned using ProbCons^73^ and curated using Gblocks^36,37^. The phylogenetic tree was constructed using PhyML^74^, and the ancestral sequence reconstruction based on the evolutionary model Jones-Taylor-Thornton (*JTT*) was carried out by GRASP^75^. PhyML uses Smart Model Selection to select the best fitting amino acid replacement model and uses the approximate Likelihood Ratio Test (aLRT) to calculate branch/node support.

### Growth conditions, protein expression & purification

Initially, *E. coli* Top10 cells were transformed with all BDH gene variants to propagate the plasmid prior to transformation of *E. coli* BL21 (DE3) for over-expression of the BDH enzyme variants.

*E. coli* strains were cultivated in Lennox lysogeny broth (LB-Lennox; 10 g/L tryptone, 5 g/L yeast extract, and 5 g/L NaCl) medium supplemented with a final concentration of 40 mg/mL kanamycin or 100 mg/mL ampicillin. LB-agar plates were prepared similarly with the addition of 15 g/L agar-agar. Precultures were prepared in 50 mL reaction tubes by the addition of material of the corresponding glycerol stock to 25 mL LB supplemented with respective antibiotics. Precultures were cultivated at 37°C, 130 rpm, overnight. Main cultures of 250 mL LB were grown in 1 L baffled flasks with the addition of 40 mg/mL kanamycin or 100 mg/mL ampicillin and inoculated to an OD_600_ of 0.1. Main cultures were then grown at 37°C, 130 rpm until an OD_600_ of 0.8-1.0 and induced by the addition of 1 M (1000x stock) isopropyl-β-D-thiogalactopyranosid.

Cells were harvested by centrifugation (Beckman Coulter, USA, Avanti JXN-26, JA-10 rotor) at 4,000 rpm at 4°C for 15 min. For purification, the harvested cells were resuspended in 20 mL of cell lysis buffer (100 mM KPi, 500 mM NaCl, 10% (v/v) glycerol, pH 8). The suspension was sonicated for 6 min (output control 7, duty cycle 60%) and then centrifuged at 13,200 rpm for 15 min at 4°C (Eppendorf Centrifuge 5415 R). The supernatants were recovered and filtered with 0.45 μm and 0.22 µm filters consecutively. Protein purification was carried out by His-tag affinity chromatography on 2 mL Ni-Sepharose 6 Fast Flow gravity columns (Cytiva, Sweden). Columns stored on 20% EtOH, were washed with 3 column volumes (CV) of dH_2_O. The column was then equilibrated by the addition of 2 × CV binding buffer (100 mM KPi, 500 mM NaCl, 10% (v/v) glycerol, 30 mM imidazole, pH 8) while shaking on ice for 15 min. Following the release of the binding buffer, the filtered cell-free extract was applied onto the column and incubated on ice under slight orbital shaking for 15 min. The flow-through was collected and unspecifically bound protein was eluted by the addition of 5 × CV binding buffer. The target protein was then eluted by the addition of 3 × CV elution buffer (100 mM KPi, 500 mM NaCl, 10% (v/v) glycerol, 500 mM imidazole, pH 8). The purified enzymes were then rebuffered by dialysis (Spectra/Por^®^ 1, MWCO 6-8 kD, Carl Roth, Germany) with storage buffer (100 mM KPi, 500 mM NaCl, 10% glycerol, pH 8) for 4 h at 4°C under gentle orbital shaking. Afterwards, the enzymes were aliquoted and stored with a final concentration of 10% (v/v) glycerol at -20°C until further use.

Protein concentrations were determined spectrophotometrically at 562 nm in a 96-well format, following the Pierce™ BCA Protein Assay Kit protocol (Thermo Fisher Scientific, US). Measurements were performed in triplicates.

SDS-PAGE was conducted to analyze the purified fractions. The samples were mixed with NuPage Sample Reducing Agent (10x), NuPage LDS Sample Buffer (4x) (Thermo Fischer Scientific, US) and dH_2_O to a final concentration of 1 mg/mL.

To denature the protein, samples were heated at 90°C for 10 min. Precast SDS gels (NuPAGE 4-12% Bis-Tris Gel, Thermo Fisher Scientific, US) were inserted into the XCell SureLock Mini-Cell (Thermo Fisher Scientific, US), filled with 1x NuPAGE MES SDS running buffer, prepared from a 20× stock solution (Thermo Fisher Scientific, US). Five μL of the protein standard (PageRuler Prestained, Thermo Fisher Scientific, US) and 10 μL of the samples were loaded onto the gel. The gel was developed under a current of 200 V and 110 mA for 35 min with an electric current generator (PowerEase®500, Invitrogen, US). Afterwards, the gels were stained by submerging them in (de-) staining solution (4.9% (v/v) phosphoric acid, 6.5% (v/v) EtOH, 0.011% (w/v) Coomassie Blue G-250 and 1% (w/v) β-cyclodextrin) for 40 min at room temperature under slow shaking on a tabletop horizontal shaker. Further destaining was facilitated by washing with water at room temperature and continued shaking for 1 h.

### Determination of specific activity and inhibition

Determination of the oxidation rates of extant enzymes, ancestors and ancestor variants towards (+)-1-borneol, (−)-1-borneol or *r*-1-isoborneol was performed with purified enzymes. Reactions containing 0.2, 0.3 or 9 mM of the respective alcohol (dissolved in DMSO), 1 or 9 mM NAD^+^, 20% total DMSO in 0.1 M Tris-HCl (pH 9) were started by the addition of 2 µM purified enzyme (final concentration). The reactions were carried out at 25°C, 300 rpm. Over time, multiple samples were taken and stopped by 1:1 addition of 1 M NaOH. Substrate and product were extracted via a two-step extraction process with 1:2 MTBE, the organic layers were collected and dried over Na_2_SO_4_ and product formation was quantified via gas chromatography. Experiments were carried out in triplicates.

### Determination of kinetic parameters

Kinetic parameters of N32, N30 and N6 were obtained by measuring initial velocities in the presence of varying concentrations of (+)-1-borneol (0.02 or 0.05–9 mM) at constant 9 mM of NAD^+^ under the conditions given above. Kinetic parameters were obtained by fitting the obtained data to the Michaelis-Menten equation with SigmaPlot 14.0 (Systat Software). Experiments were carried out in quintuplicates.

### Determination of E-value

Determination of the selectivity expressed through the enantiomeric ratio (E-value) was carried out by following the principle of the Quick-E method developed by Kazlauskas *et al*.^46^ with slight modifications. In the Quick-E method, initial rates of the enantiopure substrate are measured in competition with a reference compound via a kinetic spectroscopic measurement. In this adaptation of the assay, the competitive oxidation of a surrogate substrate mixed with the enantiopure alcohol occurs, and the reaction is analyzed by GC analysis. Determination of the reaction rates under competitive conditions is carried out under the same conditions as mentioned above, with the addition of equimolar amounts of either cyclohexanol or n-octanol.

### Quantitative GC analysis

Achiral GC analysis of 1-borneol and camphor was carried out on a Shimadzu GC-2030 GC-FID system equipped with a ZB-5 column (30 m length, 0.32 mm inner diameter, 0.25 µM film thickness, Shimadzu, JP). Samples were injected in split mode (9.1) with an injection temperature of 250°C and FID temperature of 320°C. The instrument was programmed to an initial oven temperature of 50°C. After 5 min of holding time, a linear increase of 40°C/min up to 300°C was set, followed by 5 min holding. N_2_ was set to a constant flow of 30 mL/min. Achiral analysis of reactions containing an additional surrogate substrate was carried out with the same method, on a Shimadzu GC-2010 Plus GCMS-QP2010 SE system equipped with a ZB5-ms column (30 m length, 0.25 mm inner diameter, 0.25 µM film thickness, Shimadzu, JP) with helium as carrier gas. Carrier gas flow was set to a constant flow of 15 mL/min, while the ion source temperature was set to 250°C and the interface temperature to 320°C. Chiral analysis of *r*-1-isoborneol was carried out on a Shimadzu GC-2030 GC-FID system equipped with a Hydrodex βTBDM column (25 m length, 0.25 mm inner diameter, Macherey-Nagel, Germany). Samples were injected in split mode (100:1 or 50:1) at an injection temperature of 250°C and FID temperature of 320°C. The instrument was programmed to an initial oven temperature of 90°C. After 5 min of holding time, a linear increase of 2°C/min up to 150°C was set, followed by a linear increase of 45°C/min up to 200°C and (2 min holding). N_2_ was set to a constant flow of 24 mL/min.

### Protein expression and purification for crystallization

Purification of N32 and N39 was conducted as mentioned before.^19^ *E. coli* BL21-RIL cells were transformed with a pET28a(+) vector containing either N32 or N39 fused to an N-terminal hexa-histidine-tag. Protein production was carried out in auto-induction media at 37°C for 4 h and subsequently cooled down to 16°C.^76^ Cells were grown overnight and harvested by centrifugation (10 min, 6,000 rpm at 4°C). The pellets were resuspended in buffer A (20 mM Tris/HCl pH 8.0, 500 mM NaCl). Cells were lysed by sonification at 4°C for 15 min after the addition of 0.5 mg/l DNase and the lysate was cleared by centrifugation (30 min, 21,500 rpm at 4°C). A Ni^2+^NTA column (1 CV = 1 mL, Macherey-Nagel) was equilibrated with buffer A and the lysed suspension of N32 or N39 was loaded on the column and washed with 15 CV of buffer A. The proteins of interest were eluted with buffer A supplemented with 300 mM imidazole. An additional size-exclusion chromatography (SEC) step was performed with a HighLoad Superdex S200 16/60 column (GE Healthcare), equilibrated with 20 mM Tris/HCl pH 8.0, 125 mM NaCl. Pooled protein fractions were concentrated with Amicon-Ultra-15 (Merck KGaA) to 29 mg/mL for N32 or 28 mg/mL for N39 as measured by the absorbance at 280 nm.

### Crystallization

Crystallization was conducted as described previously.^19^ Crystals of N32 were obtained by the sitting-drop vapor-diffusion method at 18°C with a reservoir solution composed of 0.1 M HEPES/NaOH pH 7.6, 0.2 M proline, 12% (w/v) PEG 3,350. Crystals were cryo-protected with 30 % (v/v) glycerol supplemented to the reservoir resolution and subsequently flash-cooled in liquid nitrogen. Crystals of N39 were obtained with a reservoir solution composed of 0.1 M HEPES/NaOH pH 7.4, 11% (w/v) PEG 3,000. Initially, inter-grown crystals were used to prepare a seed stock. With a cat whisker, seeds were transferred to a freshly prepared crystallization drop with the mentioned buffer composition and crystals were subsequently grown at 18°C. Prior to flash-freezing in liquid nitrogen, the fished crystals were transferred to a cryo-protectant solution composed of the reservoir solution supplemented with 25% (v/v) PEG 400.

### X-ray data collection, structure determination and refinement

Synchrotron diffraction data were collected at the beamline 14.2 of the MX Joint Berlin laboratory at BESSY (Berlin, Germany) and beamline P11 of PETRA III (Deutsches Elektronen Synchrotron, Hamburg, Germany) at 100 K. Diffraction data were processed with XDS (**Supplementary Table 4**).^77^ The structures were solved by molecular replacement using PHASER^78^ with a search model comprising a single protomer of *Sr*BDH1 (PDB: 6ZZ0^19^). The structure was refined by maximum-likelihood restrained refinement in PHENIX.^79,80^ Model building was performed with COOT.^81,82^ Model quality was evaluated with MolProbity^83,84^ and the JCSG validation server (JCSG Quality Control Check v3.1). Secondary structure elements were assigned with DSSP.^85^ Structure-based alignments were visualized by Alscript.^86^ Figures were prepared using PyMOL (Schrödinger Inc).

### SrBDH1 models

Structures of protein–ligand complex were generated using Chai-1^59^Haz clic o pulse aquí para escribir texto., from the corresponding FASTA sequences and the SMILES codes of NAD^+^ and borneol. The Chai-1 model contained either (+) or (−)-borneol bound in a single chain; to obtain the full tetrameric assembly, the corresponding ligand-bound chain was duplicated to reconstruct the complete tetramer using PyMol.^87^ Hydrogen atoms were added, and the protonation states of titratable residues were adjusted to pH 7.0, using the H++ webserver^88,89^, while ensuring the catalytic Tyr remained deprotonated. Hydrogens for the oxidized state of the cofactor NAD^+^ were added using Open Babel.^90^.

### Molecular dynamics simulations

The initial structures were obtained from previous docking models, and borneol poses were replicated across each chain (**Figs. 6C** and **D**). After that, protein residues were described using the AMBER ff19SB^91^ force field, while partial charges for the NAD^+^, borneol substrates and the deprotonated tyrosine were obtained the HF/6-31G(d) level of theory using restrained electrostatic potential (RESP)^92^ fitting with Antechamber (GAFF2^93^), based on gas-phase geometries optimized at the B3LYP/6-31G(d) level of theory using Gaussian 16.^94^ The remaining parameters were also obtained using GAFF2. The final systems were parametrized using the tleap module from Amber Tools.^95^We conducted an initial *in vacuo* minimization of our *Sr*BDH1 systems using sander molecular mechanics module from from Amber 24.^95,96^ After that, all systems were embedded in a truncated octahedral water box with a minimum solute–box edge distance of 13 Å, setting the stage for conventional molecular dynamics (MD) and ML/MM simulations.

Our MD and ML/MM systems used a shared equilibration protocol. Initially, all systems were energetically minimized to eliminate steric clashes. Each system was then centered in a octahedral box containing between 21,000 and 23,000 TIP3P water molecules depending on system. Subsequent minimization occurred in three phases, first targeting all protons, followed by the solvent, and finally the entire system. The systems were gradually heated from 100K to 300K over 50 ps under an NVT ensemble using a Langevin thermostat with a gamma friction coefficient of 1.0 ps^-1^. During heating, harmonic constraints of 40 kcal/mol·Å^-2^ were applied to all solute atoms. These constraints were systematically reduced to 10 kcal/mol·Å^-2^ over four simulation stages at 300 K under NVT conditions. Transitioning to an NPT ensemble, we employed the Berendsen barostat^97^ at 300K. We then continued the simulations for an additional 100 ns, applying positional restraints to the borneol molecules to allow the enzyme to equilibrate around the substrate across three independent replicas, after which we performed unconstrained 400 MD simulations (see **Supplementary Figs. 15** to **28** f**or the monitored RMSD values and catalytic distances**). A cutoff of 10.0 Å was used for calculating van der Waals interactions, and long-range electrostatics were managed *via* the Particle-Mesh Ewald (PME)^98^ method. The SHAKE^99^ algorithm was employed to restrain the hydrogen atoms in water molecules. Analysis of the MD trajectories, including monitoring RMSD and distances d_1_ and d_2_, was conducted using the CPPTRAJ^100^ module from Amber Tools.^95^

Equilibrated MD structures were used as starting points for TorchANI-AMBER ML/MM^68–71^ simulations implemented in AmberTools^101–103^, where the ligand is described by a neural-network potential trained on high-level quantum chemical data, enabling explicit treatment of polarization and short-range electronic effects at near QM/MM accuracy. For each enantiomeric complex, three independent 10 ns ML/MM replicas were generated following a validated protocol3, providing sufficient sampling for robust characterization of enantioselective interactions.

Borneol molecules bound in chain A were selected as the ML region and treated with the ANI potentials, while the remainder of the system was described at the MM level. ML/MM coupling was enabled using electrostatic embedding, with a ML-MM non-bonded cutoff of 12.0 Å. The ML region was assigned a net charge of 0 and a singlet spin state. All simulations were run for 10 ns, without constraining QM hydrogen atoms, and without Ewald or PME treatment for ML-ML and ML–MM electrostatics. Torch-based coupling was employed throughout the simulations, ensuring consistent force evaluation between the ML and MM regions. For comparison we run in parallel 10 ns of conventional MD simulations (**Supplementary Fig. S33-38**).

### Funnel metadynamics simulations

We employed funnel metadynamics, an enhanced-sampling approach that applies a restraining funnel-shaped potential to efficiently explore ligand entry, binding, and exit from the active-site cavity^63–67^. Simulations were initiated from the equilibrated systems described above, and bias was applied exclusively to a single borneol molecule, following established protocols^66,67^. Funnel coordinates were generated following the tutorial by Dominykas Lukauskis on funnel maker (https://github.com/dlukauskis/funnel_maker/tree).^104^ These simulations were performed as three independent replicas of 300 ns each using PLUMED 2.9^105,106^. Well-tempered metadynamics was applied by depositing Gaussian hills every 500 steps with an initial height of 0.25 kcal/mol and Gaussian widths of 0.5 and 0.3 along the projection and extension funnel coordinates, respectively. A bias factor of 10 Kcal/mol was used at a temperature of 300 K to ensure smooth convergence of the free-energy surface while avoiding overfilling. Free-energy surface and PMF values were obtained using PLUMED^105^ and graphics analysis with matplotlib^107^. Convergence of the simulations was carefully assessed by monitoring the time evolution of the biased collective variables and the reconstructed free-energy surfaces (**Supplementary Figures 33** to **45**). Upper walls were set up at 20 Å and 8 Å at pp.proj and pp.ext respectively.

## Notes

The atomic coordinates have been deposited in the Protein Data Bank with the accession code 8R0C (N32) and 8R0D (N39). Diffraction raw images have been deposited at proteindiffraction.org: 10.18430/M38R0C (N32) and 10.18430/M38R0D (N39). Raw data of the biochemical analysis, structure predictions and simulation data have been deposited at Zenodo. The package containis chromatograms of three kinetic measurements, the extracted peak areas and calculations. The data is located at DOI: 10.5281/zenodo.18925996.

## Acknowledgments

This research was funded in part by the Austrian Science Fund (FWF) (Grant-dois: 10.55776/P34280, 10.55776/COE17). This project has received funding from the European Union’s Horizon Europe research and innovation programme under the Marie Skłodowska-Curie grant agreement No 101072686. We accessed beamlines of the BESSY II (Berliner Elektronenspeicherring-Gesellschaft für Synchrotronstrahlung II) storage ring (Berlin, Germany) via the Joint Berlin MX-Laboratory sponsored by the Helmholtz Zentrum Berlin für Materialien und Energie, the Freie Universität Berlin, the Humboldt-Universität zu Berlin, the Max-Delbrück Centrum and the Leibniz-Institut für Molekulare Pharmakologie. Parts of this research were carried out at PETRA III at DESY, a member of the Helmholtz Association (HGF). S. C. L. K. acknowledges support from Georgia Tech, the Vasser Wooley Foundation and the Georgia Research Alliance. We acknowledge National Academic Infrastructure for Supercomputing in Sweden (NAISS) for awarding this project access to the LUMI supercomputer, owned by the EuroHPC Joint Undertaking, hosted by CSC (Finland) and the LUMI consortium through grant agreement no. NAISS 2023/8-3 and NAISS 2024/8-16. This work used the Hive cluster, which is supported by the National Science Foundation under grant number 1828187. This research was supported in part through research cyberinfrastructure resources and services provided by the Partnership for an Advanced Computing Environment (PACE) at the Georgia Institute of Technology, Atlanta, Georgia, USA. C.P.O.H. was supported by a scholarship from the Hanns Seidel Foundation with funds from Federal Ministry of Education and Research Germany (BMBF), as well as by the Deutsche Forschungsgemeinschaft (Grant RTG 2473-1). D.K. acknowledges the BioTechMed-Graz Young Researcher Group programme. R. T.J. was supported by EU grant. R.T.J. was supported by European Union’s Horizon Europe research and innovation programme under the Marie Skłodowska-Curie grant agreement No 101072686. We thank Romas Kazlauskas for the discussion about the potential influence of peripheral residues on residues located in the active site.

## Contributions

J.Z., K.K., and A.M.C. performed the experiments. C.P.O.H. and R.T-J. conducted crystal structure determination. B.D.G. performed MD simulations. S.B., N.R.M. and E.A.G. conducted ancestral sequence reconstruction (ASR). S.C.L.K and R.K. conceptualized the project and the main ideas. All authors discussed results, contributed to the manuscript writing and provided critical comments.

## Competing interests

The authors declare no competing interests.

## Notes

### Competing Interest Statement

The authors have declared no competing interest.

### Summary of Updates

This version of the manuscript has been revised to strengthen the phylogenetic framework, kinetic characterization, computational modelling, and overall presentation. The phylogenetic analysis and ancestral sequence reconstruction sections were expanded to include explicit descriptions of sequence selection, alignment curation, model selection, rooting strategy, and confidence estimation. Revised phylogenetic figures now include branch support values, posterior probabilities, and improved visualization with collapsed clades and supplementary trees. Additional mutational and kinetic analyses were incorporated to further investigate the emergence of enantioselectivity in borneol dehydrogenases. A new back-mutant of N32 was generated and characterized, confirming the central role of residue I111 in stereodiscrimination while supporting the contribution of additional adaptive mutations. Apparent kinetic parameters were determined for additional variants, and the manuscript now clarifies the experimental limitations caused by low substrate solubility and high apparent KM values. Kinetic values are now consistently reported as apparent Kapp values. Representative raw chromatographic and kinetic datasets were deposited to Zenodo and referenced throughout the manuscript. The computational workflow was substantially revised. The original docking-based models were replaced with deep-learning-based enzyme-substrate complex prediction using Chai-1. These refined complexes were subsequently analyzed using both conventional molecular dynamics and ML/MM simulations with TorchANI-AMBER. The revised simulations improved structural stability and provided clearer discrimination between borneol enantiomers. Free-energy calculations and potential-of-mean-force analyses were also added, revealing quantitative agreement between computed energetic profiles and experimental kinetic trends. Multiple figures and supplementary figures were redesigned or expanded to improve clarity and consistency. Figure annotations, legends, structural representations, and supplementary descriptions were revised extensively, including clarification of catalytic distances, active-site representations, loop conformations, and crystallographic quality metrics. Additional structural comparison figures and computational analyses were added to support the revised mechanistic interpretation. Due to the revision of the ancestral sequence reconstruction, two additional authors were added.

## References

1. Ahmed, S. N. et al. Enantioselectivity of *Candida rugosa* lipase toward carboxylic acids: a predictive rule from substrate mapping and X-ray crystallography. Biocatalysis 9, 209–225 (1994).

2. Buller, R. et al. From nature to industry: harnessing enzymes for biocatalysis. Science 382, (2023).

3. Kazlauskas, R. J. & Bornscheuer, U. T. Finding better protein engineering strategies. Nat. Chem. Biol. 5, 526–529 (2009).

4. Petersen, B. M. et al. An integrated technology for quantitative wide mutational scanning of human antibody Fab libraries. Nat. Commun. 15, 3974 (2024).

5. Gantz, M. et al. Microdroplet screening rapidly profiles a biocatalyst to enable its AI-assisted engineering. bioRxiv 10.1101/2024.04.08.588565 (2024).

6. Croteau, R., Felton, M., Karp, F. & Kjonaas, R. Relationship of camphor biosynthesis to leaf development in sage (*Salvia officinalis*). Plant Physiol. 67, 820–824 (1981).

7. Whittington, D. A. et al. Bornyl diphosphate synthase: structure and strategy for carbocation manipulation by a terpenoid cyclase. Proc. Natl Acad. Sci. U.S.A. 99, 15375–15380 (2002).

8. Ma, R. et al. Identification of (−)-bornyl diphosphate synthase from *Blumea balsamifera* and its application for (−)-borneol biosynthesis in *Saccharomyces cerevisiae*. Synth. Syst. Biotechnol. 7, 490–497 (2022).

9. Polichuk, D. R. et al. A glandular trichome-specific monoterpene alcohol dehydrogenase from *Artemisia annua*. Phytochemistry 71, 1264–1269 (2010).

10. Sarker, L. S. et al. Molecular cloning and functional characterization of borneol dehydrogenase from the glandular trichomes of *Lavandula × intermedia*. Arch. Biochem. Biophys. 528, 163–170 (2012).

11. Ma, R. et al. Molecular cloning and functional identification of a high-efficiency (+)-borneol dehydrogenase from *Cinnamomum camphora*. Plant Physiol. Biochem. 158, 363–371 (2021).

12. Tian, N. et al. Molecular cloning and functional identification of a novel borneol dehydrogenase from *Artemisia annua*. Ind. Crops Prod. 77, 190–195 (2015).

13. Hu, X. et al. Heterologous expression and characterization of a borneol dehydrogenase from *Arabidopsis lyrata*. Mol. Catal. 530, 112572 (2022).

14. Lin, X. et al. Genome-wide identification and functional characterization of borneol dehydrogenases in *Wurfbainia villosa*. Planta 258, 1–16 (2023).

15. Satyal, P. et al. Chemotypic characterization and biological activity of *Rosmarinus officinalis*. Foods 6, 1–15 (2017).

16. Croteau, R., Lee Hooper, C. & Felton, M. Biosynthesis of monoterpenes: partial purification and characterization of a bicyclic monoterpenol dehydrogenase from sage. Arch. Biochem. Biophys. 188, 182–193 (1978).

17. Drienovská, I. et al. Molecular cloning and functional characterization of two highly stereoselective borneol dehydrogenases from *Salvia officinalis*. Phytochemistry 172, 112227 (2020).

18. Dimos, N. et al. Cryo-EM analysis of small plant biocatalysts at sub-2 Å resolution. Acta Crystallogr. D Struct. Biol. 78, 113–123 (2022).

19. Chánique, A. M. et al. A structural view on the stereospecificity of plant borneol-type dehydrogenases. ChemCatChem 13, 2262–2277 (2021).

20. Calderini, E. et al. Simple plug-in synthetic step for the synthesis of (−)-camphor from renewable starting materials. ChemBioChem 22, 2951–2956 (2021).

21. Kavanagh, K. L. et al. Medium- and short-chain dehydrogenase/reductase gene and protein families: the SDR superfamily. Cell. Mol. Life Sci. 65, 3895 (2008).

22. Okrasa, K. et al. Structure and mechanism of an unusual malonate decarboxylase and related racemases. Chem. Eur. J. 14, 6609–6613 (2008).

23. Biler, M. et al. Ground-state destabilization by active-site hydrophobicity controls selectivity. J. Am. Chem. Soc. 142, 20216–20231 (2020).

24. Burton, A. J. et al. Installing hydrolytic activity into a de novo protein framework. Nat. Chem. 8, 837–844 (2016).

25. Kalvet, I. et al. Design of heme enzymes with tunable substrate binding pocket. J. Am. Chem. Soc. 145, 14307–14315 (2023).

26. Sandström, A. G. et al. Directed evolution of *Candida antarctica* lipase A. Protein Eng. Des. Sel. 22, 413–420 (2009).

27. Sandström, A. G. et al. Combinatorial reshaping of lipase substrate pocket. Proc. Natl Acad. Sci. U.S.A. 109, 78–83 (2012).

28. Bartsch, S., et al. Complete inversion of enantioselectivity. Angew. Chem. Int. Ed. 47, 1508–1511 (2008).

29. Brundiek, H. B., et al. Creation of a lipase selective for trans fatty acids. Angew. Chem. Int. Ed. 51, 412–414 (2011).

30. Jan Deniau, A., et al. Characterization and reshaping of nucleophile pocket. ChemBioChem 19, 1839–1844 (2018).

31. Chiang, C. H. et al. Evolution of monooxygenase stereoselectivity. Proc. Natl Acad. Sci. U.S.A. 120, e2218248120 (2023).

32. Hochberg, G. K. A. & Thornton, J. W. Reconstructing ancient proteins. Annu. Rev. Biophys. 46, 247–269 (2017).

33. Selberg, A. G. A. et al. Ancestral sequence reconstruction. J. Mol. Evol. 89, 157–164 (2021).

34. Zhu, X. X. et al. Evolutionary insights into stereoselectivity. Nat. Commun. 15, (2024).

35. Eggerichs, D. et al. Vanillyl alcohol oxidase from *Diplodia corticola*. J. Biol. Chem. 298, 104898 (2023).

36. Schriever, K. et al. Engineering of ancestors for terpene cyclase. J. Am. Chem. Soc. 143, 3794–3807 (2021).

37. Vongsouthi, V. et al. Ancestral reconstruction of PET-degrading cutinases. Sci. Adv. 11, eads8318 (2025).

38. Leebens-Mack, J. H. et al. One thousand plant transcriptomes. Nature 574, 679–685 (2019).

39. Carpenter, E. J. et al. 1000 plant transcriptomes initiative. GigaScience 8, giz126 (2019).

40. Polichuk, D. R. et al. A glandular trichome-specific enzyme. Phytochemistry 71, 1264–1269 (2010).

41. Tsang, H. L. et al. Borneol dehydrogenase from *Pseudomonas*. Appl. Environ. Microbiol. 82, 6378–6385 (2016).

42. Drienovská, I. et al. Borneol dehydrogenases. Phytochemistry 172, 112227 (2020).

43. Yalkowsky, S. H., et al. Handbook of aqueous solubility data (2010).

44. Straathof, A. J. J. et al. Production of fine chemicals. Curr. Opin. Biotechnol. 13, 548–556 (2002).

45. Chen, C. S. et al. Quick E method. Rec. Trav. Chim. Pays-Bas 104, 4560–4561 (1992).

46. Janes, L. E. & Kazlauskas, R. J. Quick E method. J. Org. Chem. 62, 4560–4561 (1997).

47. Tanaka, N. et al. Crystal structures of dehydrogenase. Biochemistry 35, 7715–7730 (1996).

48. Filling, C. et al. Critical residues in SDR. J. Biol. Chem. 277, 25677–25684 (2002).

49. Oppermann, U. et al. SDR update. Chem. Biol. Interact. 143–144, 247–253 (2003).

50. Shah, B. S. et al. SDR structure in *Acinetobacter*. Acta Crystallogr. F 70, 1318–1323 (2014).

51. Chánique, A. M. et al. A structural view on the stereospecificity of plant borneol-type dehydrogenases. ChemCatChem 13, 2262–2277 (2021).

52. Abramson, J. et al. Accurate structure prediction of biomolecular interactions with AlphaFold 3. Nature 630, 493–500 (2024).

53. Lipsh-Sokolik, R. & Fleishman, S. J. Addressing epistasis in the design of protein function. Proc. Natl Acad. Sci. U.S.A. 121, e2314999121 (2024).

54. Yang, Z. PAML 4: phylogenetic analysis by maximum likelihood. Mol. Biol. Evol. 24, 1586–1591 (2007).

55. Korman, T. P. et al. Structural analysis of actinorhodin polyketide ketoreductase. Biochemistry 43, 14529–14538 (2004).

56. Kumar, R. et al. Crystal structure and characterization of NADP+-farnesol dehydrogenase. Insect Biochem. Mol. Biol. 147, 103812 (2022).

57. Grantham, R. Amino acid difference formula to explain protein evolution. Science 185, 862–864 (1974).

58. Hopf, T. A. et al. The EVcouplings Python framework. Bioinformatics 35, 1582–1584 (2019).

59. Team, C. D. et al. Chai-1: decoding molecular interactions of life. bioRxiv 10.1101/2024.10.10.615955 (2024).

60. Klett, J. et al. MM-ISMSA: scoring function for protein–protein docking. J. Chem. Theory Comput. 8, 3395–3408 (2012).

61. Geronimo, B., et al. Molecular connections between catalytic mechanism and pathophysiology. J. Chem. Inf. Model. 10.1021/acs.jcim.4c02229 (2025).

62. Jiang, Y. et al. Substrate positioning dynamics in catalysis. J. Phys. Chem. Lett. 14, 11480–11489 (2023).

63. Karrenbrock, M. et al. Absolute binding free energies with OneOPES. J. Phys. Chem. Lett. 15, 9871–9880 (2024).

64. Barducci, A., Bussi, G. & Parrinello, M. Well-tempered metadynamics. Phys. Rev. Lett. 100, 020603 (2008).

65. Barducci, A., Bonomi, M. & Parrinello, M. Metadynamics. WIREs Comput. Mol. Sci. 1, 826–843 (2011).

66. Limongelli, V., Bonomi, M. & Parrinello, M. Funnel metadynamics. Proc. Natl Acad. Sci. U.S.A. 110, 6358–6363 (2013).

67. Raniolo, S. & Limongelli, V. Ligand binding free-energy calculations. Nat. Protoc. 15, 2837–2866 (2020).

68. Pickering, I. et al. TorchANI-amber: bridging neural networks and simulations. J. Phys. Chem. B 129, 11927–11938 (2025).

69. Semelak, J. A. et al. Advancing multiscale molecular modeling with ML electrostatics. J. Chem. Theory Comput. 21, 5194–5207 (2025).

70. Gao, X. et al. TorchANI: deep learning implementation of ANI potentials. J. Chem. Inf. Model. 60, 3408–3415 (2020).

71. Pickering, I. et al. TorchANI 2.0: extensible high-performance library. J. Chem. Inf. Model. 65, 11656–11671 (2025).

72. Edgar, R. C. MUSCLE: multiple sequence alignment. Nucleic Acids Res. 32, 1792–1797 (2004).

73. Do, C. B. et al. ProbCons alignment method. Genome Res. 15, 330–340 (2005).

74. Guindon, S. & Gascuel, O. Maximum likelihood phylogeny algorithm. Syst. Biol. 52, 696–704 (2003).

75. Foley, G., et al. Engineering enzyme variants using GRASP. bioRxiv 10.1101/2019.12.30.891457 (2022).

76. Studier, F. W. Protein production by auto-induction. Protein Expr. Purif. 41, 207–234 (2005).

77. Kabsch, W. XDS. Acta Crystallogr. D Biol. Crystallogr. 66, 125–132 (2010).

78. McCoy, A. J. et al. Phaser crystallographic software. J. Appl. Crystallogr. 40, 658–674 (2007).

79. Liebschner, D. et al. Macromolecular structure determination using PHENIX. Acta Crystallogr. D Struct. Biol. 75, 861–877 (2019).

80. Adams, P. D. et al. PHENIX system for structure solution. Acta Crystallogr. D Biol. Crystallogr. 66, 213–221 (2010).

81. Emsley, P. et al. Features and development of Coot. Acta Crystallogr. D Biol. Crystallogr. 66, 486–501 (2010).

82. Casañal, A. et al. Developments in Coot for cryo-EM. Protein Sci. 29, 1069–1078 (2020).

83. Williams, C. J. et al. MolProbity: improved structure validation. Protein Sci. 27, 293–315 (2018).

84. Chen, V. B. et al. MolProbity: all-atom structure validation. Acta Crystallogr. D Biol. Crystallogr. 66, 1–10 (2010).

85. Kabsch, W. & Sander, C. Dictionary of protein secondary structure. Biopolymers 22, 2577–2637 (1983).

86. Barton, G. J. ALSCRIPT: formatting sequence alignments. Protein Eng. 6, 37–40 (1993).

87. DeLano, W. L. PyMOL: an open-source molecular graphics tool. Protein Crystallogr. Newsl. 40, (2002).

88. Gordon, J. C. et al. H++: estimating pKa and adding hydrogens. Nucleic Acids Res. 33, (2005).

89. Anandakrishnan, R. et al. H++ 3.0 for biomolecular preparation. Nucleic Acids Res. 40, 537–541 (2012).

90. O’Boyle, N. M. et al. Open Babel: chemical toolbox. J. Cheminform. 3, 33 (2011).

91. Tian, C. et al. ff19SB force field. J. Chem. Theory Comput. 16, 528–552 (2020).

92. Woods, R. J. & Chappelle, R. Restrained electrostatic potential charges. J. Mol. Struct. THEOCHEM 527, 149–156 (2000).

93. He, X. et al. High-quality charge model for AMBER force field. J. Chem. Phys. 153, (2020).

94. Frisch, M. J. et al. Gaussian 16 (Gaussian, Inc., 2016).

95. Case, D. A. et al. AmberTools. J. Chem. Inf. Model. 63, 6183–6191 (2023).

96. Case, D. A. et al. AMBER (University of California, San Francisco, 2024).

97. Berendsen, H. J. C. et al. Molecular dynamics with coupling to an external bath. J. Chem. Phys. 81, 3684–3690 (1984).

98. Darden, T. et al. Particle mesh Ewald method. J. Chem. Phys. 98, 10089–10092 (1993).

99. Ryckaert, J.-P. et al. Numerical integration of constrained systems. J. Comput. Phys. 23, 327–341 (1977).

100. Roe, D. R. & Cheatham, T. E. CPPTRAJ analysis software. J. Chem. Theory Comput. 9, 3084–3095 (2013).

101. Case, D. A. et al. AMBER 25 (University of California, San Francisco, 2025).

102. Case, D. A. et al. AmberTools. J. Chem. Inf. Model. 63, 6183–6191 (2023).

103. Case, D. A. et al. Recent developments in AMBER simulations. J. Chem. Inf. Model. 65, 7835–7843 (2025).

104. Hedges, L. O. et al. Tutorials for BioSimSpace. Living J. Comput. Mol. Sci. 5, (2023).

105. Tribello, G. A. et al. PLUMED 2. Comput. Phys. Commun. 185, 604–613 (2014).

106. Bonomi, M. et al. Transparency in molecular simulations. Nat. Methods 16, 670–673 (2019).

107. Hunter, J. D. Matplotlib: a 2D graphics environment. Comput. Sci. Eng. 9, 90–95 (2007).

