## Supplemental Information for "Deciphering the evolutionary origin of the enantioselectivity of short-chain dehydrogenases from plants toward 1-borneol"

[1] Institute of Molecular Biotechnology, Graz University of Technology, Petersgasse 14, Graz 8010, Austria. [2] Institute of Chemistry and Biochemistry, Laboratory of Structural Biochemistry, Freie Universität Berlin, Takustraße 6, 14195 Berlin, Germany. [3] School of Chemistry and Biochemistry, Georgia Institute of Technology, 901 Atlantic Drive NW, Atlanta, Georgia 30332-0400, United States. [4] Department of Chemistry - BMC, Uppsala University, BMC Box 576, S-751 23 Uppsala, Sweden. [5] UVA Amsterdam, Department of Chemistry and Pharmaceutical Sciences, Vrije Universiteit Amsterdam, De Boelelaan 1108, 1081 HZ Amsterdam, The Netherlands. [6] BioTechMed-Graz, Mozartgasse 12, 8010 Graz, Austria. [7]Department of Biology, Georgia State University, Atlanta, Georgia, 30303, USA. [8] School of Chemical and Biomolecular Engineering, Georgia Institute of Technology, 311 Ferst Dr., Atlanta, Georgia 30332-0400, United States. [9] Department of Chemistry, Lund University, Box 124, 22100 Lund, Sweden[#]

Equally contributing authors

*Corresponding authors:

Bernhard Loll, Institute of Chemistry and Biochemistry, Laboratory of Structural Biochemistry, Freie Universität Berlin, Takustr. 6, 14195 Berlin, Germany; https://orcid.org/0000-0001-7928-4488;

### Table of contents

Table of contents ..... S3
Supplementary Figures ..... S4
- Sequence logo BDH ..... S4
- Phylogenetic tree of plant BDHs ..... S5
- (-)-borneol conversion of ancestral variants ..... S6
- Addition of DMSO ..... S7
- Apparent Michaelis-Menten kinetics ..... S8
- Apparent E-value ..... S9
- Structural Alignment ..... S10
- N6 AlphaFold prediction ..... S11
- Ancestral Reconstruction Node Probabilities ..... S12
- EVcouplings analysis ..... S12
- Peripheral mutations ..... S13
- Kinetic data of mutant variants ..... S14
- Docking and ML/MM simulations ..... S14
- SDS-PAGE ..... S40
- Structure determination ..... S43

Supplementary Tables ..... S44
- Kinetic values of ancestral mutant variants ..... S44
- Protein yield ..... S44
- Apparent $K_M$-values ..... S45
- Crystallographic data ..... S45
- Docking and ML/MM data ..... S49
- Structural comparisons ..... S49
- Isoborneol converison ..... S51

Supplementary Data ..... S52
- Sequences of ancestral proteins ..... S52
- Equations ..... S53

Supplementary References ..... S55

**Supplementary Figures**

*Sequence logo BDH*

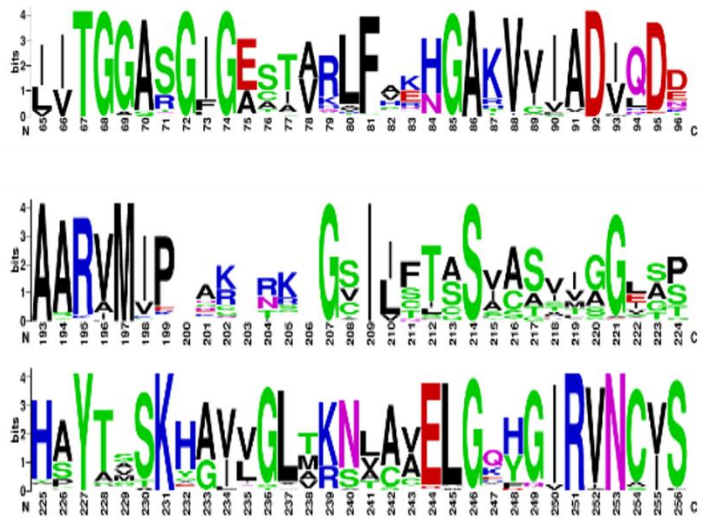

**Supplementary Figure 1**. Graphic representation of the conserved motifs between the extant enzymes. The motif (SxxxxxxxxxxxxYxxxK) can be identified as S214, Y227 and K231 in the consensus sequence. The motif (TGxxx[AG]xG) can be identified between positions 67 and 74.

*Phylogenetic tree of plant BDHs*

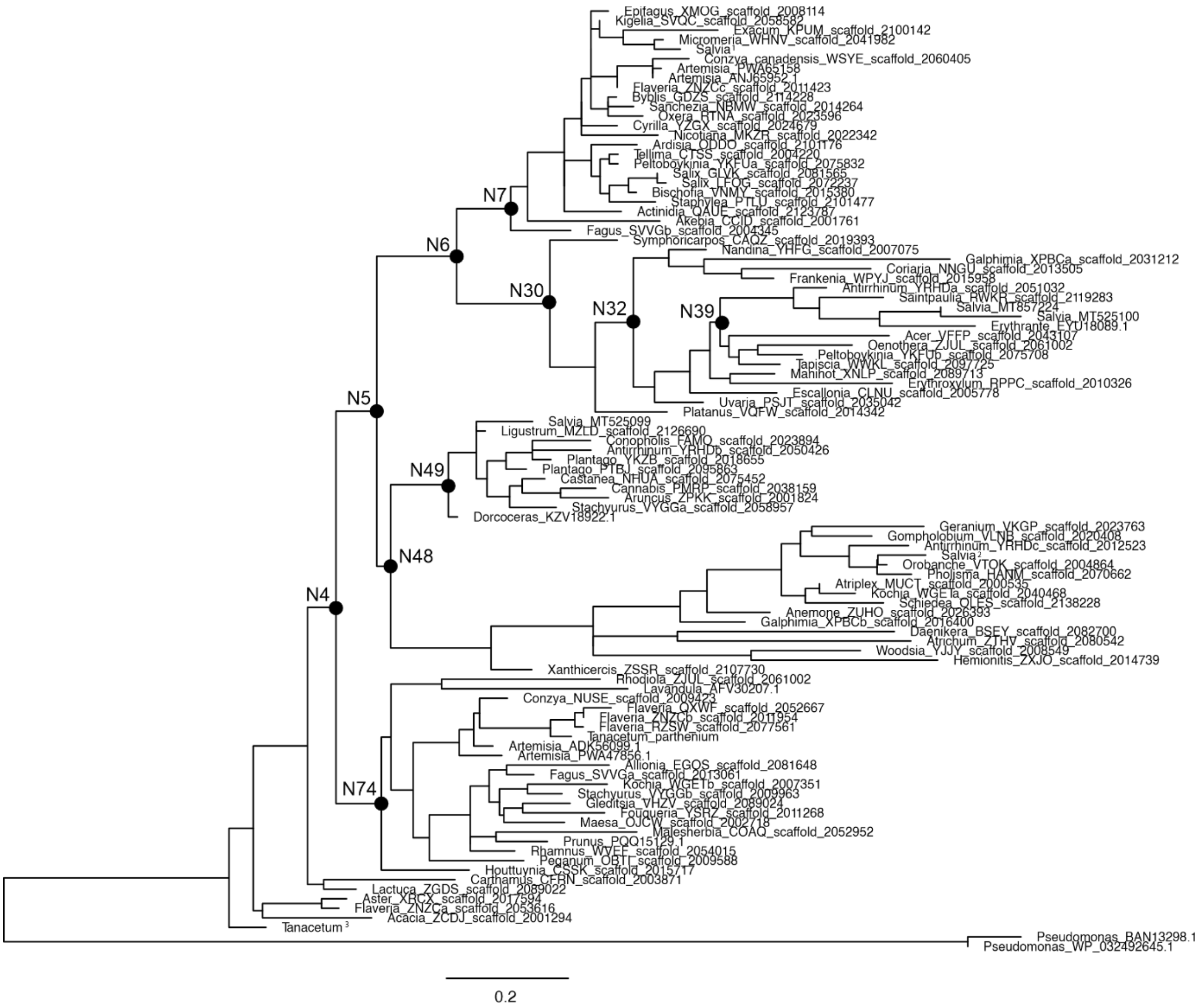


**Supplementary Figure 2**. Phylogenetic tree of plant BDHs. The ten highlighted ancestors were characterized computationally or biochemically, or both. *Salvia* BDH sequences were reported by Chánique et. al. (2021), *Tanacetum* BDHs are reported under https://db.cngb.org/onekp/species/Tanacetum%20parthenium; sample code = DUQG.

*(-)-borneol conversion of ancestral variants*

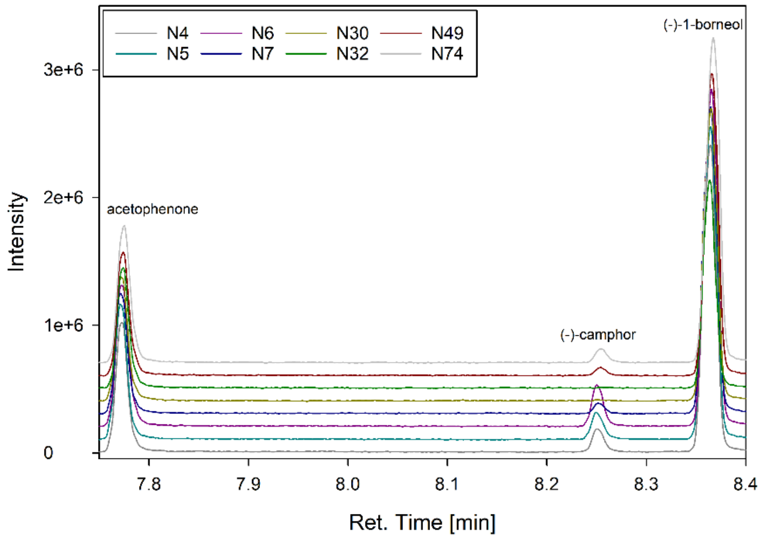


**Supplementary Figure 3.** Chromatogram of endpoint measurements (1.5 h) of (−)-1-borneol conversion with BDH ancestors N4, N5, N6, N7, N30, N32, N49 and N74 measured on GC-MS.

*Addition of DMSO*

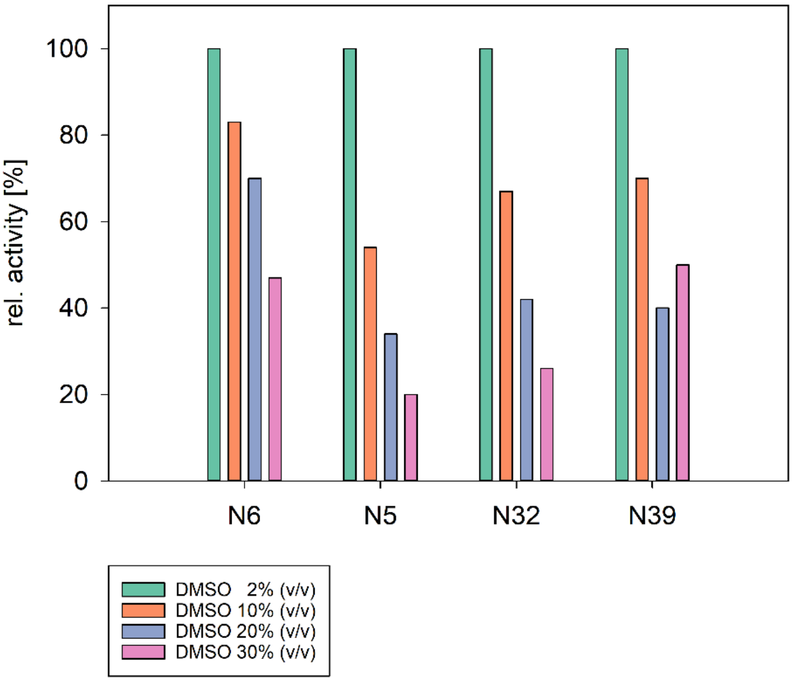


**Supplementary Figure 4.** DMSO tolerance measurement with (+)-1-borneol of ancestors N6, N5, N32, and N39 at 2%, 10%, 20% and 30% (v/v) DMSO.

*Apparent Michaelis-Menten kinetics*

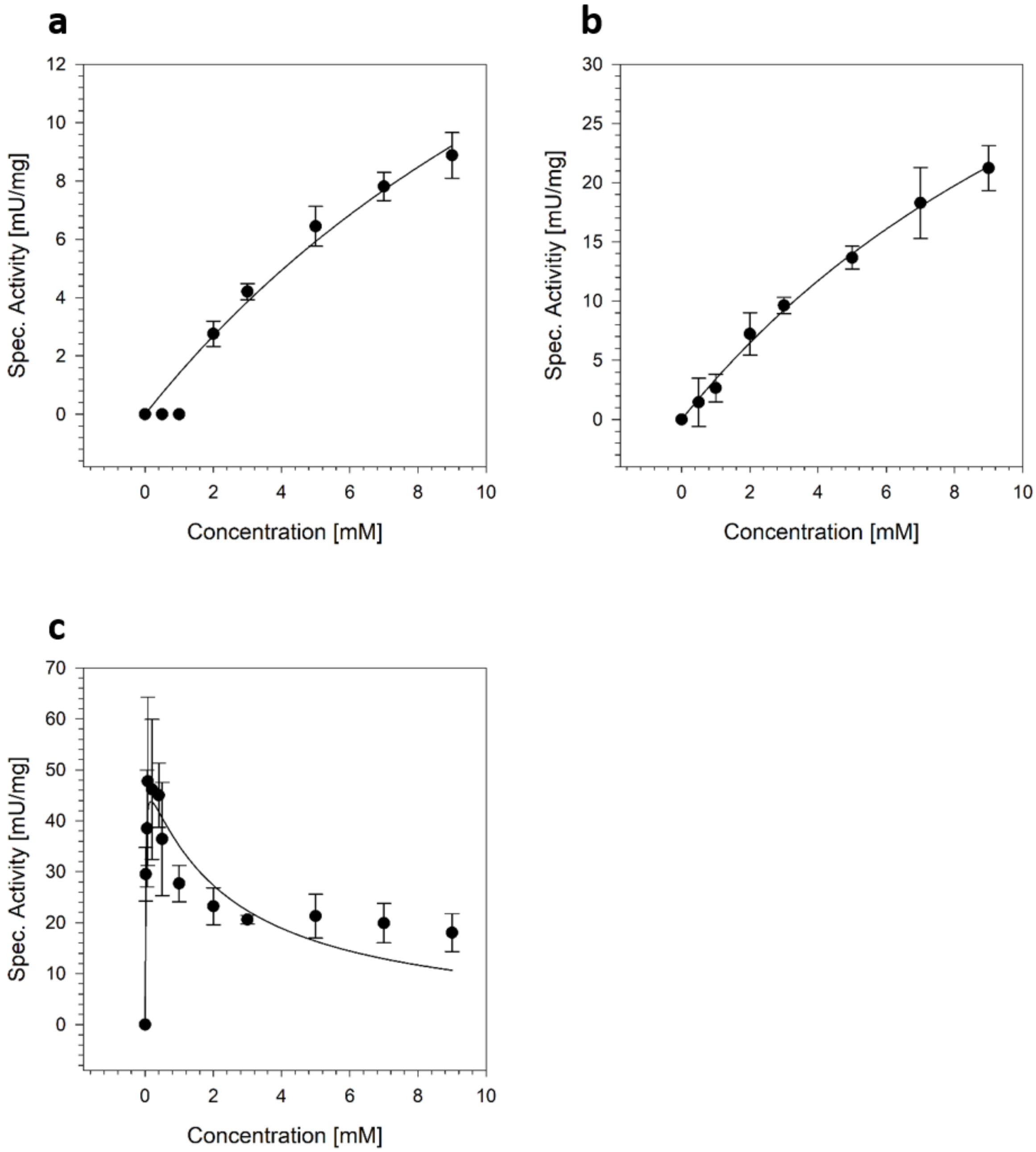


**Supplementary Figure 5.** Specific activities of the ancestors plotted against (+)-1-borneol concentration and Michaelis Menten regression visualized in Sigma Plot. Kinetic data represent variants: (A) N30, (B) N32, (C) N6. Error bars represent the standard deviation of measurements carried out in quintuplicates. Raw data of the measurement is deposited under 10.5281/zenodo.18925996 (Raw data kinetics_N6_N30_N32.xlsx).

*Apparent E-value*

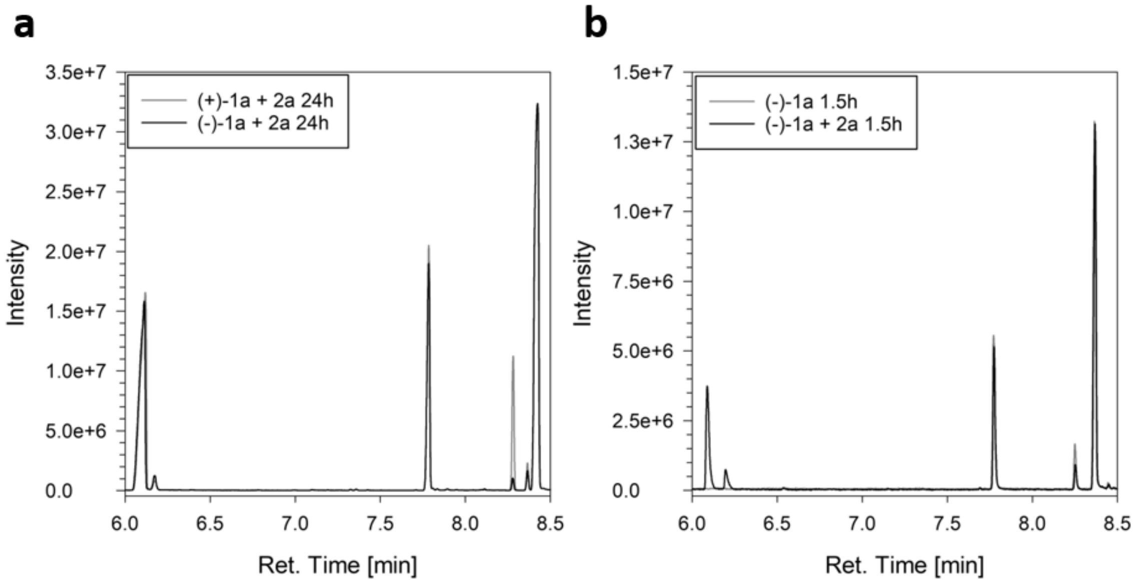


**Supplementary Figure 6.** GC-MS chromatograms highlighting the conversion of (+)-1-borneol ((+)-1a), (−)-1-borneol ((−)-1a) in comparison to the surrogate substrate cyclohexanol (2a). Ret. times: cyclohexanol 6.06 min, cyclohexanone 6.17 min, acetophenone 7.79 min, camphor 8.29 min, isoborneol 8.35 min, and borneol 8.41 min. (A) Ancestor N6 under competitive conditions, (B) N6 under (non-)competitive conditions.

*Structural Alignment*

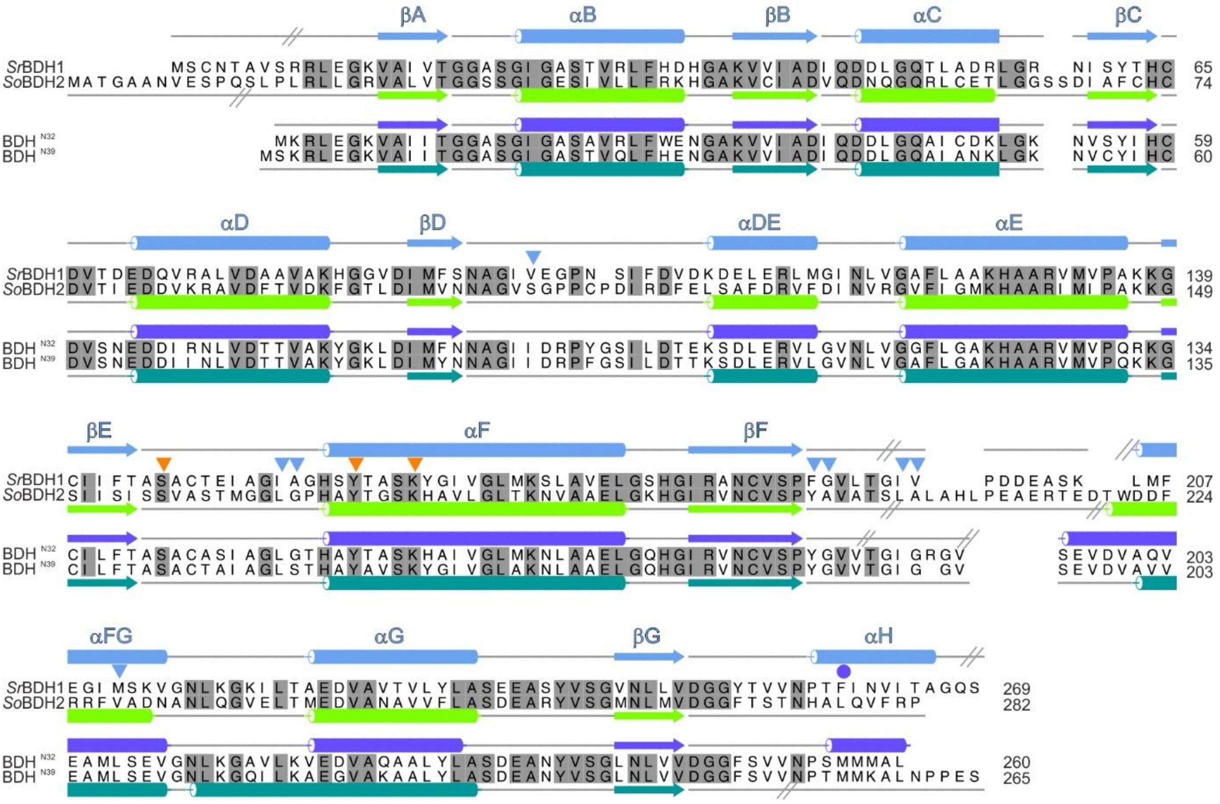


**Supplementary Figure 7.** Structure-based sequence alignment of *Sr*BDH1 (GeneBank ID MT857224) *So*BDH2 (GeneBank ID MT525099), N32, and N39. Secondary structure elements are drawn above the alignment for *Sr*BDH1 and below for *So*BDH2 or N32 and below for N39 with α-helices depicted as cylinders and β-strands as arrows. Grey, inclined lines indicated sections of the structures, since they were not resolved in the reconstruction and could not be modelled. Orange triangles indicate the catalytic motif. Amino acids lining the putative active site of *Sr*BDH1, based on its crystal structure (PDB ID: 6ZYZ) with bound $NAD^+$, are indicated by blue triangles and as dark blue triangle if derived from another *Sr*BDH1 monomer within the tetramer. The dark green triangle marks a residue derived from another protomer of *So*BDH2 that completes the active site. Grey shaded amino acids are identical. The TGxxx[AG]xG $NAD^+$ binding motif, between βA and αB is indicated with a magenta rectangle.

*N6 AlphaFold prediction*

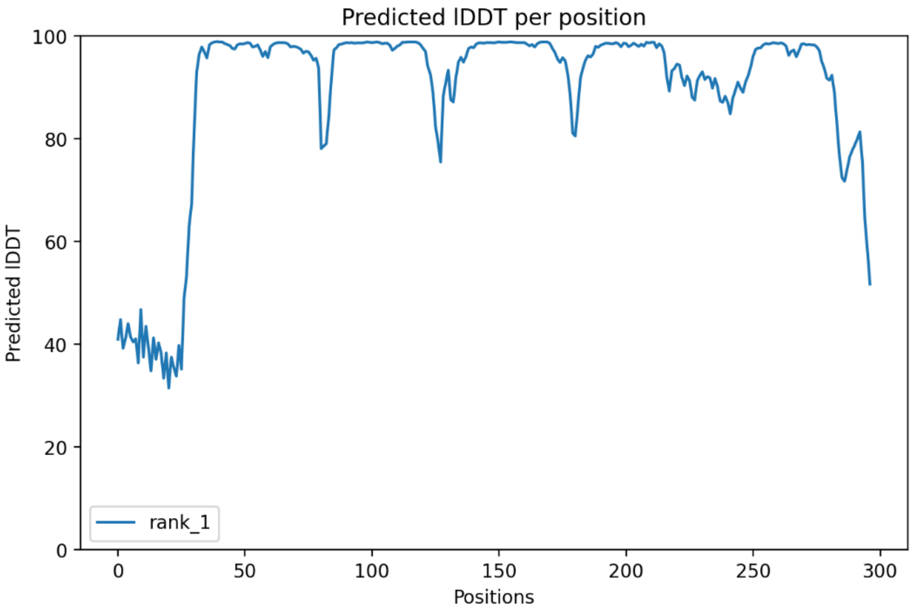


**Supplementary Figure 8.** pLDDT score distribution of the model of N6 created with AlphaFold3.

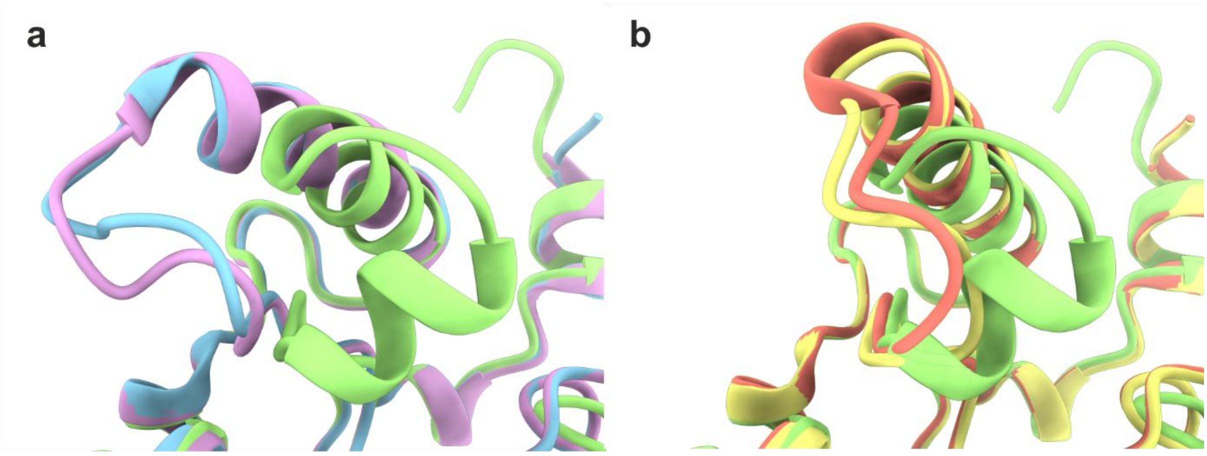


**Supplementary Figure 9**. Comparison of the loop-αFG motif structure in the N6 AlphaFold3 model and in the 4 chains of *Sr*BDH1 (PDB: 6ZYZ). a Chains A (blue) and B (lilac) show an open conformation of the loop, differing from the predicted N6 model. b Chains C (yellow) and D (salmon), while not completely aligning with the predicted model, show a closed conformation, closer to N6.

*Ancestral Reconstruction Node Probabilities*

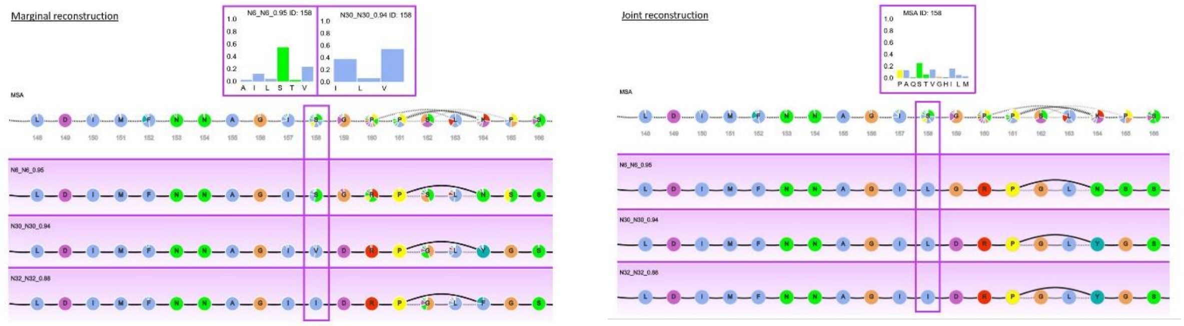


**Supplementary Figure 10**. Comparison of marginal and joint sequence reconstruction conducted on the GRASP webserver. Statistics indicate the posterior probability of the residue corresponding to amino acid position 111 in N32 and N30. Visualization based on the GRASP web interface. Purple boxes highlight the reconstructed sequence of the ancestral nodes at the indicated positions. Dashed lines indicate the probable occurrence of gaps in the sequence.

*EVcouplings analysis*

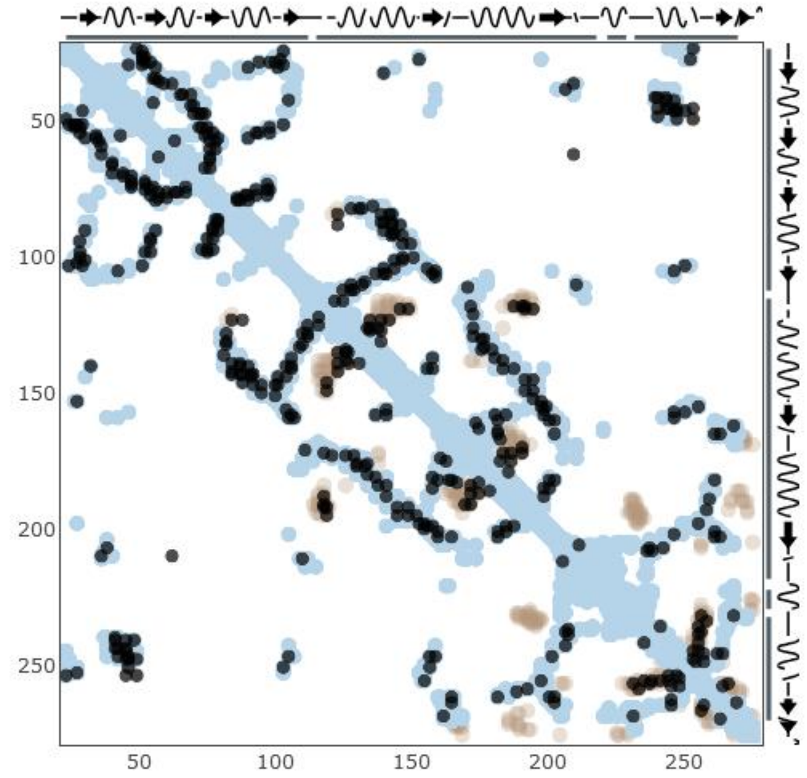


**Supplementary Figure 11.** Distance map computed by EVcouplings based on N32. Black dots indicate evolutionary couples, light blue dots monomer contacts, and brown dots multimer contacts.

*Peripheral mutations*

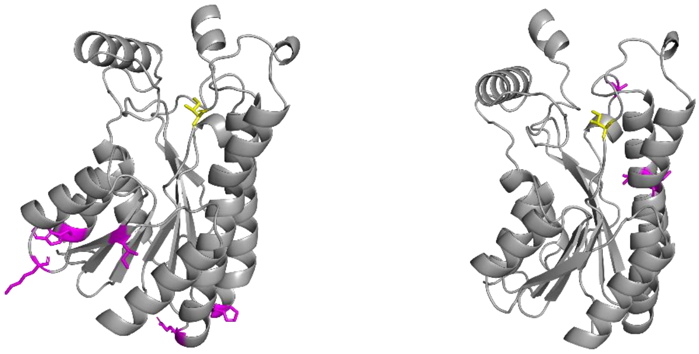

**Supplementary Figure 12**. AlphaFold3 structure with highlighted mutations in the (left) N30_L111I/L47W/E74K/V79I/E107A/E153Q/Y197H variant and in the (right) N30_L111I/V136L/G169A/V183I variant. Yellow indicates position 111, purple indicates additional mutations.

*Kinetic data of mutant variants*

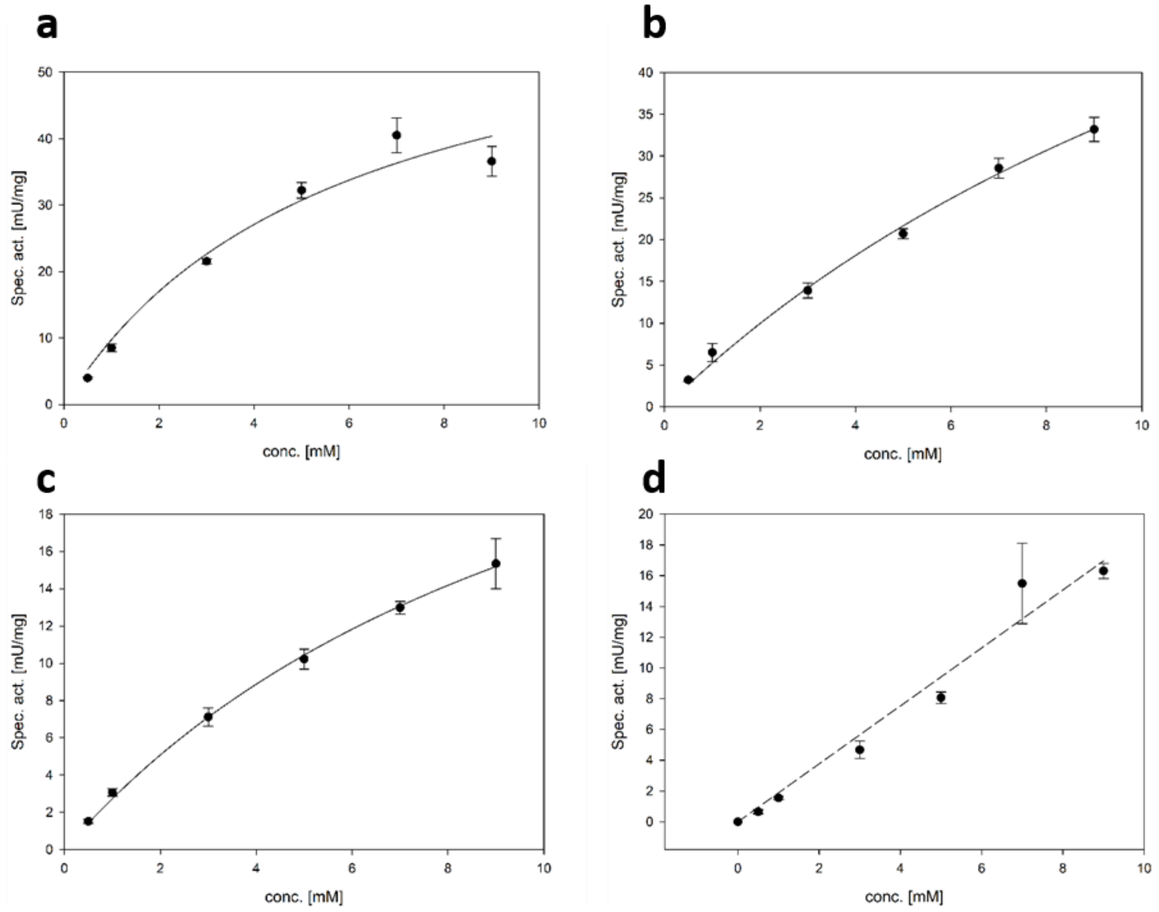


**Supplementary Figure 13.** Kinetic data fitted to the Michaelis-Menten model of (A) N32_I111L, (B) N30_ L111I and (C) N30_ L111I/G169A/V136L/V183I, and N30_L111I/V136L/G169A/V183I (D). Varying substrate conc. of (+)-borneol, 20% DMSO, 9 mM NAD+, 20 µM purified enzyme, 0.1 M Tris-HCl pH = 9. Measured on GC-FID equipped with a ZB-5 column. (n=3). Raw data and GC-chromatograms of the measurement are deposited under 10.5281/zenodo.18925996 (Chromatogramms.zip; N32_I111L_kin.xlsx; N30_L111IG169AV136LV183I_kin.xlsx; and N30_L111I_kin.xlsx;).

*Docking and ML/MM simulations*

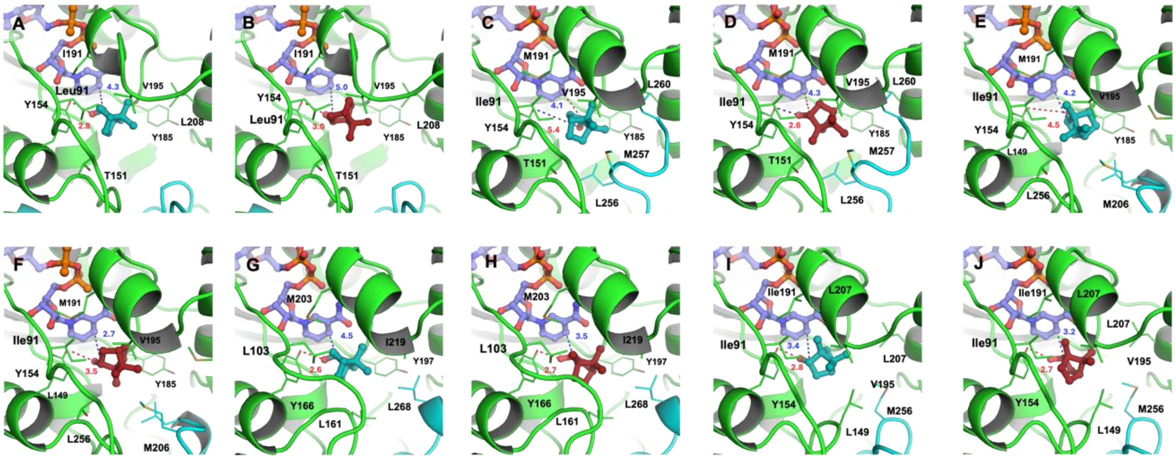

**Supplementary Figure 14.** Selected docking poses for the complexes between: N32_I111L (A-B); N30_L111I (C-D); N30_L111I/G169A/V136L/V183I (E-F); N6 (G-H); N32 (I-J) with (+)-borneol is and (−)-borneol showed in teal and firebrick color. Relevant amino acids in the binding are represented in sticks as well as $NAD^+$ colored in slate.

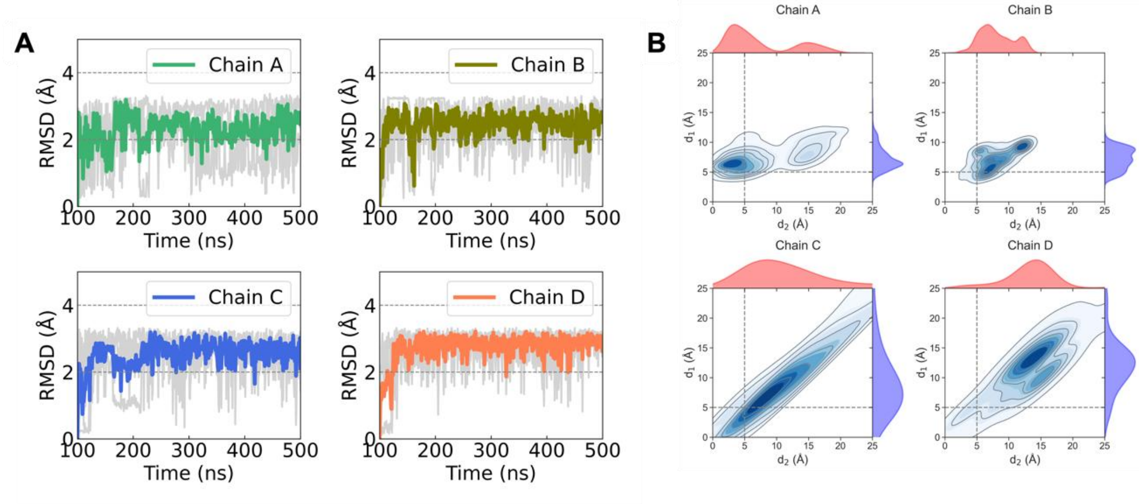


**Supplementary Figure 15.** (A) RMSD values (Å) from 3x400 ns MD simulations of *Sr*BDH1 complex with (+)-borneol. Individual replica trajectories are shown in gray, while the colored trace represents the mean RMSD (Å) averaged over the replicas. (B) Distributions of the two monitored catalytic distances (Å) across the simulation replicas; $d_1$ corresponds to the hydride transfer geometry between the C4 atom of the nicotinamide ring of $NAD^+$ and the oxidizable C5 carbon of borneol, and $d_2$ corresponds to the hydrogen-bond interaction between the borneol hydroxyl oxygen and the catalytic oxygen atom of the conserved tyrosine side chain.

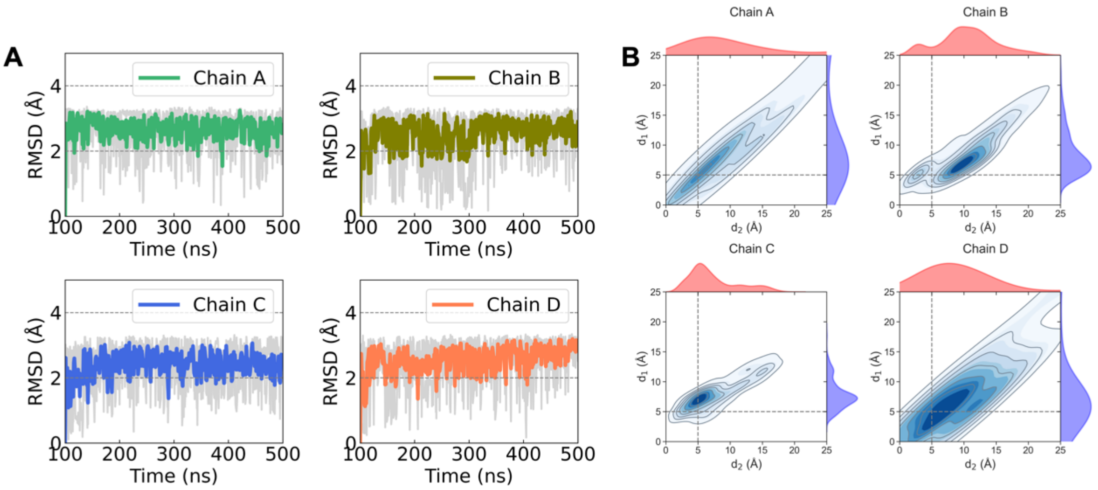


**Supplementary Figure 16**. (A) RMSD values (Å) from 3x400 ns MD simulations of *Sr*BDH1 complex with (−)-borneol. Individual replica trajectories are shown in gray, while the colored trace represents the mean RMSD (Å) averaged over the replicas. (B) Distributions of the two monitored catalytic distances (Å) across the simulation replicas; $d_1$ corresponds to the hydride transfer geometry between the C4 atom of the nicotinamide ring of $NAD^+$ and the oxidizable C5 carbon of borneol, and $d_2$ corresponds to the hydrogen-bond interaction between the borneol hydroxyl oxygen and the catalytic oxygen atom of the conserved tyrosine side chain.

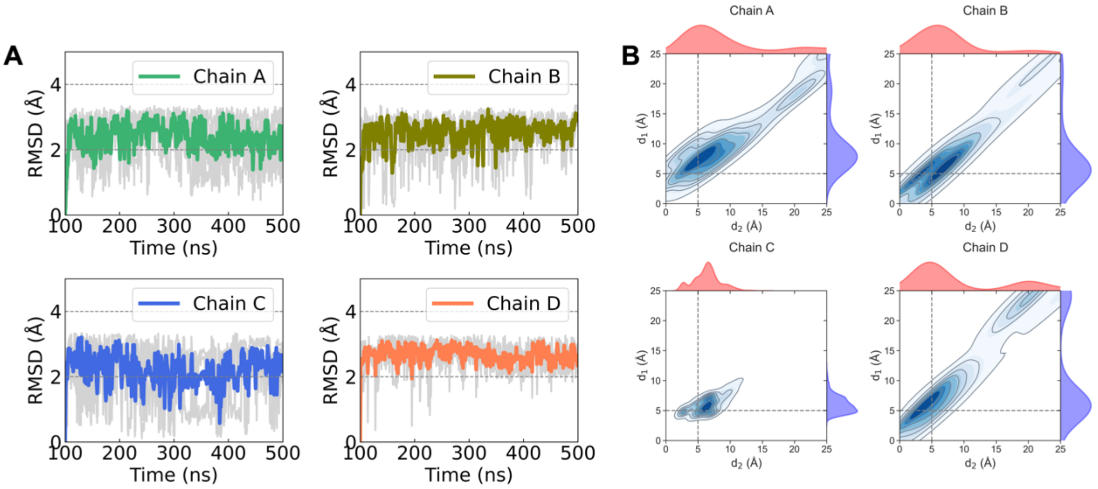


**Supplementary Figure 17**. (A) RMSD values (Å) from 3x400 ns MD simulations of *Sr*BDH1 N32_I111L ancestor complex with (+)-borneol. Individual replica trajectories are shown in gray, while the colored trace represents the mean RMSD (Å) averaged over the replicas. (B)

Distributions of the two monitored catalytic distances (Å) across the simulation replicas; $d_1$ corresponds to the hydride transfer geometry between the C4 atom of the nicotinamide ring of $NAD^+$ and the oxidizable C5 carbon of borneol, and $d_2$ corresponds to the hydrogen-bond interaction between the borneol hydroxyl oxygen and the catalytic oxygen atom of the conserved tyrosine side chain.

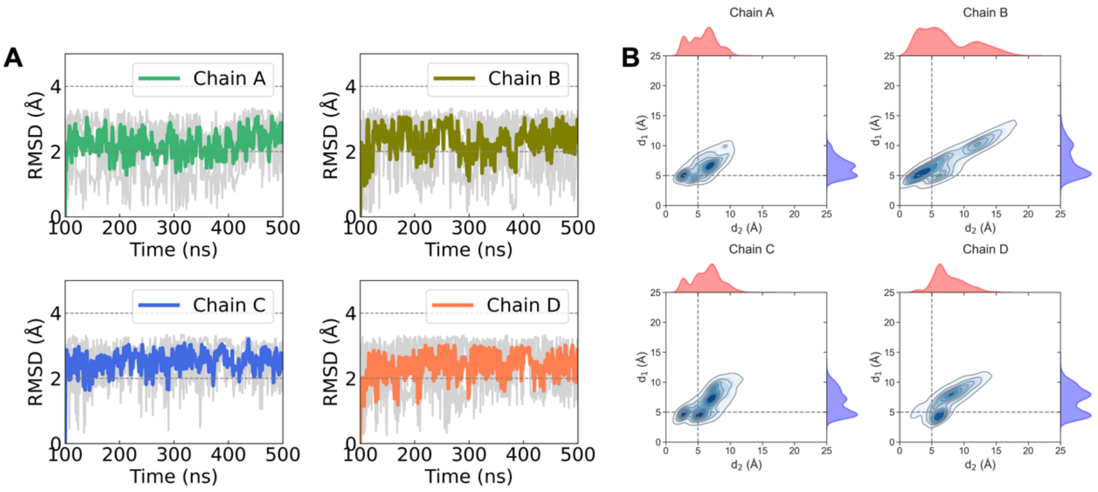


**Supplementary Figure 18**. (A) RMSD values (Å) from 3x400 ns MD simulations of *Sr*BDH1 N32_I111L ancestor complex with (−)-borneol. Individual replica trajectories are shown in gray, while the colored trace represents the mean RMSD (Å) averaged over the replicas. (B) Distributions of the two monitored catalytic distances (Å) across the three simulation replicas; $d_1$ corresponds to the hydride transfer geometry between the C4 atom of the nicotinamide ring of $NAD^+$ and the oxidizable C5 carbon of borneol, and $d_2$ corresponds to the hydrogen-bond interaction between the borneol hydroxyl oxygen and the catalytic oxygen atom of the conserved tyrosine side chain.

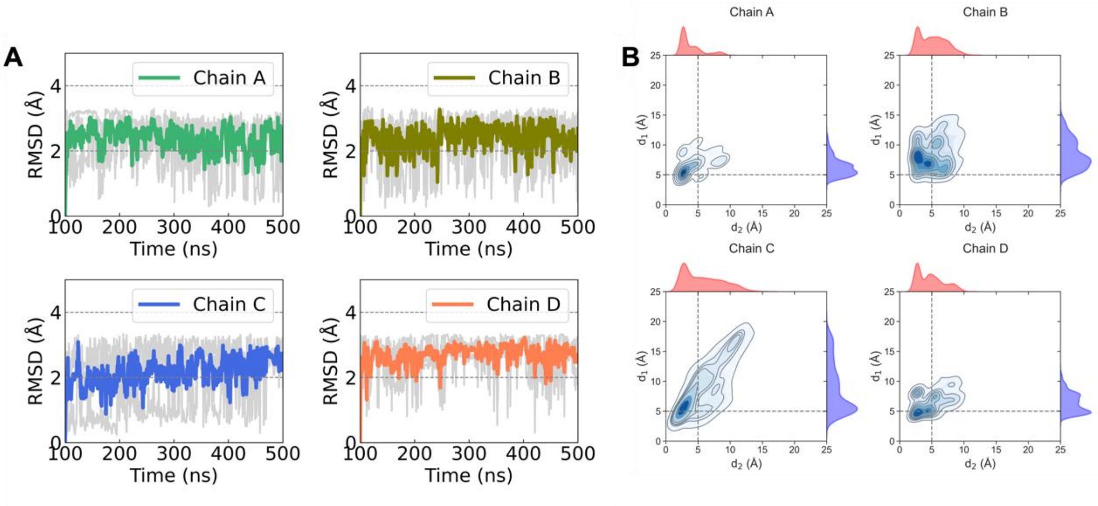


**Supplementary Figure 19**. (A) RMSD values (Å) from 3x400 ns MD simulations of *Sr*BDH1 N30_L111I ancestor complex with (+)-borneol. Individual replica trajectories are shown in gray, while the colored trace represents the mean RMSD (Å) averaged over the replicas. (B) Distributions of the two monitored catalytic distances (Å) across the simulation replicas; $d_1$ corresponds to the hydride transfer geometry between the C4 atom of the nicotinamide ring of $NAD^+$ and the oxidizable C5 carbon of borneol, and $d_2$ corresponds to the hydrogen-bond interaction between the borneol hydroxyl oxygen and the catalytic oxygen atom of the conserved tyrosine side chain.

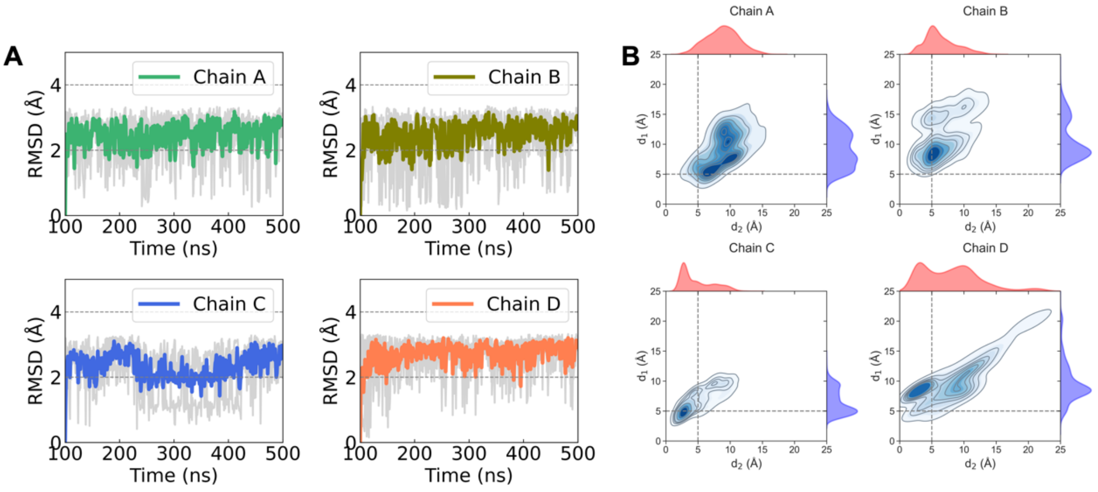


**Supplementary Figure 20**. (A) RMSD values (Å) from 3x400 ns MD simulations of *Sr*BDH1 N30_L111I ancestor complex with (−)-borneol. Individual replica trajectories are shown in gray, while the colored trace represents the mean RMSD (Å) averaged over the replicas. (B) Distributions of the two monitored catalytic distances (Å) across the simulation replicas; $d_1$ corresponds to the hydride transfer geometry between the C4 atom of the nicotinamide ring of $NAD^+$ and the oxidizable C5 carbon of borneol, and $d_2$ corresponds to the hydrogen-bond interaction between the borneol hydroxyl oxygen and the catalytic oxygen atom of the conserved tyrosine side chain.

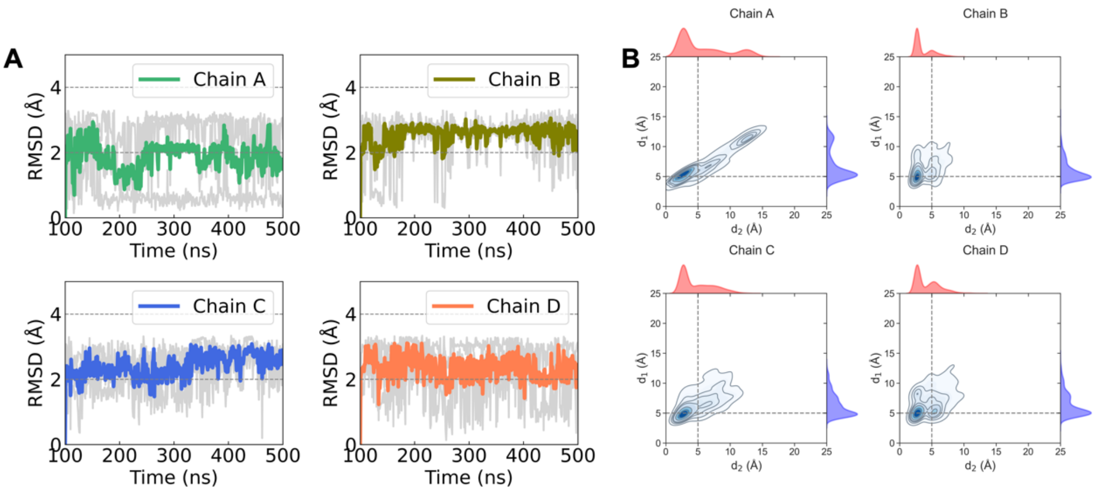


**Supplementary Figure 21**. (A) RMSD values (Å) from 3x400 ns MD simulations of *Sr*BDH1 N30_L111I/G169A/V136L/V183I ancestor complex with (+)-borneol. Individual replica trajectories are shown in gray, while the colored trace represents the mean RMSD (Å) averaged over the replicas. (B) Distributions of the two monitored catalytic distances (Å) across the simulation replicas; $d_1$ corresponds to the hydride transfer geometry between the C4 atom of the nicotinamide ring of $NAD^+$ and the oxidizable C5 carbon of borneol, and $d_2$ corresponds to the hydrogen-bond interaction between the borneol hydroxyl oxygen and the catalytic oxygen atom of the conserved tyrosine side chain.

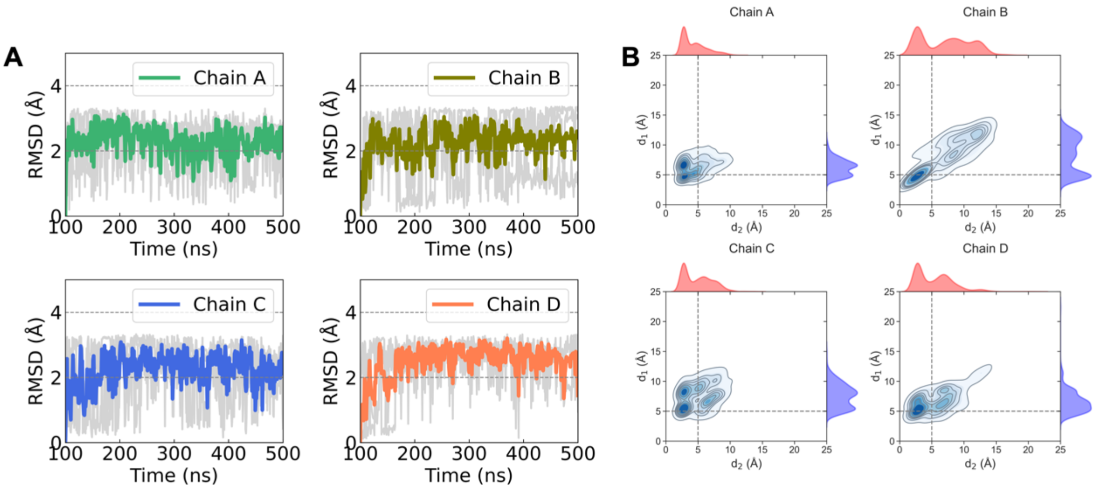


**Supplementary Figure 22**. (A) RMSD values (Å) from 3x400 ns MD simulations of *Sr*BDH1 N30_L111I/G169A/V136L/V183I ancestor complex with (−)-borneol. Individual replica trajectories are shown in gray, while the colored trace represents the mean RMSD (Å) averaged over the replicas. (B) Distributions of the two monitored catalytic distances (Å) across the simulation replicas; $d_1$ corresponds to the hydride transfer geometry between the C4 atom of the nicotinamide ring of $NAD^+$ and the oxidizable C5 carbon of borneol, and $d_2$ corresponds to the hydrogen-bond interaction between the borneol hydroxyl oxygen and the catalytic oxygen atom of the conserved tyrosine side chain.

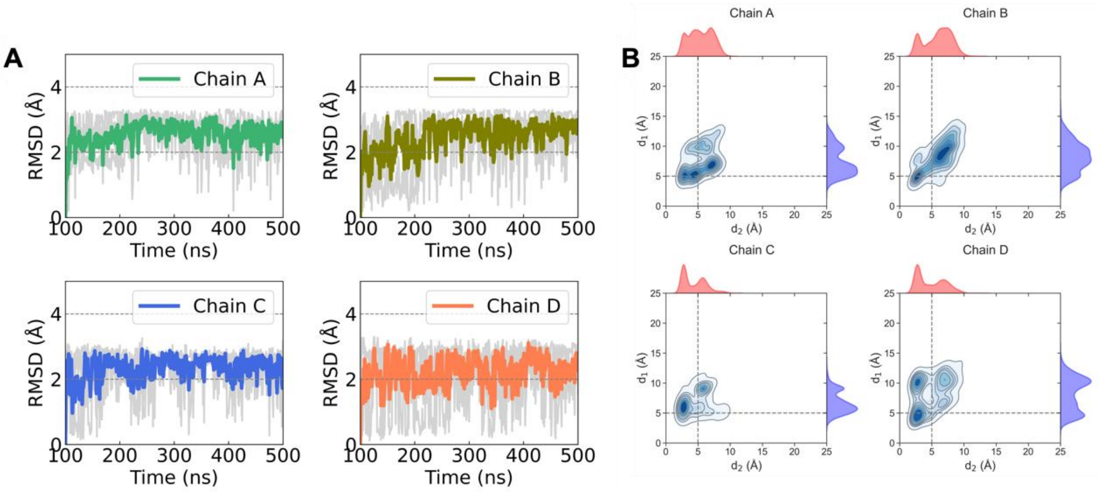


**Supplementary Figure 23**. (A) RMSD values (Å) from 3x400 ns MD simulations of *Sr*BDH1 N6 ancestor complex with (+)-borneol. Individual replica trajectories are shown in gray, while the colored trace represents the mean RMSD (Å) averaged over the replicas. (B) Distributions of the two monitored catalytic distances (Å) across the simulation replicas; $d_1$ corresponds to the hydride transfer geometry between the C4 atom of the nicotinamide ring of $NAD^+$ and the oxidizable C5 carbon of borneol, and $d_2$ corresponds to the hydrogen-bond interaction between the borneol hydroxyl oxygen and the catalytic oxygen atom of the conserved tyrosine side chain.

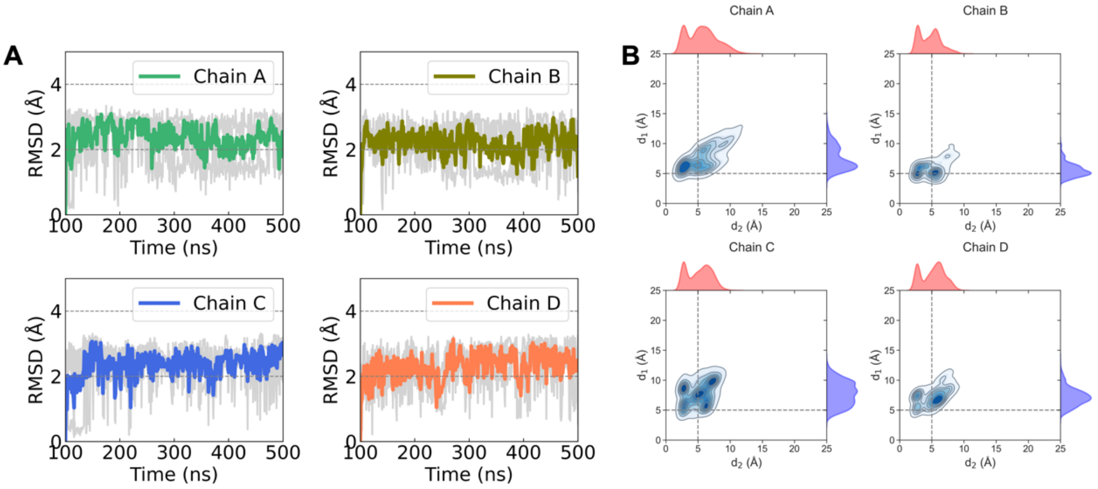


**Supplementary Figure 24**. (A) RMSD values (Å) from 3x400 ns MD simulations of *Sr*BDH1 N6 ancestor complex with (−)-borneol. Individual replica trajectories are shown in gray, while the colored trace represents the mean RMSD (Å) averaged over the replicas. (B) Distributions of the two monitored catalytic distances (Å) across the simulation replicas; $d_1$ corresponds to the hydride transfer geometry between the C4 atom of the nicotinamide ring of $NAD^+$ and the oxidizable C5 carbon of borneol, and $d_2$ corresponds to the hydrogen-bond interaction between the borneol hydroxyl oxygen and the catalytic oxygen atom of the conserved tyrosine side chain.

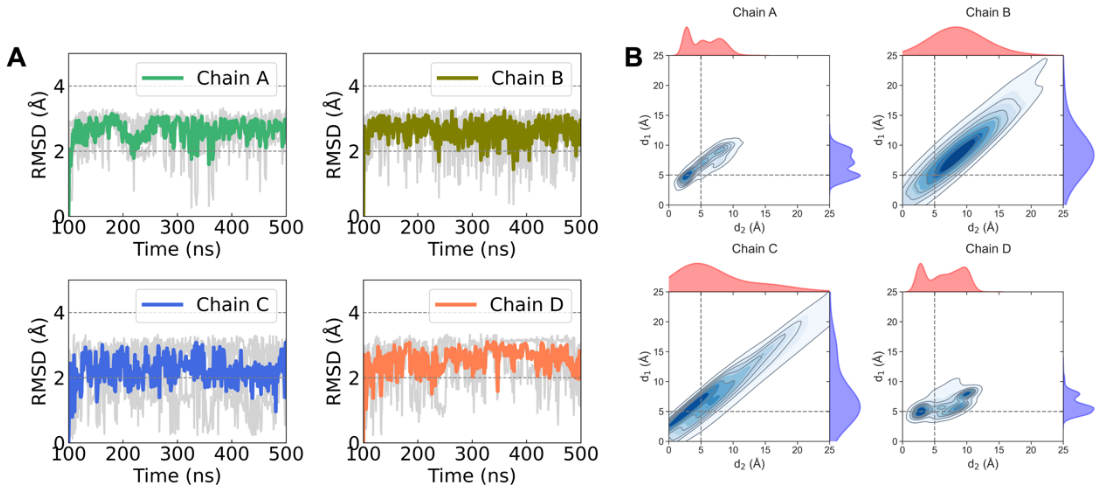


**Supplementary Figure 25**. (A) RMSD values (Å) from 3x400 ns MD simulations of *Sr*BDH1 N32 ancestor complex with (+)-borneol. Individual replica trajectories are shown in gray, while the colored trace represents the mean RMSD (Å) averaged over the replicas. (B) Distributions of the two monitored catalytic distances (Å) across the simulation replicas; $d_1$ corresponds to the hydride transfer geometry between the C4 atom of the nicotinamide ring of $NAD^+$ and the oxidizable C5 carbon of borneol, and $d_2$ corresponds to the hydrogen-bond interaction between the borneol hydroxyl oxygen and the catalytic oxygen atom of the conserved tyrosine side chain.

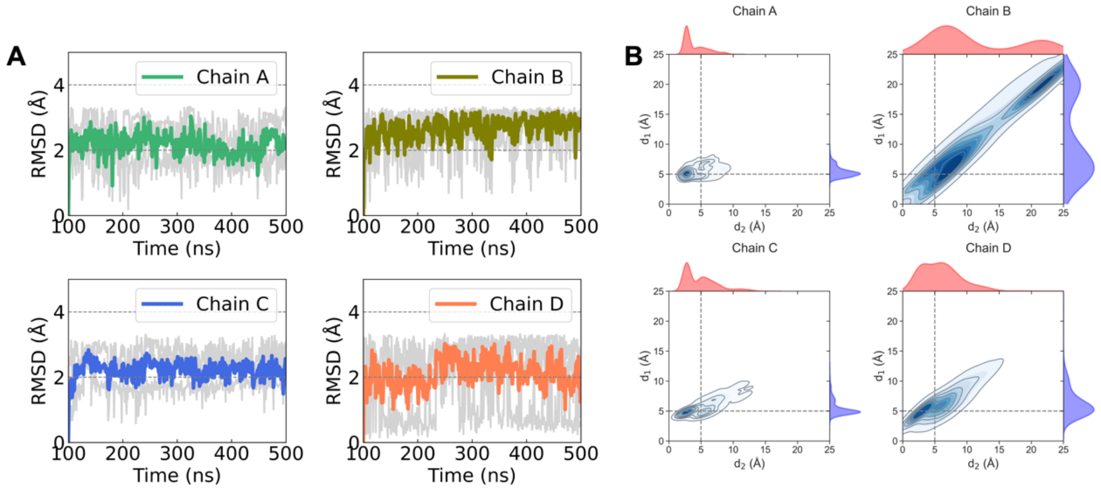


**Supplementary Figure 26**. (A) RMSD values (Å) from 3x400 ns MD simulations of *Sr*BDH1 N32 ancestor complex with (−)-borneol. Individual replica trajectories are shown in gray, while the colored trace represents the mean RMSD (Å) averaged over the replicas. (B) Distributions of the two monitored catalytic distances (Å) across the simulation replicas; $d_1$ corresponds to the hydride transfer geometry between the C4 atom of the nicotinamide ring of $NAD^+$ and the oxidizable C5 carbon of borneol, and $d_2$ corresponds to the hydrogen-bond interaction between the borneol hydroxyl oxygen and the catalytic oxygen atom of the conserved tyrosine side chain.

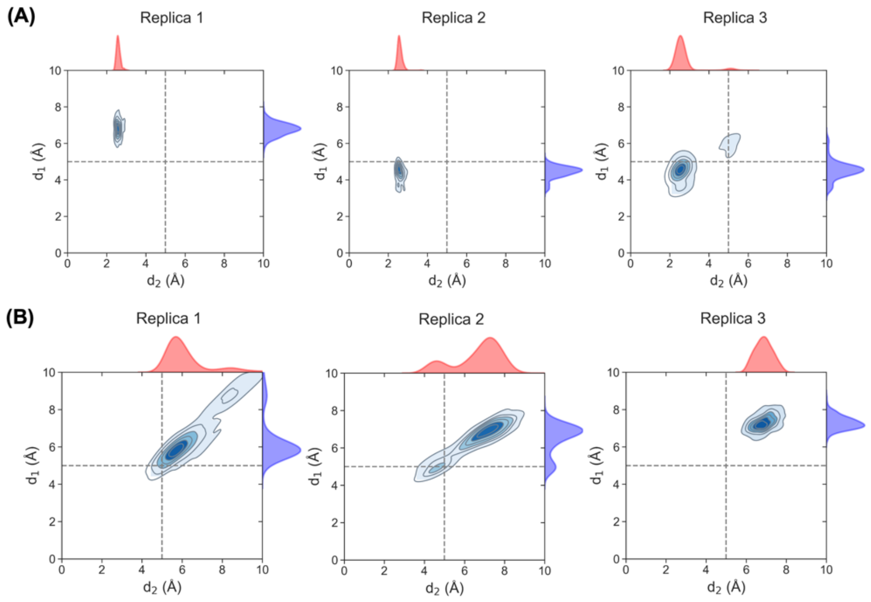


**Supplementary Figure 27**. Kernel density estimation (KDE) of the catalytic distance distributions obtained from 10 ns ML/MM simulations of the *Sr*BDH1 N32_I111L variant in complex with (+)- and (−)-borneol. Distance $d_1$ (blue) corresponds to the hydride-transfer distance, while $d_2$ (red) represents the interaction between the catalytic tyrosine and the substrate. Three independent replicas were used for sampling.

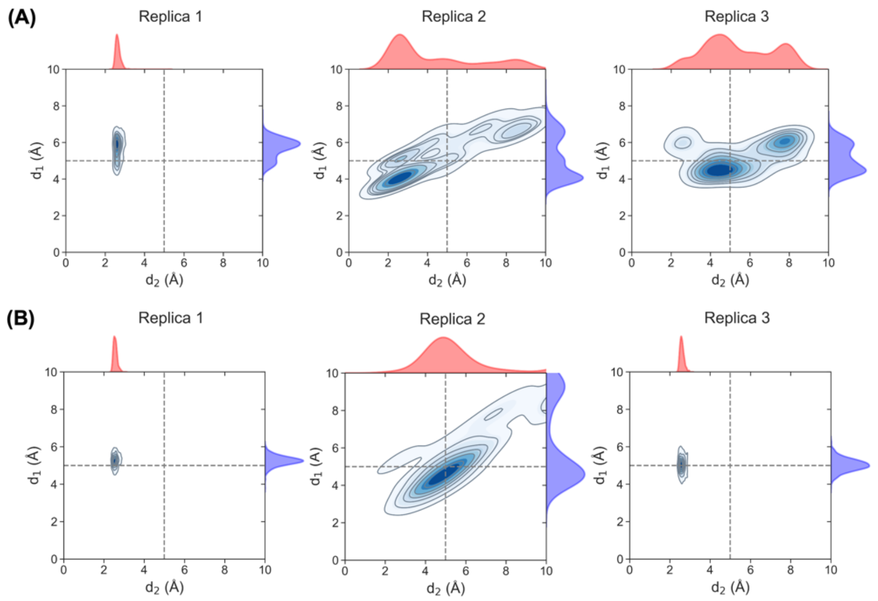


**Supplementary Figure 28**. Kernel density estimation (KDE) of the catalytic distance distributions obtained from 10 ns ML/MM simulations of the *Sr*BDH1 N32_I111L variant in complex with (+)- and (−)-borneol. Distance $d_1$ (blue) corresponds to the hydride-transfer distance, while $d_2$ (red) represents the interaction between the catalytic tyrosine and the substrate. Three independent replicas were used for sampling.

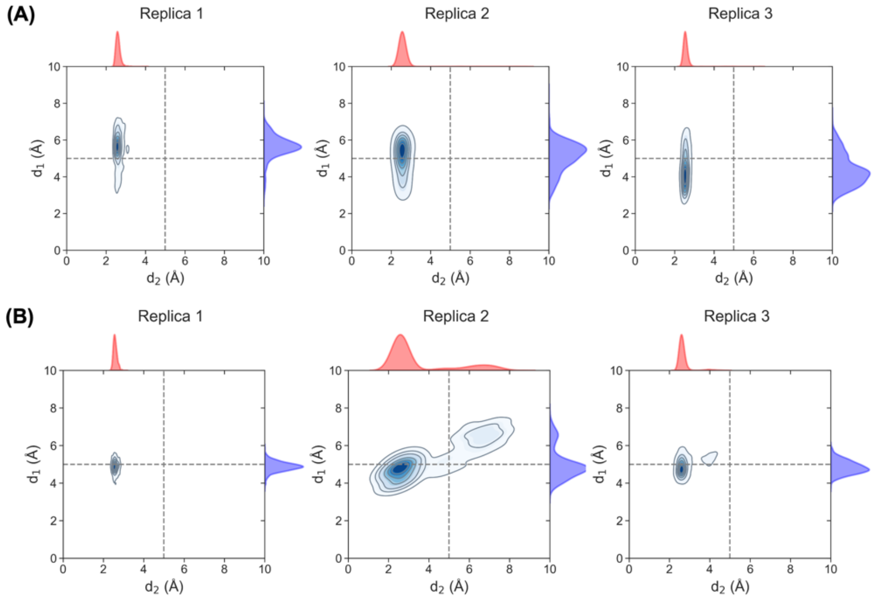


**Supplementary Figure 29**. Kernel density estimation (KDE) of the catalytic distance distributions obtained from 10 ns ML/MM simulations of the *Sr*BDH1 N30_L111I variant in complex with (+)- and (−)-borneol. Distance $d_1$ (blue) corresponds to the hydride-transfer distance, while $d_2$ (red) represents the interaction between the catalytic tyrosine and the substrate. Three independent replicas were used for sampling.

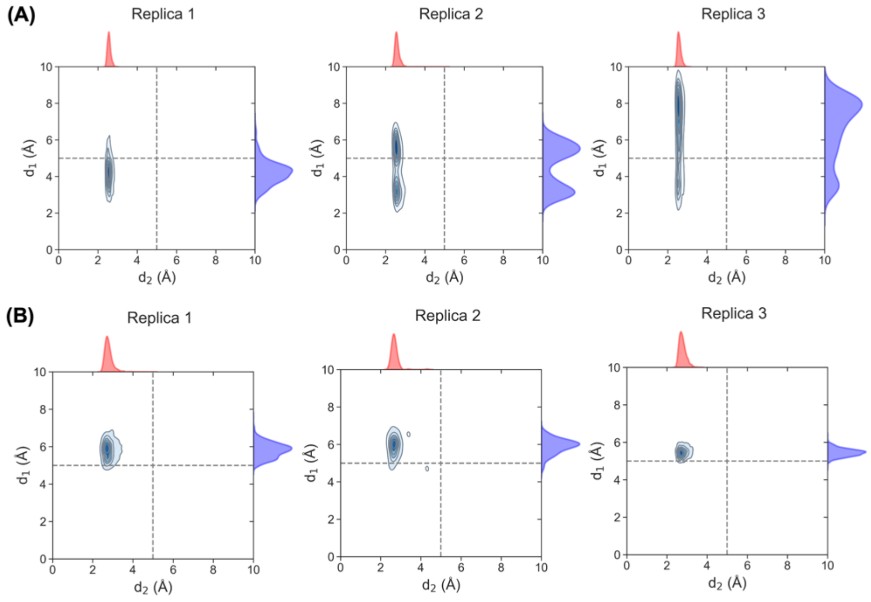


**Supplementary Figure 30**. Kernel density estimation (KDE) of the catalytic distance distributions obtained from 10 ns ML/MM simulations of the *Sr*BDH1 N30_L111I/G169A/V136L/V183I variant in complex with (+)- and (−)-borneol. Distance $d_1$ (blue) corresponds to the hydride-transfer distance, while $d_2$ (red) represents the interaction between the catalytic tyrosine and the substrate. Three independent replicas were used for sampling.

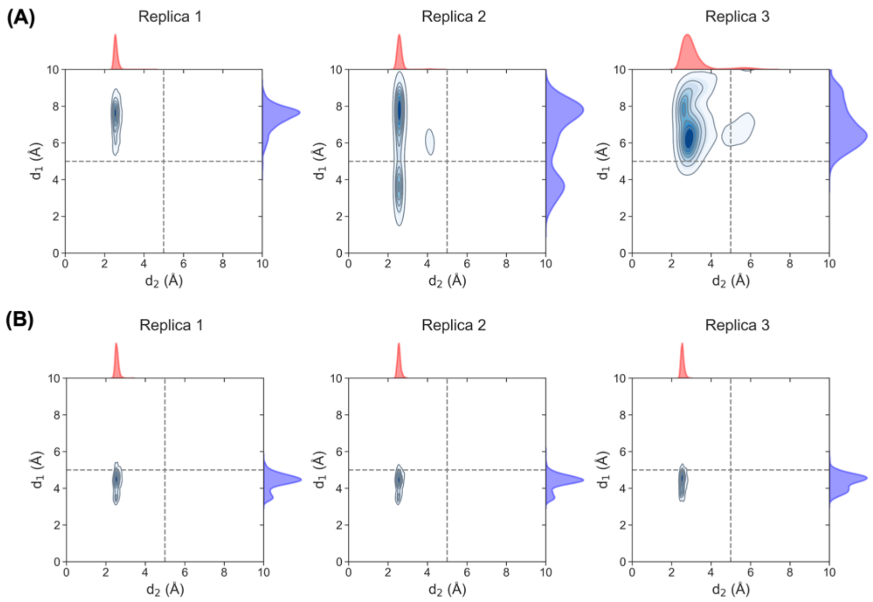


**Supplementary Figure 31**. Kernel density estimation (KDE) of the catalytic distance distributions obtained from 10 ns ML/MM simulations of the *Sr*BDH1 N6 variant in complex with (+)- and (−)-borneol. Distance $d_1$ (blue) corresponds to the hydride-transfer distance, while $d_2$ (red) represents the interaction between the catalytic tyrosine and the substrate. Three independent replicas were used for sampling.

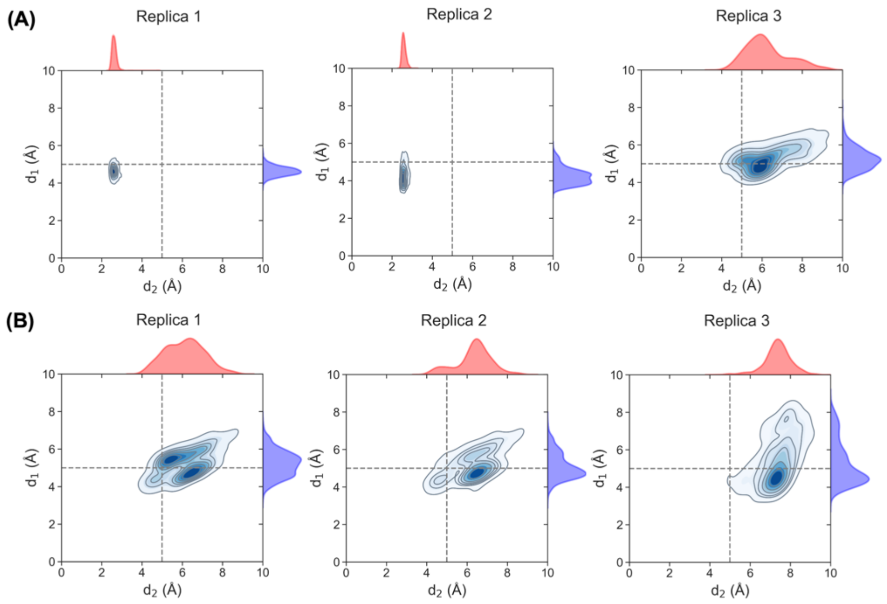


**Supplementary Figure 32**. Kernel density estimation (KDE) of the catalytic distance distributions obtained from 10 ns ML/MM simulations of the *Sr*BDH1 N32 variant in complex with (+)- and (−)-borneol. Distance $d_1$ (blue) corresponds to the hydride-transfer distance, while $d_2$ (red) represents the interaction between the catalytic tyrosine and the substrate. Three independent replicas were used for sampling.

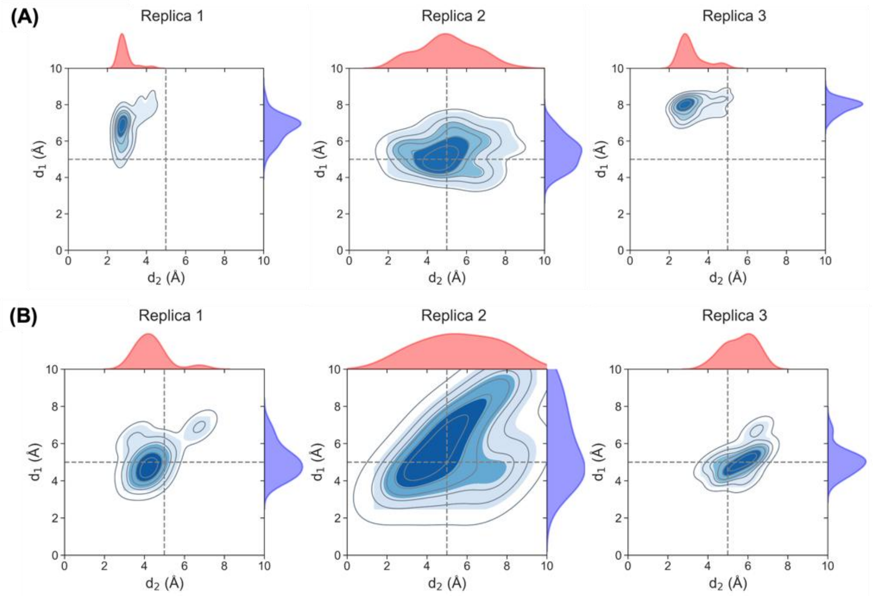


**Supplementary Figure 33**. Kernel density estimation (KDE) of the catalytic distance distributions obtained from 10 ns conventional MD simulations of the *Sr*BDH1 wild-type in complex with (+)- and (−)-borneol. Distance $d_1$ (blue) corresponds to the hydride-transfer distance, while $d_2$ (red) represents the interaction between the catalytic tyrosine and the substrate. Three independent replicas were used for sampling.

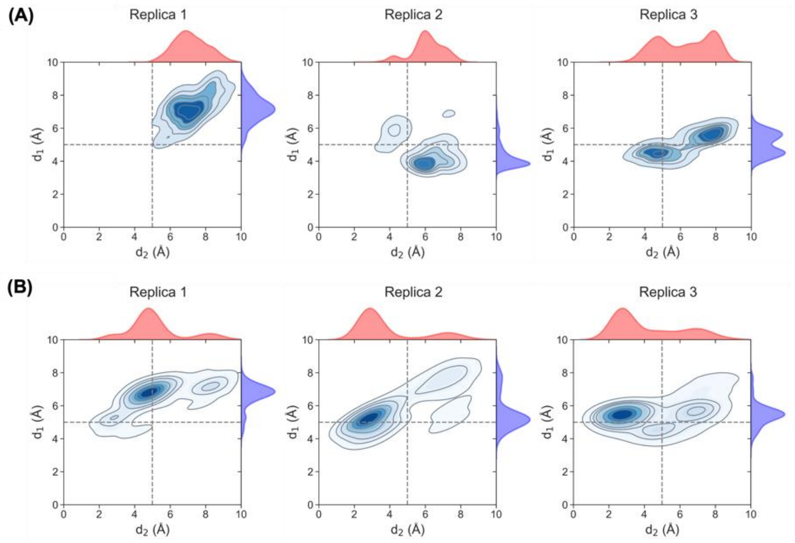


**Supplementary Figure 34**. Kernel density estimation (KDE) of the catalytic distance distributions obtained from 10 ns conventional MD simulations of the *Sr*BDH1 N32_I111L variant in complex with (+)- and (−)-borneol. Distance $d_1$ (blue) corresponds to the hydride-transfer distance, while $d_2$ (red) represents the interaction between the catalytic tyrosine and the substrate. Three independent replicas were used for sampling.

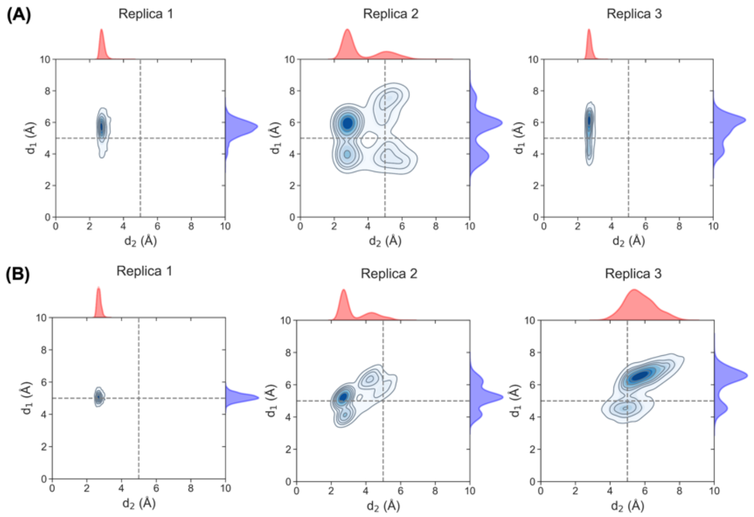


**Supplementary Figure 35**. Kernel density estimation (KDE) of the catalytic distance distributions obtained from 10 ns conventional MD simulations of the *Sr*BDH1 N30_L111I variant in complex with (+)- and (−)-borneol. Distance $d_1$ (blue) corresponds to the hydride-transfer distance, while $d_2$ (red) represents the interaction between the catalytic tyrosine and the substrate. Three independent replicas were used for sampling.

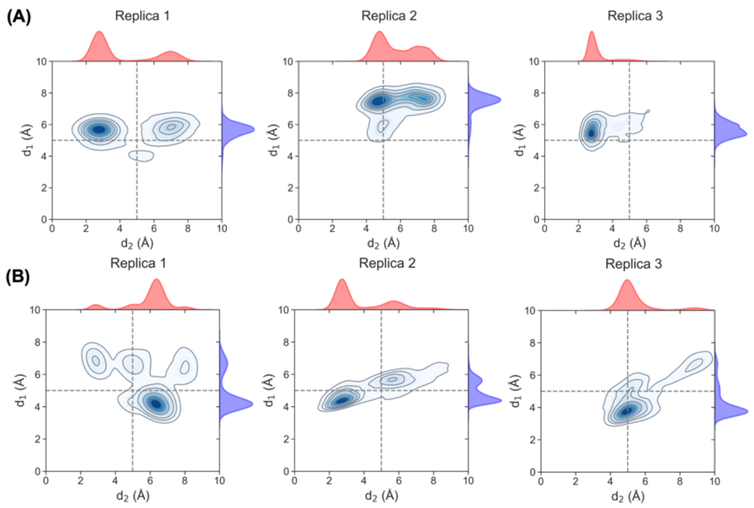


**Supplementary Figure 36**. Kernel density estimation (KDE) of the catalytic distance distributions obtained from 10 ns conventional MD simulations of the *Sr*BDH1 N30_L111I/G169A/V136L/V183I variant in complex with (+)- and (−)-borneol. Distance $d_1$ (blue) corresponds to the hydride-transfer distance, while $d_2$ (red) represents the interaction between the catalytic tyrosine and the substrate. Three independent replicas were used for sampling.

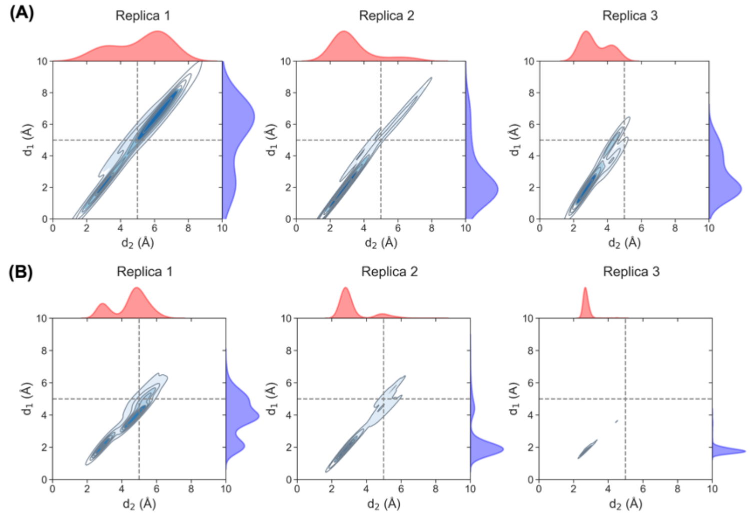


**Supplementary Figure 37**. Kernel density estimation (KDE) of the catalytic distance distributions obtained from 10 ns conventional MD simulations of the *Sr*BDH1 N6 variant in complex with (+)- and (−)-borneol. Distance $d_1$ (blue) corresponds to the hydride-transfer distance, while $d_2$ (red) represents the interaction between the catalytic tyrosine and the substrate. Three independent replicas were used for sampling.

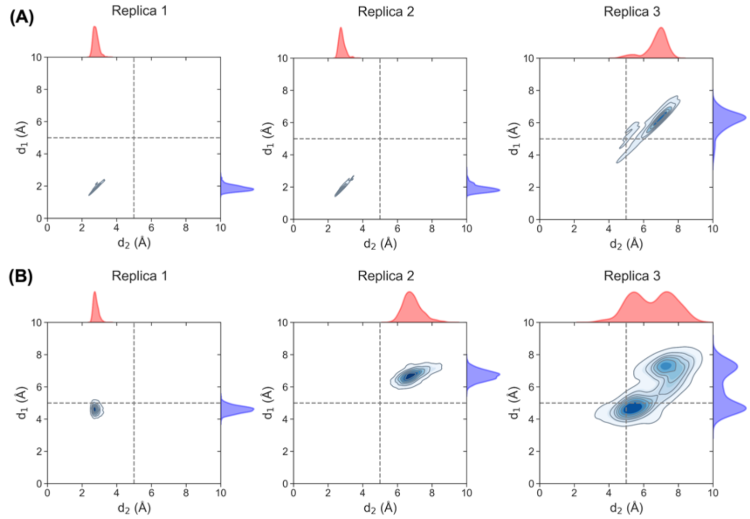


**Supplementary Figure 38**. Kernel density estimation (KDE) of the catalytic distance distributions obtained from 10 ns conventional MD simulations of the *Sr*BDH1 N32 variant in complex with (+)- and (−)-borneol. Distance $d_1$ (blue) corresponds to the hydride-transfer distance, while $d_2$ (red) represents the interaction between the catalytic tyrosine and the substrate. Three independent replicas were used for sampling.

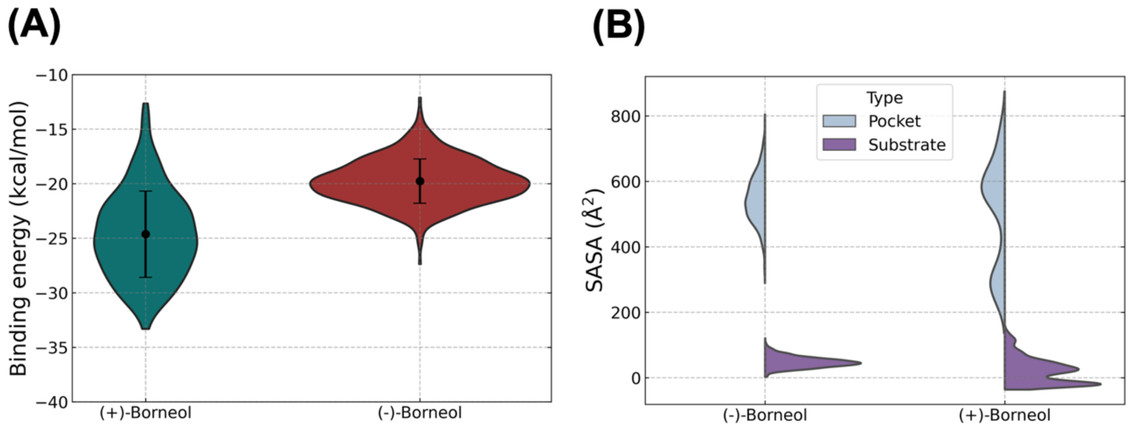


| System | Mean ± SD |
|---|---|
| (+)-Borneol | -24.6 ± 4.0 |
| (-)-Borneol | -19.8 ± 2.0 |

| System | Pocket | Substrate |
|---|---|---|
| (-)-Borneol | 546.5 ± 74.4 | 47.9 ± 16.6 |
| (+)-Borneol | 503.9 ± 166.0 | 21.0 ± 39.4 |

**Supplementary Figure 39**. Violin plots of (A) total binding-energy from MM-ISMSA[11] analysis for (+)-borneol (C) (−)-borneol and (B) of solvent-accessible surface area (SASA, Å²) for the binding pocket (grey blue) and the substrate (purple) obtained from 10 ns ML/MM [2–52–5] simulations within 3 replicas. Mean values and standard deviations are summarized in the accompanying tables.

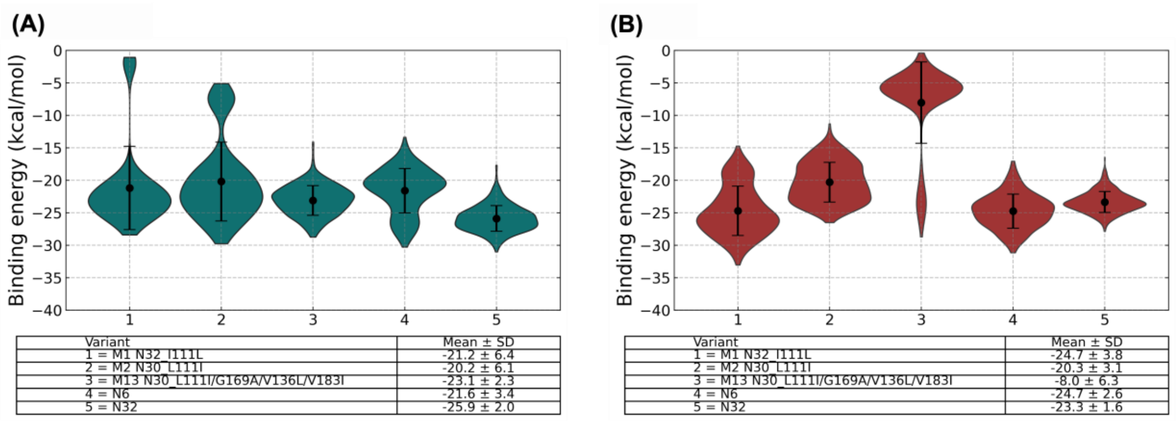


**Supplementary Figure 40**. Mean MM-ISMSA[11] binding energies in *Sr*BDH1 variants (N32_I111L, N30_L111I, N30_L111I/G169A/V136L/V183I, N6, N32) in complex with (A) (+)- and (B) (−)-borneol from 3x10 ns Torch-ANI AMBER ML/MM simulations[2–52–5].

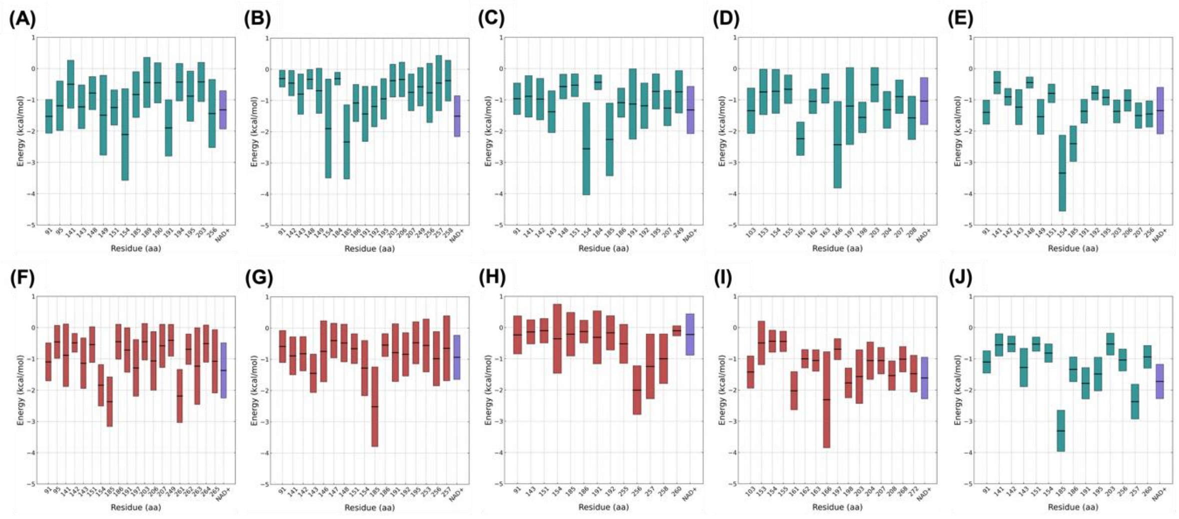


**Supplementary Figure 41**. Per-residue energy decomposition from MM-ISMSA analysis for (+)-borneol (A-E) (−)-borneol (F-J) from 3x10 ns Torch-ANI AMBER ML/MM simulations[2–52–5]. Each plot displays the interaction energies for key residues within (A, F) N32_I111L, (B, G) N30_L111I, (C, H) N30_L111I/G169A/V136L/V183I, (D, I) N6, (E, J) N32 BDH. Shown here are energies for (+)-borneol (teal color) and (−)-borneol (firebrick) and $NAD^+$ (slate blue).

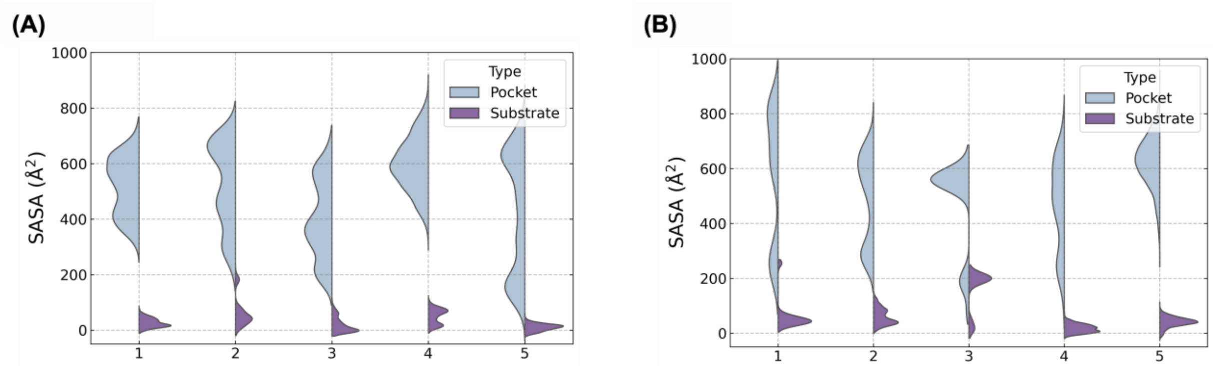


(A)

| Node | Pocket | Substrate |
|---|---|---|
| 1=M1 N32_I111L | 517.6 ± 104.5 | 26.8 ± 16.6 |
| 2=M2 N30_L111I | 520.7 ± 152.1 | 60.6 ± 50.7 |
| 3=M13 N30_L111I/G169A/V136L/V183I | 397.1 ± 140.6 | 19.2 ± 27.2 |
| 4=N6 | 598.1 ± 98.2 | 50.2 ± 27.5 |
| 5=N32 | 424.8 ± 219.5 | 10.6 ± 12.8 |

(B)

| Node | Pocket | Substrate |
|---|---|---|
| 1=M1 N32_I111L | 572.6 ± 241.4 | 62.9 ± 57.9 |
| 2=M2 N30_L111I | 489.8 ± 161.1 | 61.4 ± 30.1 |
| 3=M13 N30_L111I/G169A/V136L/V183I | 472.2 ± 169.4 | 160.4 ± 75.5 |
| 4=N6 | 468.2 ± 166.7 | 18.3 ± 15.5 |
| 5=N32 | 624.9 ± 91.2 | 41.6 ± 18.6 |

**Supplementary Figure 42**. Mean solvent accessible surface area (SASA, $Å^2$) distributions for pocket and substrate in *Sr*BDH1 variants (N32_I111L, N30_L111I, N30_L111I/G169A/V136L/V183I, N6, N32) in complex with (A) (+)- and (B) (−)-borneol from 3x10 ns Torch-ANI AMBER ML/MM simulations2–52–5.

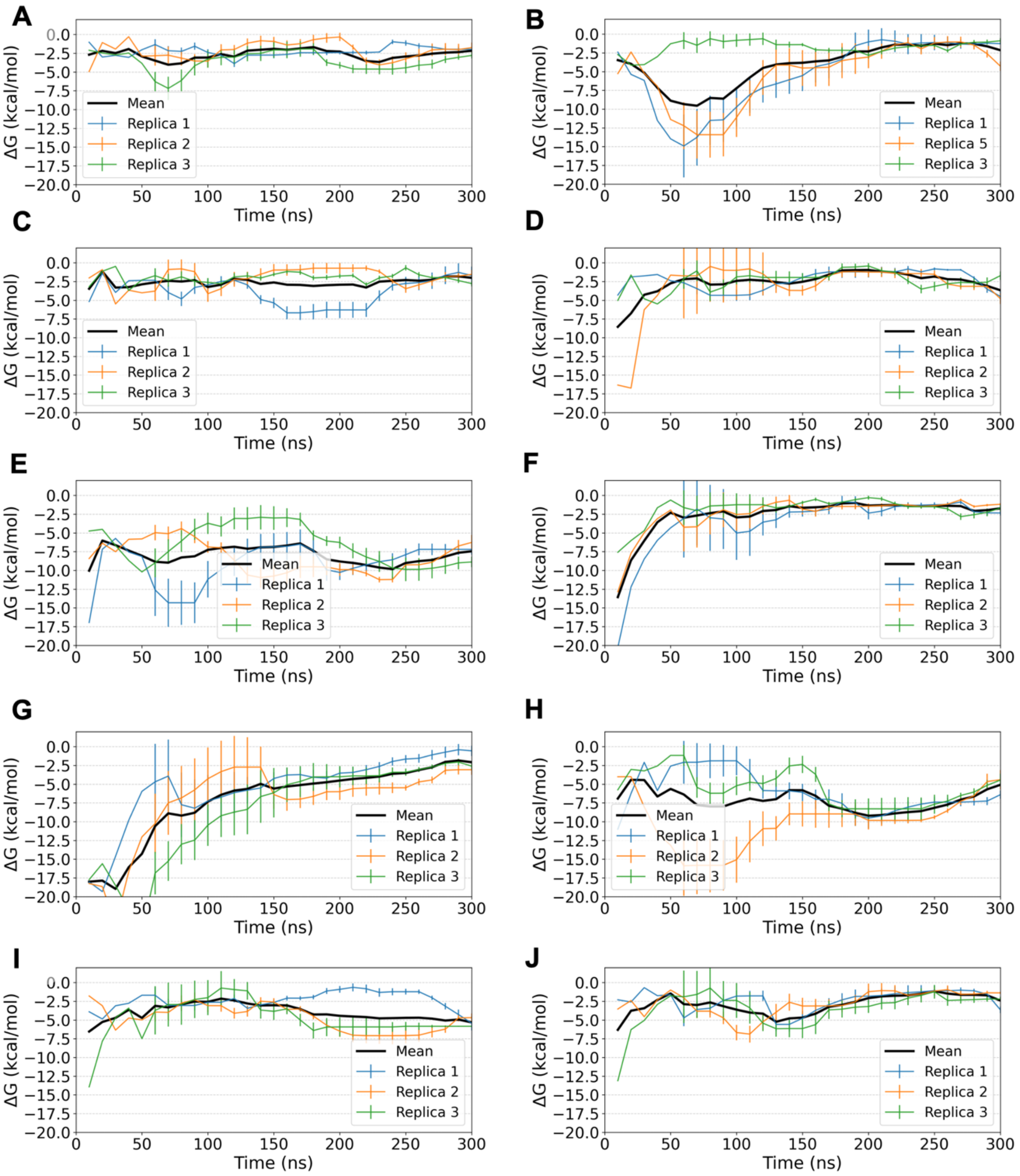


**Supplementary Figure 43**. Binding free-energy profiles (ΔG) as a function of simulation time obtained from funnel metadynamics simulations6–106–10 of *Sr*BDH1 variants in complex with (+) and (−)-borneol. Panels show variants (A, B) N32_I111L, (C, D) N30_L111I, (E, F) N30_L111I/G169A/V136L/V183I, (G, H) N6, and (I, J) N32, with (+)-borneol on the left panels and (−)-borneol in the right panels. Three independent 300 ns replicas are shown as colored lines, while the corresponding mean ΔG profile is shown in black. Error bars indicate the standard deviation across replicas.

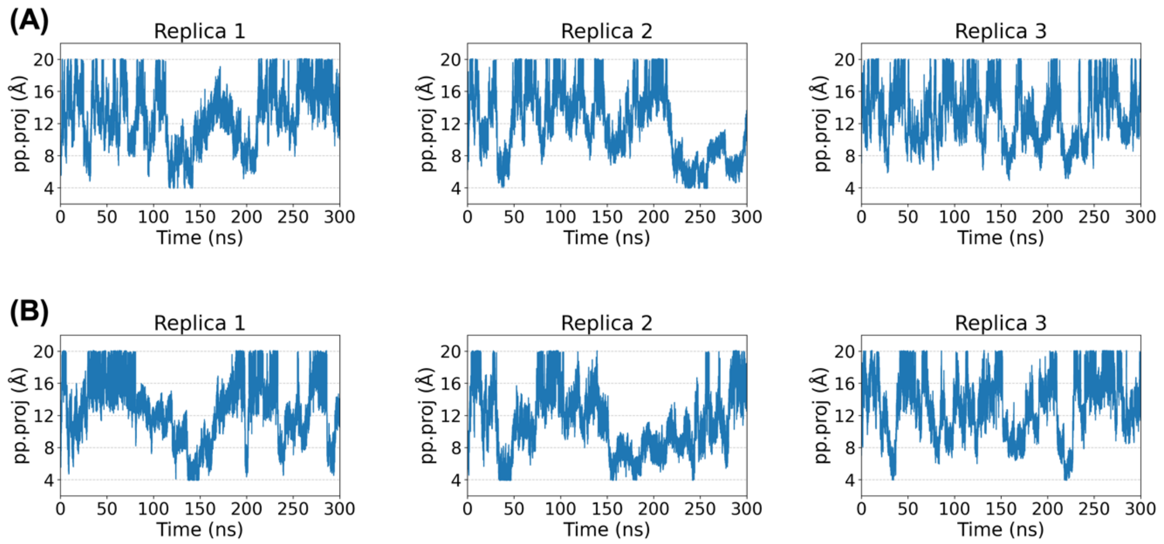


**Supplementary Figure 44**. Time evolution of the funnel coordinate pp.proj along the funnel axis, during 300 ns funnel metadynamics simulations of wild-type *Sr*BDH1. Three independent replicas are shown for (A) (+)-borneol and (B) (−)-borneol, illustrating ligand movement along the binding pathway over time.

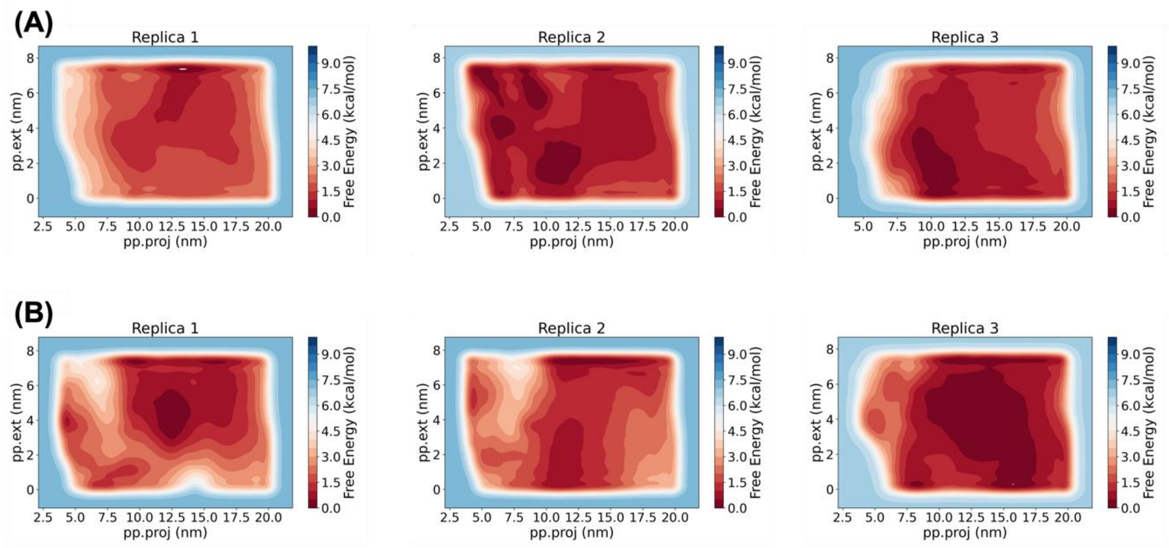


**Supplementary Figure 45**. Two-dimensional free-energy surfaces projected onto the funnel collective variables pp.proj and pp.ext, obtained from funnel metadynamics simulations of wild-type *Sr*BDH1. Three independent replicas are shown for (A) (+)-borneol and (B) (−)-borneol.

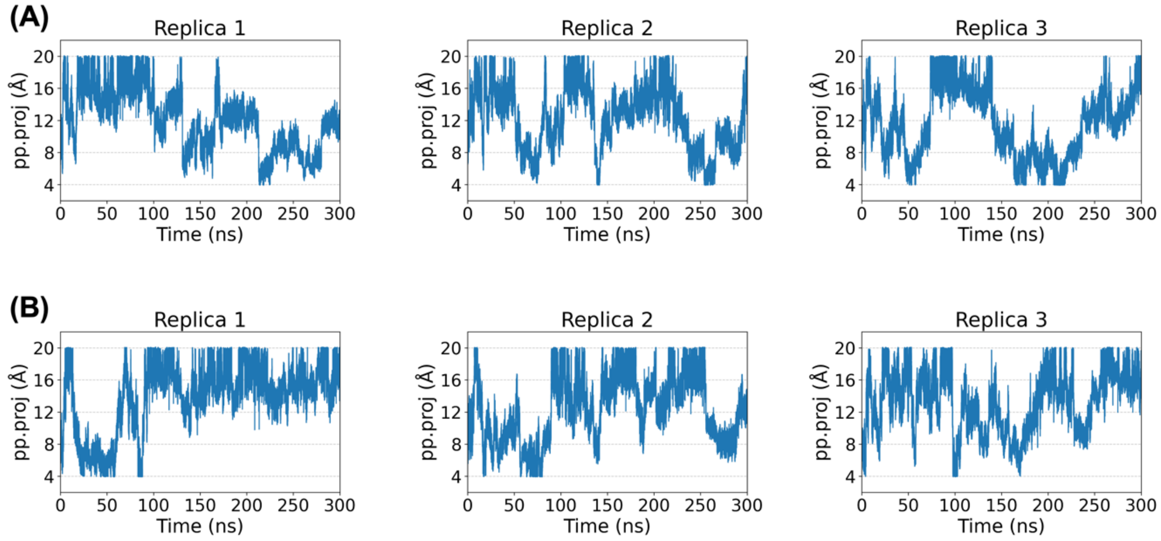


**Supplementary Figure 46**. Time evolution of the funnel coordinate pp.proj along the funnel axis, during 300 ns funnel metadynamics simulations of the N32_I111L BDH variant. Three independent replicas are shown for (A) (+)-borneol and (B) (−)-borneol, illustrating ligand movement along the binding pathway over time.

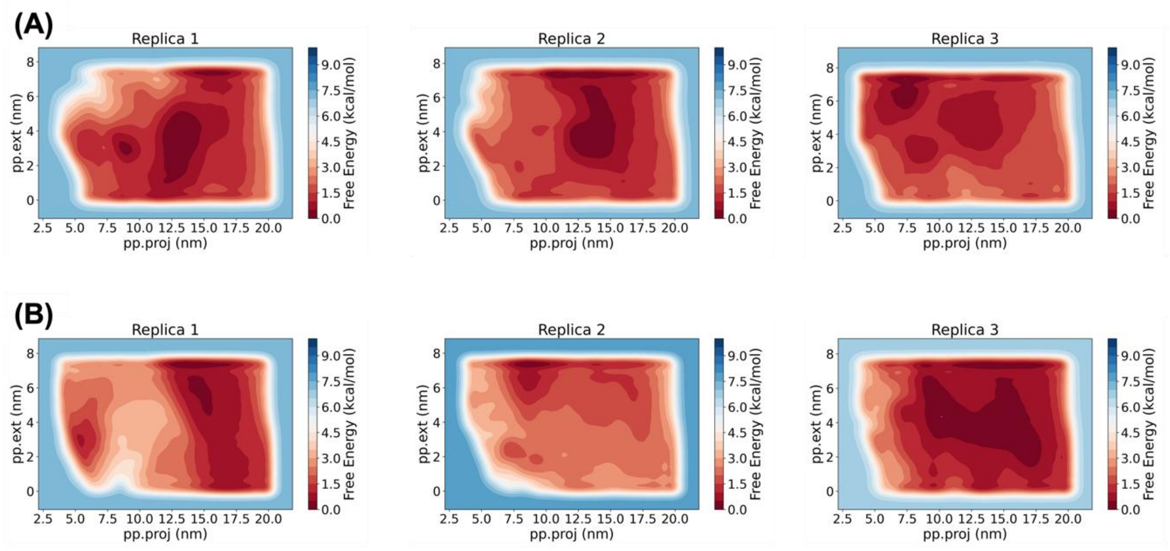


**Supplementary Figure 47**. Two-dimensional free-energy surfaces projected onto the funnel collective variables pp.proj and pp.ext, obtained from funnel metadynamics simulations of the N32_I111L variant. Three independent replicas are shown for (A) (+)-borneol and (B) (−)-borneol.

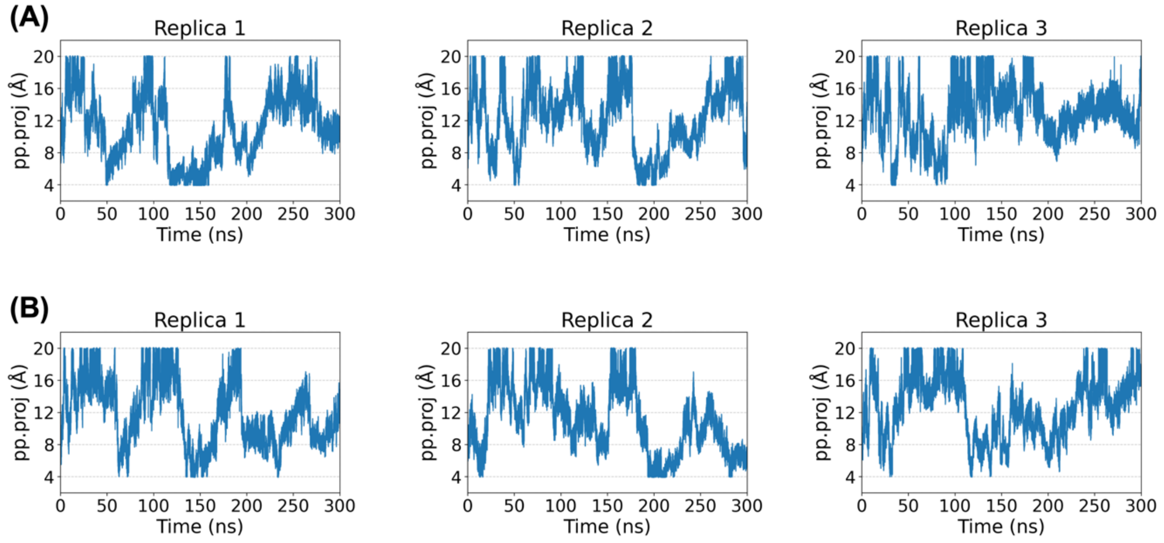


**Supplementary Figure 48.** Time evolution of the funnel coordinate pp.proj along the funnel axis, during 300 ns funnel metadynamics simulations of the N30_L111I variant. Three independent replicas are shown for (A) (+)-borneol and (B) (−)-borneol, illustrating ligand movement along the binding pathway over time.

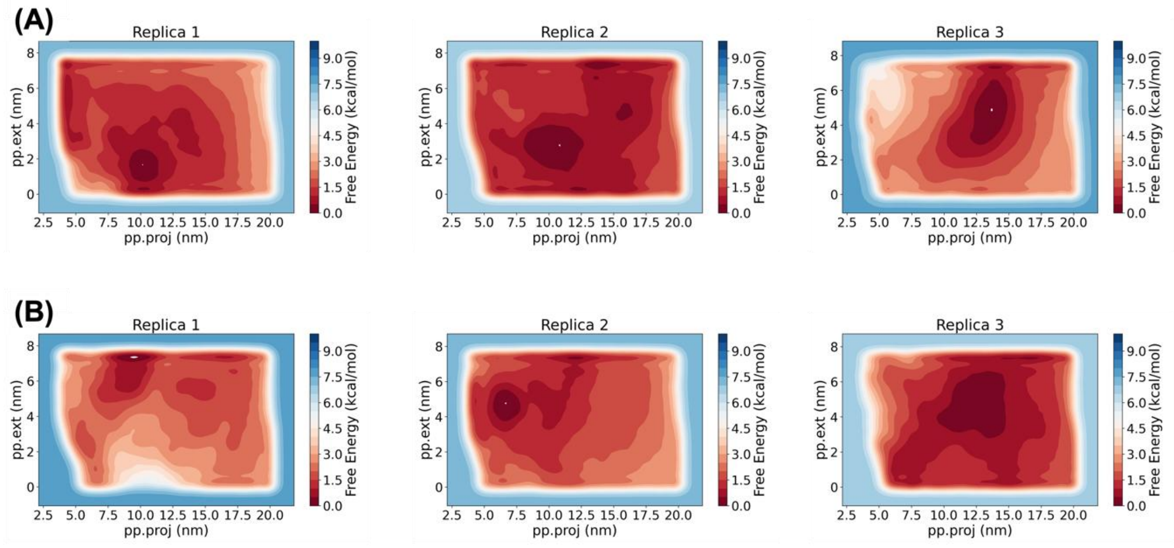


**Supplementary Figure 49**. Two-dimensional free-energy surfaces projected onto the funnel collective variables pp.proj and pp.ext, obtained from funnel metadynamics simulations of the N30_L111I variant. Three independent replicas are shown for (A) (+)-borneol and (B) (−)-borneol.

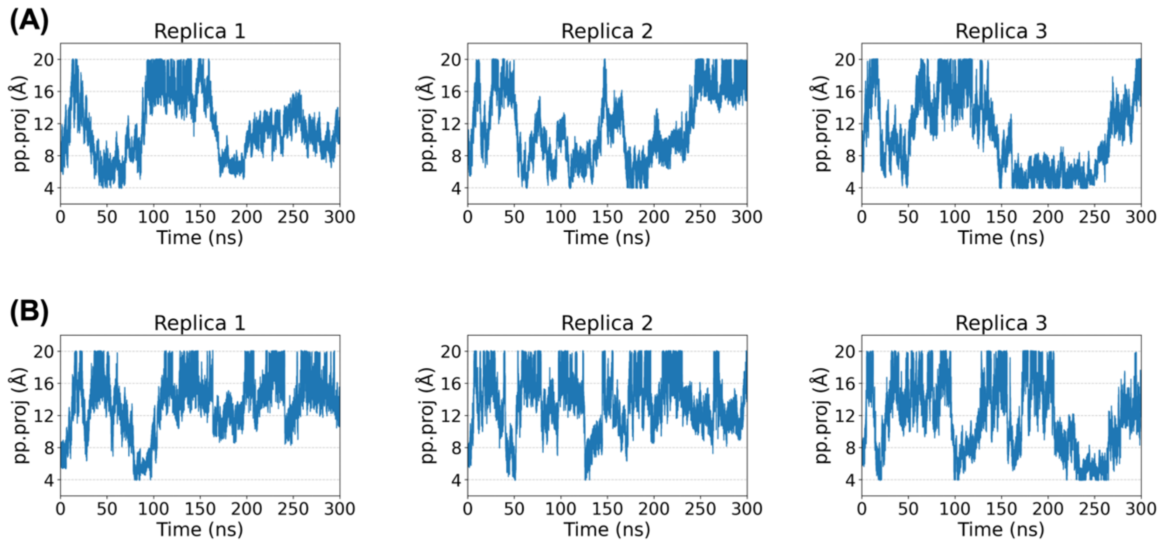


**Supplementary Figure 50**. Time evolution of the funnel coordinate pp.proj along the funnel axis, during 300 ns funnel metadynamics simulations of the N30_L111I/G169A/V136L/V183I variant. Three independent replicas are shown for (A) (+)-borneol and (B) (−)-borneol, illustrating ligand movement along the binding pathway over time.

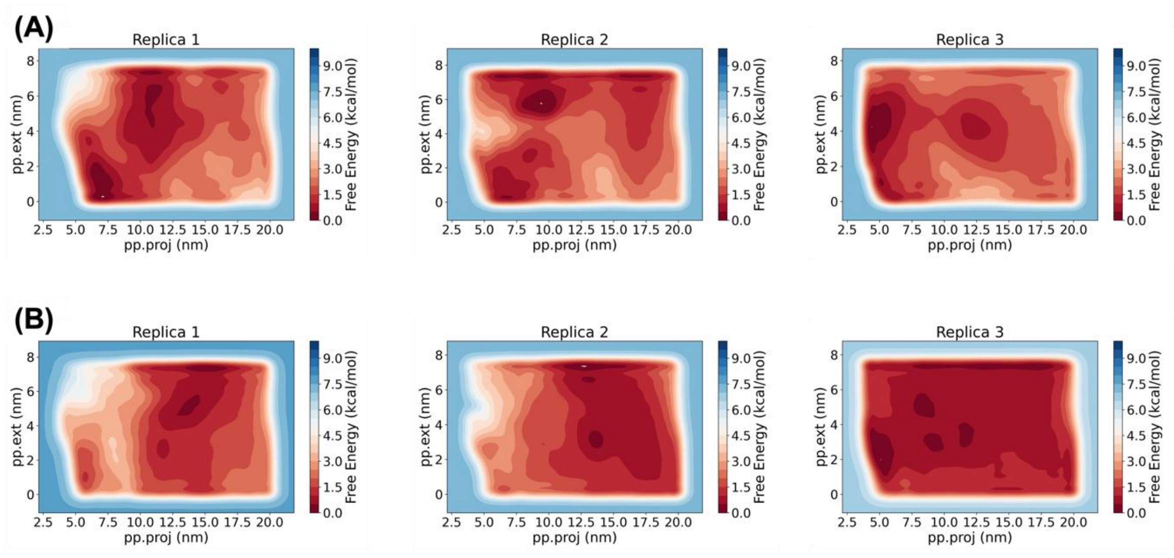


**Supplementary Figure 51**. Two-dimensional free-energy surfaces projected onto the funnel collective variables pp.proj and pp.ext, obtained from funnel metadynamics simulations of the N30_L111I/G169A/V136L/V183I variant. Three independent replicas are shown for (A) (+)-borneol and (B) (−)-borneol.

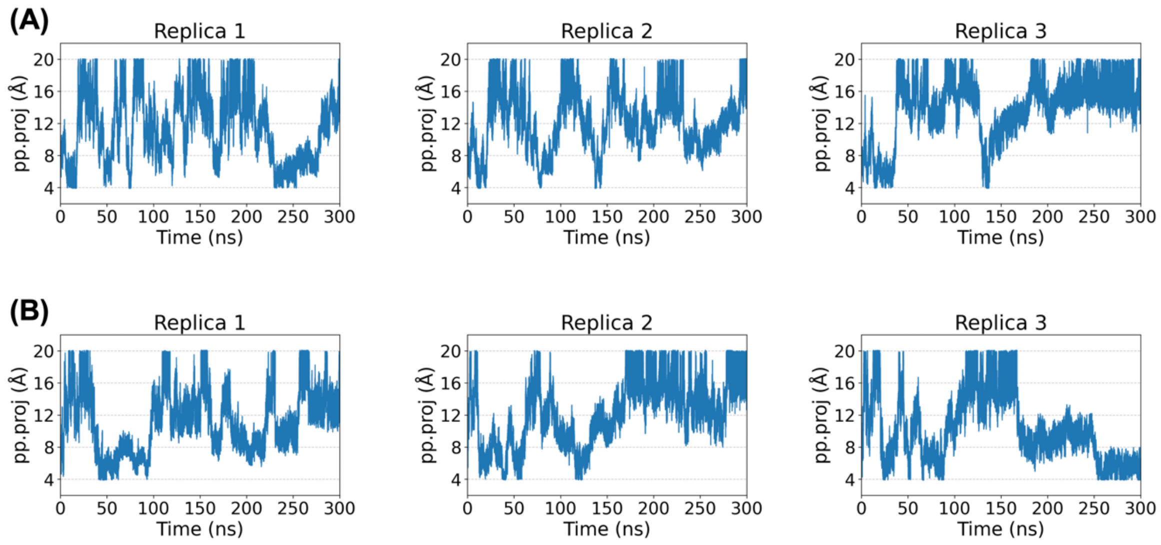


**Supplementary Figure 52**. Time evolution of the funnel coordinate pp.proj along the funnel axis, during 300 ns funnel metadynamics simulations of the N6 variant. Three independent replicas are shown for (A) (+)-borneol and (B) (−)-borneol, illustrating ligand movement along the binding pathway over time.

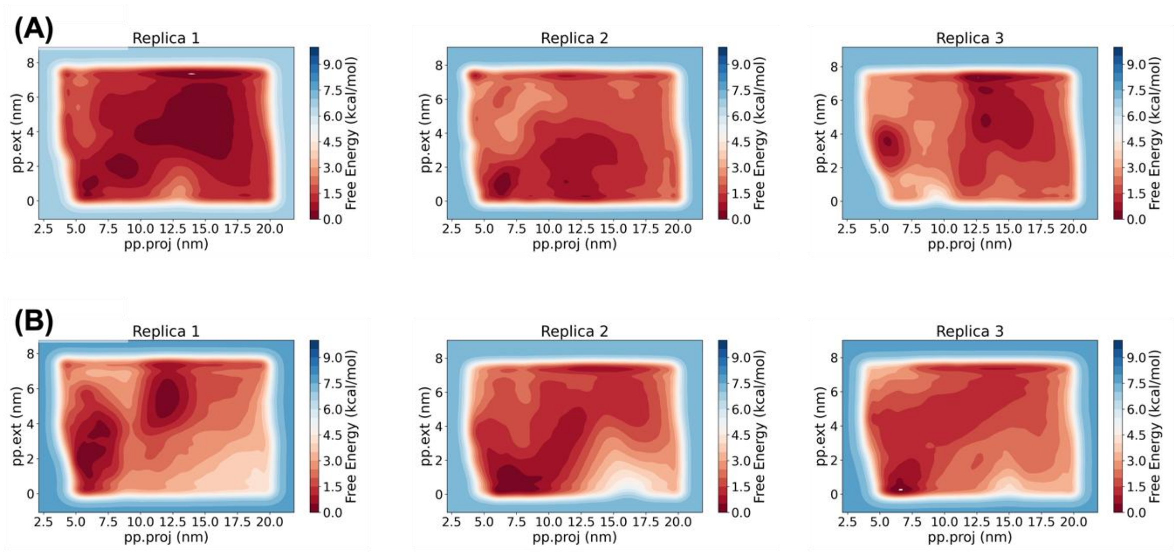


**Supplementary Figure 53**. Two-dimensional free-energy surfaces projected onto the funnel collective variables pp.proj and pp.ext, obtained from funnel metadynamics simulations of the N6 variant. Three independent replicas are shown for (A) (+)-borneol and (B) (−)-borneol.

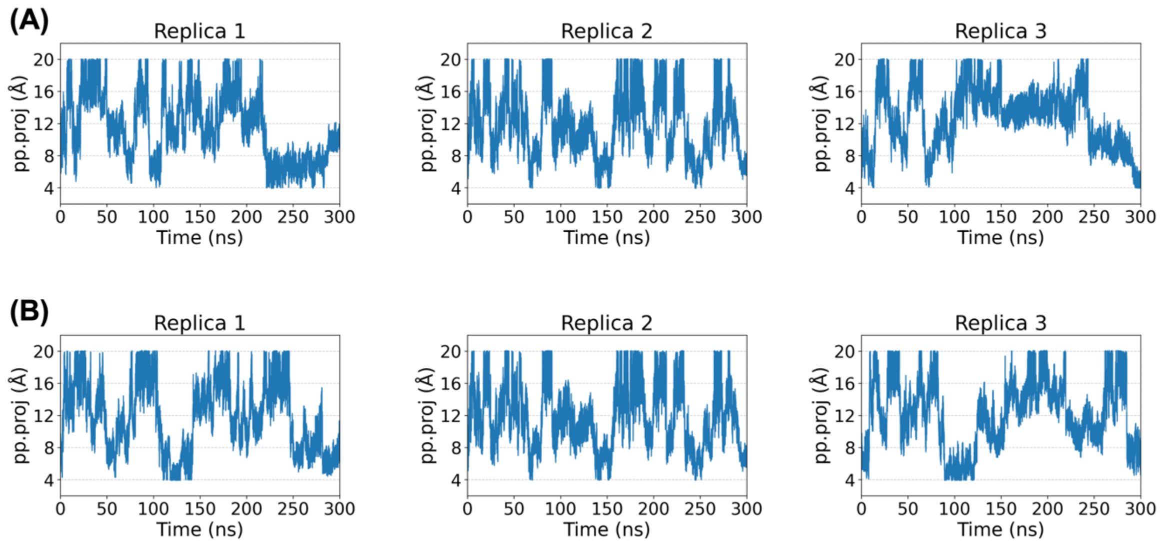


**Supplementary Figure 54**. Time evolution of the funnel coordinate pp.proj along the funnel axis, during 300 ns funnel metadynamics simulations of the N32 variant. Three independent replicas are shown for (A) (+)-borneol and (B) (−)-borneol, illustrating ligand movement along the binding pathway over time.

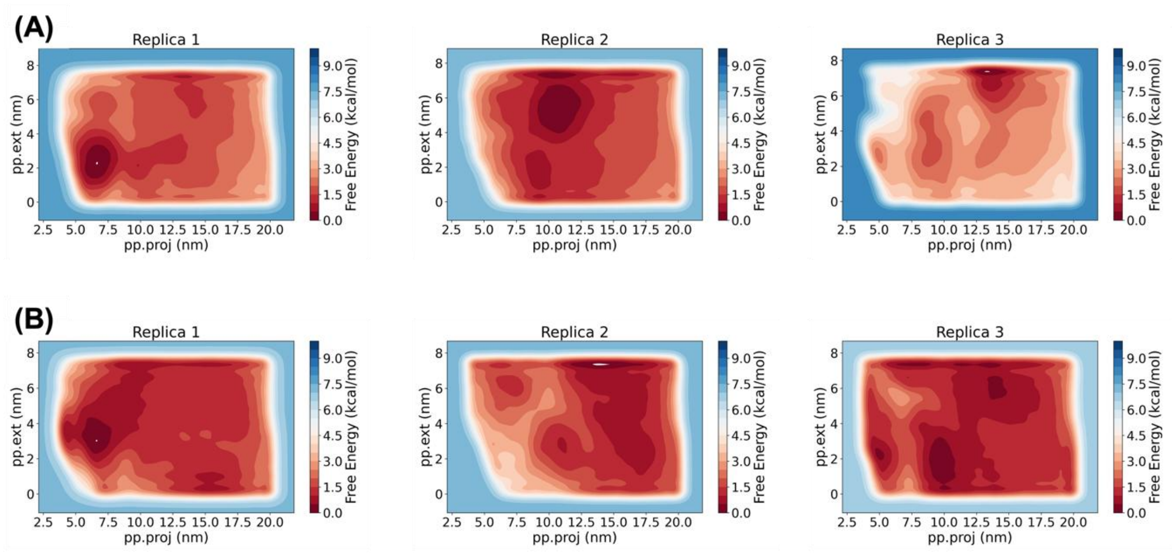


**Supplementary Figure 55**. Two-dimensional free-energy surfaces projected onto the funnel collective variables pp.proj and pp.ext, obtained from funnel metadynamics simulations of the N32 variant. Three independent replicas are shown for (A) (+)-borneol and (B) (−)-borneol.

*SDS-PAGE*

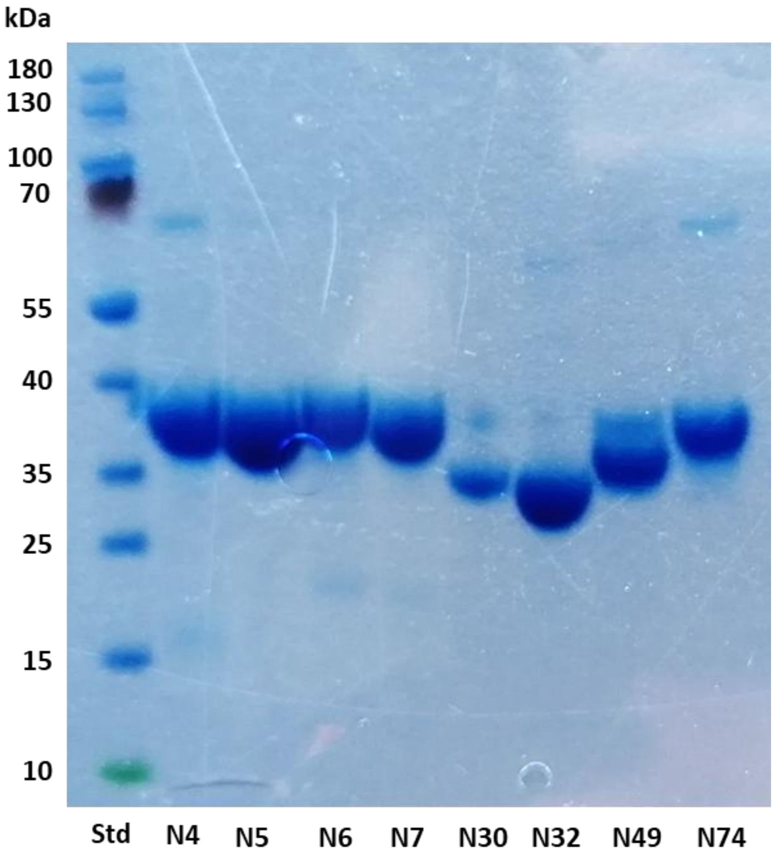


**Supplementary Figure 56.** SDS-PAGE of BDH ancestors. Expression from pET28a(+) in *E. coli* BL21(DE3) at 28°C with 1 mM IPTG, overnight. Purified fractions after His-tag affinity chromatography are shown. 1 mg/mL protein applied to the SDS-PAGE. BDH ancestors are found between 25-40 kDa. PageRuler Prestained (ThermoScientific) was applied as standard (Std).

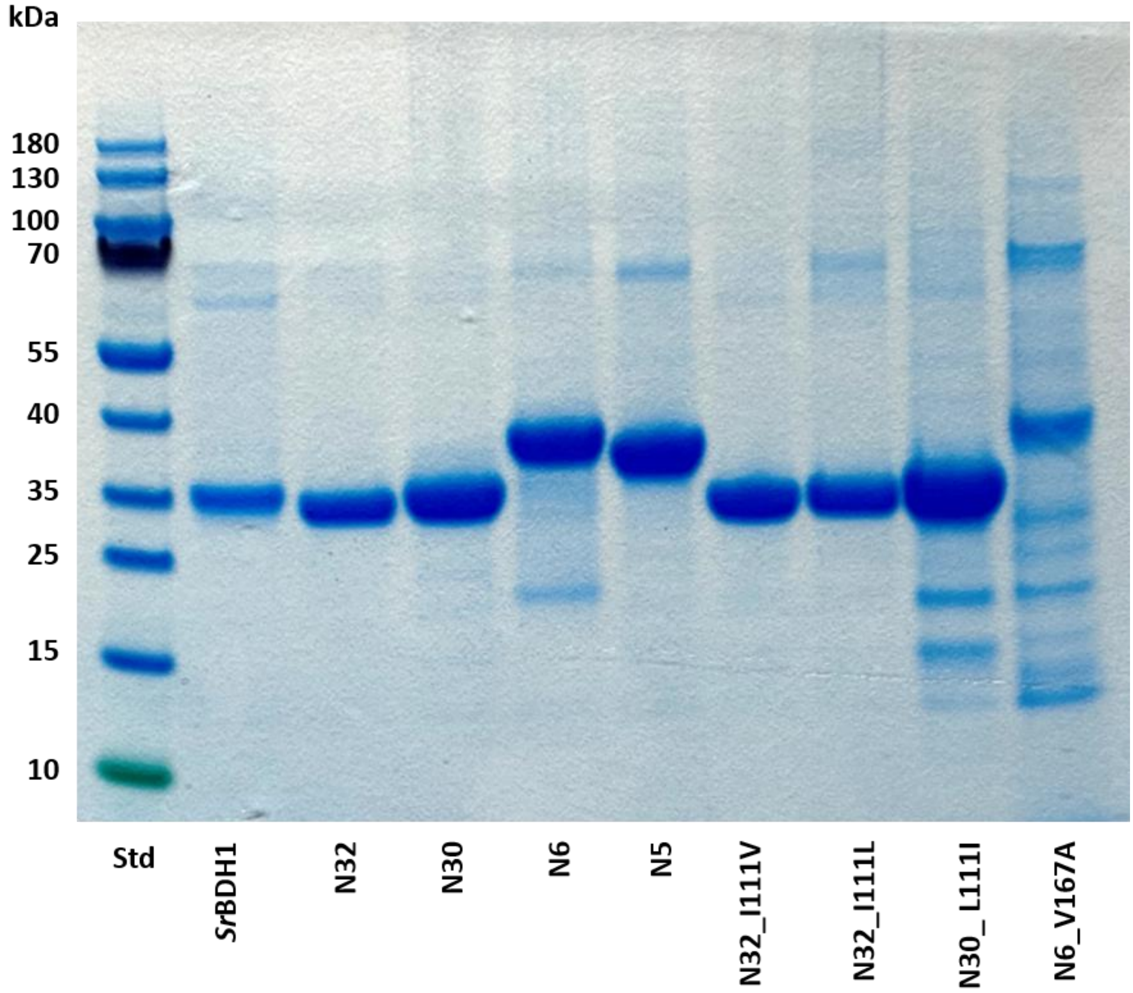


**Supplementary Figure 57.** SDS-PAGE of BDH ancestors. Expression from pET28a(+) in *E. coli* BL21(DE3) at 28°C with 1 mM IPTG, overnight. Purified fractions after His-tag affinity chromatography are shown. 1mg/mL protein applied to the SDS-PAGE. BDH ancestors are found between 25-40 kDa. PageRuler Prestained (ThermoScientific) was applied as standard (Std).

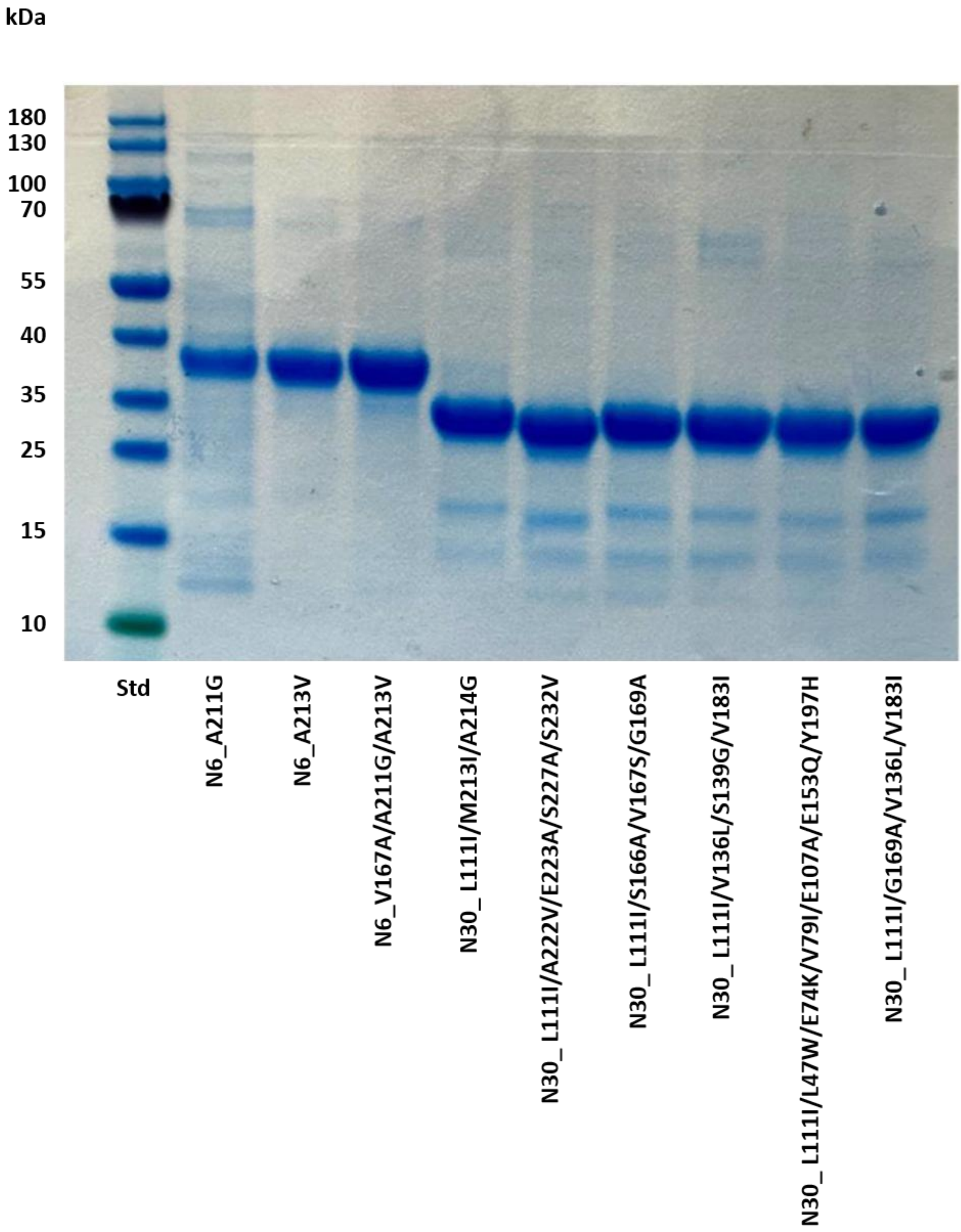


**Supplementary Figure 58.** SDS-PAGE of BDH variants. Expression from pET28a(+) in *E. coli* BL21(DE3) at 28°C with 1 mM IPTG, overnight. Purified fractions after His-tag affinity chromatography are shown. 1mg/mL protein applied to the SDS-PAGE. BDH ancestors are found between 25-40 kDa. PageRuler Prestained (ThermoScientific) was applied as standard (Std).

*Structure determination*

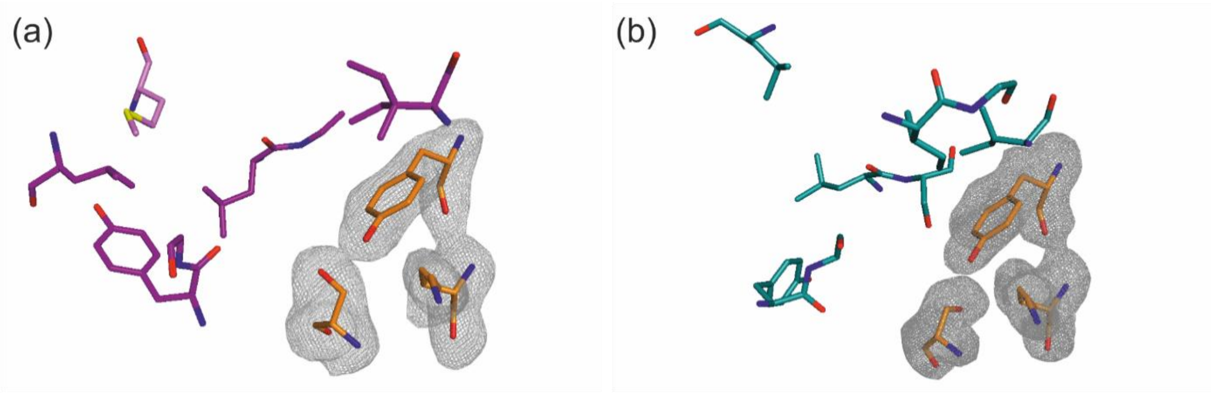


**Supplementary Figure 59.** (A) Polder map contoured at 3σ, showing electric density of the catalytic residues of N32. (B) Polder map contoured at 3σ of the catalytic residues of N39.

**Supplementary Tables**

*Kinetic values of ancestral mutant variants*

**Supplementary Table 1**. Specific activity and $K_{app}$-values determined by gas chromatography for BDH variants by oxidation of (+)-1-borneol enantiomers.

| **Enzyme** | **Specific activity[[mU/mg]** (+)-**1-borneol** | $K_{app}$ **[mM]** |
|---|---|---|
| N32_I111L | 36.6 ± 2.3 | 5.80 ± 2.5 |
| N30_L111I | 33.2 ± 1.4 | 18.11 ± 5.0 |
| N30_L111I/G169A/V136L/V183I | 15.3 ± 1.3 | 11.86 ± 1.4 |

*Protein yield*

**Supplementary Table 2.** Enzyme yield mg Protein per litre cultivation media. Expression from pET28a(+) in *E. coli* BL21(DE3) at 28 °C with 1 mM IPTG, overnight. Purified fractions after His-tag affinity chromatography.

| Enzyme | Yield (mg $L^{-1}$ of culture) |
|---|---|
| N4 | 21.3 |
| N5 | 17.8 |
| N6 | 45.9 |
| N7 | 34.9 |
| N30 | 60.4 |
| N32 | 92.0 |
| N39 | 70.0 |
| N48 | n.d. |
| N49 | 30.2 |

| N74 | 77.8 |
|---|---|

*Apparent $K_M$-values*

**Supplementary Table 3.** Kinetic parameters of the ancestors as determined by applying Michaelis Menten kinetics calculated in Sigma Plot.

| | $K_{app}$ [mM] | $v_{max}$ [mU/mg] | $K_i$ [mM] |
|---|---|---|---|
| **N6** | 0.01±0.01 | 49.81±4.40 | 2.45±0.90 |
| **N30** | 20.27±7.21 | 29.95±8.02 | n.d. |
| **N32** | 17.36±5.40 | 62.61±14.05 | n.d. |

*Crystallographic data*

**Supplementary Table 4.** Crystallographic data collection, refinement, and validation statistics.

| | **N32** | **N39** |
|---|---|---|
| PDB entry | 8R0C | 8R0D |
| **Data collection** | | |
| Wavelength [Å] | 0.9184 | 0.9184 |
| Temperature [K] | 100 | 100 |
| Space group | $P2_1$ | I222 |
| Unit Cell Parameters<br>a, b, c [Å]<br>α, β, γ [°] | <br>71.6 93.5 78.8<br>90.0 105.6 90.0 | <br>62.9 85.5 109.7<br>90.0 90.0 90.0 |
| Resolution [Å][a] | 50.00 - 1.99<br>(2.11 - 1.99) | 50.00 - 1.85<br>(1.96 - 1.85) |

| | | |
|---|---|---|
| Reflections [a] | 470,800 (73,733) | 138,314 (21,199) |
| Unique [a] | 67,766 (10,487) | 25,625 (4,084) |
| Completeness [%][a] | 99.1 (95.3) | 99.7 (99.1) |
| Multiplicity [a] | 6.9 (7.0) | 5.4 (5.2) |
| Data quality | | |
| Intensity [I/σ(I)] [a] | 10.30 (1.00) | 16.8 (1.13) |
| $R_{meas}$ [%][a,b] | 0.150 (1.873) | 0.168 (1.446) |
| $CC_{1/2}$ [a,c] | 99.8 (46.8) | 99.6 (49.7) |
| Wilson B value [$Å^2$] | 40.0 | 29.3 |
| **Refinement** | | |
| Resolution [Å][a] | 50.00 - 1.99 (2.04 - 1.99) | 50.00 - 1.85 (1.92 - 1.85) |
| Reflections [a] | | |
| Number | 67,722 (4,039) | 25.614 (2,788) |
| Test Set [%] | 3.1 | 5.0 |
| $R_{work}$ [%][a] | 0.174 (0.330) | 0.175 (0.329) |
| $R_{free}$ [%][a] | 0.219 (0.375) | 0.210 (0.404) |
| Asymmetric Unit | 4 | 1 |
| Protein residues chain A | 247 | 251 |
| Protein residues chain B | 251 | - |
| Protein residues chain C | 260 | - |
| Protein residues chain D | 253 | - |
| glycerol molecules | 4 | - |
| ethylene glycol molecules | - | 7 |

| | | |
|---|---|---|
| formiat molecules | - | 1 |
| water molecules | 329 | 154 |
| Temperature factors [Å$^2$] | | |
| All Atoms | 43.6 | 32.0 |
| Chain A | 45.9 | 31.2 |
| Chain B | 45.2 | - |
| Chain C | 42.0 | - |
| Chain D | 41.3 | - |
| glycerol | 52.2 | - |
| ethylene glycol | - | 40.9 |
| formiat | - | 32.5 |
| water molecules | 42.6 | 40.6 |
| RMSD from Target Geometry [d] | | |
| Bond Lengths [Å] | 0.007 | 0.010 |
| Bond Angles [°] | 0.888 | 1.015 |
| **Validation Statistics** | | |
| Ramachandran Plot [f] | | |
| Residues in Allowed Regions [%] | 2.6 | 3.2 |
| Residues in Favoured Regions [%] | 97.3 | 96.8 |
| Ramachandran plot Z-score [f] (RMSD) | | |
| whole | -0.90 (0.24) | -0.75 (0.45) |
| helix | -0.38 (0.20) | 0.35 (0.39) |
| sheet | -0.74 (0.38) | -1.54 (0.68) |
| loop | -0.54 (0.31) | -1.11 (0.59) |

| | | |
|---|---|---|
| MOLPROBITY Clashscore [f] | 4.11 | 1.99 |
| MOLPROBITY score [g] | 1.50 | 1.29 |
| Poor rotamers [%] | 1.47 | 1.46 |
| C-β deviations [%] | 0 | 0 |

[a] data for the highest resolution shell in parenthesis

[b] $R_{meas}(I) = \sum_h [N/(N-1)]^{1/2} \sum_i | I_{ih} - \langle I_h \rangle | / \sum_h \sum_i I_{ih}$, in which $\langle I_h \rangle$ is the mean intensity of symmetry-equivalent reflections h, $I_{ih}$ is the intensity of a particular observation of h and N is the number of redundant observations of reflection h .[11]

[c] $CC_{1/2} = (\langle I^2 \rangle - \langle I \rangle^2) / (\langle I^2 \rangle - \langle I \rangle^2) + \sigma^2_{\varepsilon}$, in which $\sigma^2_{\varepsilon}$ is the mean error within a half-dataset .[12]

[d] RMSD – root mean square deviation.

[e] calculated with PHENIX.[13]

[f] Clashscore is the number of serious steric overlaps (>0.4 Å) per 1,000 atoms

[g] calculated with MOLPROBITY .[14]

**Supplementary Table 5.** Structural comparison of the crystal structures of N32, N39, *So*BDH1, *Sr*BDH2. R.m.s.d. (Å) calculated with SSM[15]as implemented in COOT.

| R.m.s.d. (Å) | N32 | N39 | *So*BDH1 | *Sr*BDH2 |
|---|---|---|---|---|
| N32 | | | | |
| N39 | 1.19 | | | |
| *So*BDH1 | 1.35 | 1.22 | | |
| *Sr*BDH2 | 0.97 | 0.82 | 1.34 | |

**Supplementary Table 6.** RMSDs for AlphaFold3[16] structures.

| RMSD | N6 | N30 | N32 | *Sr*BDH1 |
|---|---|---|---|---|
| N6 | x | 0.246 | 0.249 | 0.308 |
| N30 | | x | 0.15 | 0.322 |
| N32 | | | x | 0.284 |
| *Sr*BDH1 | | | | x |

*Docking and ML/MM data*

**Supplementary Table 7**. Relevant catalytic distances $d_1$ and $d_2$ (Å), obtained from Chai-11211 generated models of different *Sr*BDH1 variants in complex with (+)- and (−)-borneol.

| System | (+)-borneol | | (−)-borneol | |
|---|---|---|---|---|
| | $d_1$ | $d_2$ | $d_1$ | $d_2$ |
| Wildtype | 2.1 | 3.3 | 2.8 | 3.2 |
| N32_I111L | 2.8 | 4.3 | 3.0 | 5.3 |
| N30_L111I | 5.4 | 4.1 | 2.6 | 4.3 |
| N30_L111I/G169A/V136L/V183I | 4.5 | 4.2 | 3.1 | 4.5 |
| N6 | 2.8 | 3.4 | 2.7 | 3.2 |
| N32 | 2.6 | 4.5 | 3.5 | 2.7 |

**Supplementary Table 8.** Summary of funnel metadynamics binding free energies for wild-type and engineered *Sr*BDH1 variants in complex with (+) and (−)-borneol.

| System | ΔG (+) borneol | ΔG (−) borneol | ΔΔG | ΔΔG / $\sigma_{\Delta\Delta G}$ | Assessment[a] |
|---|---|---|---|---|---|
| Wildtype | -5.1 ± 0.5 | -3.6 ± 1.0 | -1.5 ± 1.1 | 1.4 | Not significant |
| N32_I111L | -2.2 ± 0.6 | -2.1 ± 1.8 | -0.1 ± 1.9 | 0.1 | Not significant |
| N30_L111I | -2.0 ± 0.7 | -3.7 ±1.7 | +1.7 ± 1.8 | 0.9 | Weak trend |
| N30_L111I/G169A/ V136L/V183I | -4.9 ± 1.3 | -1.7 ± 0.6 | -3.2 ± 1.4 | 2.3 | Meaningful |
| N6 | -2.1 ± 1.3 | -5.1 ± 1.1 | +3.0 ± 1.7 | 1.8 | Meaningful |
| N32 | -5.3 ± 0.6 | -2.4 ± 1.1 | -2.9 ± 1.3 | 2.2 | Meaningful |

[a] Reported values correspond to the calculated standard binding free energies ($\Delta G_{calc}$), obtained by averaging the final 10 ns of the reconstructed free-energy profiles from each simulation. Values are reported as mean ± standard deviation across three independent 300 ns replicas, with uncertainties estimated using block analysis to account for time correlation in the metadynamics trajectories. Enantioselectivity was quantified as the free-energy difference between enantiomers (ΔΔG = ΔG(+)-borneol - ΔG(−)-borneol), and the associated uncertainty was obtained by propagation of errors assuming independent uncertainties for each enantiomer. The ratio $\Delta\Delta G/\sigma_{\Delta\Delta G}$ is reported as a dimensionless signal-to-noise metric to assess whether the observed enantioselective differences exceed statistical uncertainty. Systems with $\Delta\Delta G/\sigma_{\Delta\Delta G}$ values approaching or exceeding 2 were classified as statistically meaningful, while lower values indicate weak or statistically unresolved trends.

*Structural comparisons*

**Supplementary Table 9**. Structural comparison of the crystal structures of N32, N39, *So*BDH1, *Sr*BDH2. Sequence identity (%) calculated with SSM as implemented in COOT.

| **Sequence identity (%)** | **N32** | **N39** | ***So*BDH1** | ***Sr*BDH2** |
|---|---|---|---|---|
| **N32** | | | | |
| **N39** | 86.87 | | | |
| ***So*BDH1** | 54.10 | 53.73 | | |
| ***Sr*BDH2** | 67.67 | 68.37 | 50.94 | |

*Isoborneol converison*

**Supplementary Table 10**. Overview of biochemical data on 1-isoborneol oxidation of the second mutant variant set.[a]

| | ***r*-1-isobor conv. [mU/mg]** |
|---|---|
| **N30_L111I/N30_L111I13I/A214G** | 40.57 ± 22.60 |
| **N30_L111I/A222V/E223A/S227A/S232V** | 13.20 ± 0.68 |
| **N30_L111I/S166A/V167S/G169A** | 9.56 ± 2.99 |
| **N30_L111I/V136L/S139G/V183I** | 8.95 ± 1.63 |
| **N30_L111I/L47W/E74K/V79I/E107A/E153Q/Y197H** | 18.48 ± 4.03 |
| **N30_L111I/V136L/G169A/V183I** | 43.74 ± 2.80 |
| ***SrBDH1* V111L** | 2.73 ± 0.34 |
| ***Sr*BDH1** | 68.29 ± 16.40 |
| **N32 L111V** | 40.38 ± 12.65 |
| **N30** | n.d. |
| **N32** | n.d. |
| **N32_I111L** | 109.52 ± 38.5 |
| **N30_L111I** | 23.65 ± 8.58 |

[a] Since the mutants displayed a higher selectivity toward the 1-borneol enantiomers, selectivity toward the racemic diastereomer 1-isoborneol was investigated. All variants just as the wildtype *Sr*BDH1 only convert (+)-1-isoborneol and do not accept the enantiomer (−)-1-isoborneol as substrate. Conversion rates toward the substrate, however, vary between the mutants. N32_I111L shows 1.6-fold increased activity, while other variants show up to 25-fold decreased activity compared to *Sr*BDH1. Here *SrBDH1* V97L nearly loses the capacity to convert the diastereomer, thus establishing it as the most stereospecific variant found in this study.

### Supplementary Data

#### *Sequences of ancestral proteins*

>N4_0.25

MASSSSLSSSAKRLEGKVALITGGASGIGECTARLFAKHGAKVVIADIQDDLGQAVCEDLGSESASYVHCDVTIESDVENAVDFA
VSKYGKLDIMFNNAGILDPPKPSILDNEKSDFERVLSVNVTGVFLGMKHAARVMIPARSGSIISTASVASVIGGVASHAYTCSKHA
VVGLTKNVAVELGQYGIRVNCVSPYAVATPMARNFLKLDEEAVENMVSVYANLKGVVLKAEDVAEAALYLASDEAKYVSGHNLV
VDGGFTIVNPSFGMFKQPPNS

>N5_0.85

RLEGKVALITGGASGIGESTVRLFVKHGAKVVIADIQDDLGQAVCEDLGSTESVSYVHCDVTIESDVQNAVDFTVSKYGKLDIMFN
NAGILGPPNPSILDNDKSDFERVLSVNVTGVFLGMKHAARVMIPAKKGSIISTASVASVIGGLGPHAYTASKHAVVGLTKNVAVEL
GQYGIRVNCVSPYAVATPMARNALKVDEEAVENMVSASANLKGVVLKAEDVAEAALYLASDEAKYVSGHNLVVDGGFTSVNPS
LGMFR

>N6_0.95

MSSSSSSSPPAKRLEGKVAIITGGASGIGESTVRLFVQHGAKVVIADIQDDLGQAICEDLGSTENVSYVHCDVTNESDVQNLVDT
TVSKYGKLDIMFNNAGILGRPNSSILDTDKSDFERVLGVNVTGAFLGAKHAARVMIPAKKGCILFTASVASVIGGLGPHAYTASKH
AVVGLTKNLAVELGQYGIRVNCVSPYAVATPMARNALKVDEEAVENMISESANLKGVVLKAEDVAEAALYLASDEAKYVSGLNLV
VDGGFSTVNPSLTMAMKPPNS

>N7_0.84

MSGSSSRAPIAKRLEGKVAIITGGASGIGESTVRLFVQHGAKVVIADIQDDLGQSICKDLGSDENVSYVHCDVTNDSDVQNLVDTT
VSKYGKLDIMFNNAGISGNLNSSILDTDNEDFKRVFDVNVYGAFLGAKHAARVMIPAKKGCILFTSSVASVISGLGPHAYTASKHA
VVGLTKNLCVELGQYGIRVNCISPYAVATPMLRNAMKVDESAVENMISESANLKGVVLKAEDVAEAALYLASDESKYVSGLNLVV
DGGYSTINQSLTMAMKSLSS

>N30_0.94

KRLEGKVAIITGGASGIGASAVRLFLENGAKVVIADIQDDLGQAICDKLGENVSYVHCDVSNEDDIRNLVDTTVAKYGKLDIMFNNA
GILDRPYGSILDTEKSDLERVLGVNVVGSFLGAKHAARVMVPERKGCILFTASACSVIGGLGTHAYTASKHAVVGLMKNLAAELG
QYGIRVNCVSPYGVVTGMARGVSEVDAEQVESMLSESGNLKGAVLKVEDVAQAALYLASDEANYVSGLNLVVDGGFSVVNPSL
MMAL

>N32_0.88

KRLEGKVAIITGGASGIGASAVRLFWENGAKVVIADIQDDLGQAICDKLGKNVSYIHCDVSNEDDIRNLVDTTVAKYGKLDIMFNNA
GIIDRPYGSILDTEKSDLERVLGVNLVGGFLGAKHAARVMVPQRKGCILFTASACASIAGLGTHAYTASKHAIVGLMKNLAAELGQ
HGIRVNCVSPYGVVTGIGRGVSEVDVAQVEAMLSEVGNLKGAVLKVEDVAQAALYLASDEANYVSGLNLVVDGGFSVVNPSMM
MAL

>N39_0.75

SKRLEGKVAIITGGASGIGASTVQLFHENGAKVVIADIQDDLGQAIANKLGKNVCYIHCDVSNEDDIINLVDTTVAKYGKLDIMYNN
AGIIDRPFGSILDTTKSDLERVLGVNLVGAFLGAKHAARVMVPQKKGCILFTASACTAIAGLSTHAYAVSKYGIVGLAKNLAAELGQ
HGIRVNCVSPYGVVTGIGGVSEVDAVVEAMLSEVGNLKGQILKAEGVAKAALYLASDEANYVSGLNLVVDGGFSVVNPTMMKA
LNPPES

>N48_0.57RLEGKVALITGGASGIGESTVRLFVKHGAKVVIADIQDDLGQALCETLGSTSSVSYVHCDVTIESDVQNAVDFTVSKY
GKLDIMFNNAGILGPPNPSILDFDLSDFERVLSVNVKGVFLGMKHAARVMIPAKKGSIISTASVASVIGGLGPHAYTASKHAVVGLT
KNVAVELGQYGIRVNCVSPYAVATPMALNALKVDEEAVENFVGASANLKGVVLKAEDVAEAALYLASDEAKYVSGHNLVVDGGF
TSSNHSLGMFR

>N49_0.88

MAATSSSRSSLPAQRLLGKVALVTGGASGIGESIVRLFYKHGAKVCIADIQDDLGQRLCETLGGNSSVSFCHCDVTIEADVQRAV
DFTVDKFGSLDIMVNNAGISGPPCPDIRDFDLSVFERVFDVNVKGVFLGMKHAARVMIPAKKGSIISTCSVASVIGGIGPHAYTGS
KHAVLGLTKNVAAELGKHGIRVNCVSPYAVATSLALAHLPEDERTEDALVGFRNFVGANANLQGVELTAEDVANAVVFLASDEA
RYVSGANLMVDGGFTSSNHSLRVFR

>N74_0.93

MASSSLLSAVAKRLEGKVALITGGASGIGECTARLFSKHGAKVVIADIQDDLGQSVCEDLGSESASFVHCDVTKESDVENAVDFA
VSKYGKLDIMFNNAGIVDEPKPSILDNEKSDFERVLSVNLTGVFLGTKHAARVMIPARSGSIISTASIASVIGGVASHAYTCSKHGV
VGLTKNAAVELGQYGIRVNCVSPYAVATPMARNFLKLDEEAVEGVVSVYANLKGVVLKAEDVAEAALYLASDEAKYVSGHNLVV
DGGFTIVNPSFGMF

*Equations*

**Supplementary Equation S1.** The ratio of the activity ratios of the singular enantiomers over the competitive substrates in relation to each other, mirror the selectivity of the enantiomers over each other. E = enantiomeric ratio. (S) or (R) = activity of enantiomers. reference = activity of competitive substrate.

$$E = \frac{(S)}{(R)} selectivity = \frac{\frac{(S)}{reference} selectivity}{\frac{(R)}{reference} selectivity}$$

**Supplementary Equation S2** Michaelis Menten equation

$$v = \frac{v_{max} * [S]}{K_m + [S]}$$

**Supplementary Equation S3** Michalis Menten Inhibition Model competitive inhibition

$$v = \frac{v_{max} * [S]}{K_m + [S] * (1 + \frac{[S]}{K_i})}$$

**Supplementary Equation S4.** Enantiomeric excess (ee) of most abundant substrate (S).

$$ee_S = \left|\frac{[S^R] - [S^S]}{[S^R] + [S^S]}\right|$$

**Supplementary Equation S5.** Enantiomeric excess (ee) of most abundant product (P).

$$ee_P = \left|\frac{[P^R] - [P^S]}{[P^R] + [P^S]}\right|$$

**Supplementary Equation S6.** E-value formular for irreversible reactions.

$$E = \frac{\ln\left[\frac{1 - ee_S}{1 + \frac{ee_S}{ee_P}}\right]}{\ln\left[\frac{1 + ee_S}{1 + \frac{ee_S}{ee_P}}\right]}$$

**Supplementary Equation S7.** Apparent E-value $E_{app}$ given as the ratio of specific activities of the preferred enantiomer over the other.

$$E_{app} = \frac{v_R}{v_S}$$

### Supplementary References


1. Klett, J. *et al.* MM-ISMSA: An Ultrafast and Accurate Scoring Function for Protein–Protein Docking. *J. Chem. Theory Comput.* **8**, 3395–3408 (2012).

2. Semelak, J. A. *et al.* Advancing Multiscale Molecular Modeling with Machine Learning-Derived Electrostatics. *J. Chem. Theory Comput.* **21**, 5194–5207 (2025).

3. Gao, X., Ramezanghorbani, F., Isayev, O., Smith, J. S. & Roitberg, A. E. TorchANI: A Free and Open Source PyTorch-Based Deep Learning Implementation of the ANI Neural Network Potentials. *J. Chem. Inf. Model.* **60**, 3408–3415 (2020).

4. Pickering, I., Xue, J., Huddleston, K., Terrel, N. & Roitberg, A. E. TorchANI 2.0: An Extensible, High-Performance Library for the Design, Training, and Use of NN-IPs. *J. Chem. Inf. Model.* **65**, 11656–11671 (2025).

5. Pickering, I., Semelak, J. A., Xue, J. & Roitberg, A. E. TorchANI-Amber: Bridging Neural Network Potentials and Classical Biomolecular Simulations. *J. Phys. Chem. B* **129**, 11927–11938 (2025).

6. Karrenbrock, M. *et al.* Absolute Binding Free Energies with OneOPES. *J. Phys. Chem. Lett.* **15**, 9871–9880 (2024).

7. Barducci, A., Bonomi, M. & Parrinello, M. Metadynamics. *WIREs Computational Molecular Science* **1**, 826–843 (2011).

8. Barducci, A., Bussi, G. & Parrinello, M. Well-Tempered Metadynamics: A Smoothly Converging and Tunable Free-Energy Method. *Phys. Rev. Lett.* **100**, 20603 (2008).

9. Limongelli, V., Bonomi, M. & Parrinello, M. Funnel metadynamics as accurate binding free-energy method. *Proceedings of the National Academy of Sciences* **110**, 6358–6363 (2013).

10. Raniolo, S. & Limongelli, V. Ligand binding free-energy calculations with funnel metadynamics. *Nat. Protoc.* **15**, 2837–2866 (2020).

11. Diederichs, K. & Karplus, P. A. Improved R-factors for diffraction data analysis in macromolecular crystallography. *Nature Structural Biology 1997 4:4* **4**, 269–275 (1997).

12. Karplus, P. A. & Diederichs, K. Linking crystallographic model and data quality. *Science (1979).* **336**, 1030–1033 (2012).

13. Adams, P. D. *et al.* PHENIX: a comprehensive Python-based system for macromolecular structure solution. *Acta Crystallogr. D Biol. Crystallogr.* **66**, 213–221 (2010).

14. Williams, C. J. *et al.* MolProbity: More and better reference data for improved all-atom structure validation. *Protein Science* **27**, 293–315 (2018).

15. Krissinel, E. & Henrick, K. Secondary-structure matching (SSM), a new tool for fast protein structure alignment in three dimensions. *Acta Crystallogr. D Biol. Crystallogr.* **60**, 2256–2268 (2004).

16. Abramson, J. *et al.* Accurate structure prediction of biomolecular interactions with AlphaFold 3. *Nature 2024 630:8016* **630**, 493–500 (2024).

17. Boitreaud, J. *et al.* Chai-1: Decoding the molecular interactions of life. Preprint at https://doi.org/10.1101/2024.10.10.615955 (2024).
